# Chromosome organization by a conserved condensin-ParB system in the actinobacterium *Corynebacterium glutamicum*

**DOI:** 10.1101/649749

**Authors:** Kati Böhm, Giacomo Giacomelli, Andreas Schmidt, Axel Imhof, Romain Koszul, Martial Marbouty, Marc Bramkamp

**Affiliations:** Ludwig-Maximilians-Universität München, Fakultät Biologie, Großhaderner Straße 2-4, 82152 Planegg-Martinsried, Germany; Laboratoire Régulation Spatiale des Génomes, Institut Pasteur, UMR 3525, CNRS, 75015 Paris, France; Ludwig-Maximilians-Universität München, Zentrallabor für Proteinanalytik, Biomedizinisches Centrum München, 82152 Planegg-Martinsried, Germany

**Keywords:** ParB, *parS*, SMC, MksB, condensin, actinobacteria, chromosome conformation capture, chromatin immunoprecipitation, super-resolution microscopy, DNA segregation

## Abstract

Higher-order chromosome folding and segregation is tightly regulated in all domains of life. In bacteria, details on nucleoid organization regulatory mechanisms and function remains poorly characterized, especially in non-model species. Here, we investigate the role of DNA partitioning protein ParB and condensin complexes, two key players in bacterial chromosome structuring, in the actinobacterium *Corynebacterium glutamicum*. Chromosome conformation capture reveals SMC-mediated long-range interactions around ten centromere-like *parS* sites clustered at the replication origin (*oriC*). At least one *oriC*-proximal *parS* site is necessary for a reliable chromosome segregation. Using a combination of chromatin immunoprecipitation and photoactivated single molecule localization microscopy evidences the formation of distinct ParB-nucleoprotein subclusters in dependence of *parS* numbers. We further identified and functionally characterized two condensin paralogs. Whereas SMC/ScpAB complexes are loaded via ParB at *parS* sites mediating chromosomal inter-arm contacts like in *Bacillus subtilis*, the MukBEF-like SMC complex MksBEFG does not contribute to chromosomal DNA-folding. Rather, the MksBEFG complex is involved in plasmid maintenance and interacts with the polar *oriC*-tethering factor DivIVA. These data complement current models of ParB-SMC/ScpAB crosstalk, while showing that some condensin complexes evolved functions uncoupled from chromosome folding.

## Introduction

Each organism must complete genome replication and separation in the course of one cell cycle prior to cell division in concert with transcriptional processes. To this end, chromosomes are highly organized structures in terms of segregation and overall folding patterns ^1^. The functional organization of bacterial genomes, structured into the nucleoid, has been predominantly investigated in a limited number of model species, e.g. *E. coli*, *V. cholerae*, *B. subtilis* or *C. crescentus*, revealing diverse levels of compaction and segregation strategies^2–4^.

ParABS systems and condensins are two (nearly) ubiquitous bacterial enzyme machineries that contribute to chromosome homeostasis. With a few exceptions amongst γ-proteobacteria, all branches of bacteria and several Archaea harbor *parS* sites that recruit partitioning protein ParB ^5^. The ParABS system contains one or several *parS* sites usually in the vicinity to the chromosomal origin of replication (*oriC*). ParB proteins bind to these sequence-specific motives and form large nucleocomplexes by spreading and 3D-bridging between ParB dimers ^6–9^, resulting in large topological domains encompassing the *oriC*, that have been revealed by Hi-C for *B. subtilis* ^10^. In an alternative model termed nucleation and caging, ParB-nucleation at *parS* is stabilized by dynamic ParB dimer-dimer interactions and weak interactions with non-specific DNA generating a scaffold for locally high ParB concentrations confined around *parS* ^11^. The ParB segregation is driven by a ParA ATPase, which binds nonspecifically to the nucleoid and is released from DNA upon ATP hydrolysis triggered by transient ParB-interactions ^12,13^. In the course of chromosome replication ParB-*oriC* complexes act in combination with ParA as Brownian ratchets along dynamic DNA loci: slow ParA-DNA rebinding rates generate ParA-gradients, which serve as tracks for directed movement of partition complexes away from their sisters ^14–17^. Perturbation of the system by placing *parS* sites at ectopic, *oriC*-distal regions can cause severe DNA segregation phenotypes ^18,19^. To date, only few studies investigated the impact of chromosomal *parS* localization on DNA-segregation and folding ^18–22^.

In addition to ParABS systems, most bacteria harbor condensin complexes, members of the structural maintenance of chromosomes (SMC) family of proteins found in all kingdoms of life ^23^. In standard model organisms, condensins are equally essential for faithful chromosome segregation by compacting DNA into separate nucleoids ^24–26^. The SMC/ScpAB (structural maintenance of chromosomes) complex is well-studied in *B. subtilis*, where it consists of two large SMC subunits and the kleisin ScpA associated with dimeric accessory protein ScpB that assemble into a ring-like structure ^27^. A recent study suggests progressive extrusion of condensin-encircled DNA loops upon conformational changes in the SMC subunit, which leads to a gradual size increase of trapped DNA molecules ^28^. The active process(es) driving DNA extrusion ^29,30^ allow(s) for translocation along the chromosome with velocities of around 50 Kb/ min ^31^, and depend(s) on the ATPase activity of SMC ^32^. To be loaded on *parS* sites, SMC/ScpAB complexes necessitate ParB ^20,22,33,34^. They redistribute to distant chromosomal regions, promoting topological changes, and notably the co-alignment of right and left replichores ^10,21,22,31,35^. In sharp contrast with SMC/ScpAB, the *E. coli* condensin MukBEF does not promote the co-alignment of chromosomal arms ^36,37^, but facilitate *cis* structuration by establishing long-range contacts between loci belonging to the same replichores from stochastic chromosomal loci (except for the *ter* region containing the replication terminus) ^37,38^. Despite the importance of condensins in chromosome organization, the role of SMC homologs besides the model species *B. subtilis*, *C. crescentus* and *E. coli* remain largely unexplored. These species all contain a single condensin complex, yet a broad range of bacteria possesses combinations of SMC/ScpAB and MksBEFG (MukB-like SMC), for which functional characterizations are non-existent to date ^39^. Current work in bacteria as well as knowledge from eukaryotic studies convey the general assumption that all SMCs are likely to play role(s) in chromosome organization. In bacteria it is unknown, why some species harbor more than one type of condensin, and whether and how they would work in concert with each other and coordinate with systems such as ParABS.

In this work, we used a combination of high-resolution microscopy and genomic chromosome conformation capture (3C/Hi-C) ^35^ to unveil the global organization of the diploid *C. glutamicum* genome. *C. glutamicum* is a polar growing actinobacterium, whose genome encodes both SMC/ScpAB and MksBEFG. In this species, the two *oriC*s are continuously associated with the polar scaffold protein DivIVA, while newly replicated sister *oriC*s segregate towards division septa via the ParABS system ^40–42^. In contrast to *B. subtilis*, *C. glutamicum* ParAB are by themselves crucially important drivers of reliable nucleoid separation prior to cell division, where ParAB deletions yield in 20 % of anucleoid cells ^43–45^. Here, analyses of chromosomal ParB-binding patterns evince ten redundant *parS* sites, which mediate ParB subcluster formation at *oriC*. A single *parS* site maintains ParB propagation over 32 Kb neighboring regions, and is sufficient to promote the SMC-dependent alignment of the two chromosomal arms. Hi-C also reveal SMC-dependent long-range contacts surrounding *oriC*. On the contrary, we showed that the polarly positioned MksBEFG condensin acts exclusively on plasmid-transmission to daughter cells, without influencing nucleoid architecture.

## Results

### Chromosome segregation is governed by a cluster of ten *oriC*-proximal *parS* sites

Previous studies on *C. glutamicum* chromosome partitioning have revealed two stable ParB-*oriC* clusters at each cell pole, while newly replicated origins are segregated towards a division septa formed at midcell ^40^. In *B. subtilis*, *C. crescentus* and *P. aeruginosa* ParAB-mediated chromosome segregation and folding depends on *parS* sites ^18,19,21^. In *C. glutamicum*, *parS* positions have not been characterized yet. Initially, four to eight *parS* sites were predicted in *Corynebacterineae* ^5^. However, we identified ten *B. subtilis*-like 16 bp consensus sequences in *C. glutamicum* localized in one cluster within a 35 Kb region distant 73 Kb from *oriC* using blast (1% of the 3.21 Mb chromosome; Fig. 1A). Out of the ten *parS* sites, only the one located furthermost from *oriC* (*parS1*) lies within a coding sequence (*trpCF*). All other *parS* sequences (labelled *parS 2-10*) are located in intergenic regions. Degenerated *parS* sequences exhibiting at least three base-pair mismatches were also identified further away from *oriC*, e.g. 5’ of *cg0146* or within the *fusA* and *cg1994* coding region. To test whether these putative *parS* were responsible for the recruitment of ParB, α-mCherry-ChIP analyses were performed with a strain harboring a mCherry-tagged version of the native ParB (note that all mutant strains used in this study derive from clean allelic replacements and have, unless otherwise noted, a wild type-like phenotype). Distinct and very reproducible enrichment signals were obtained at ten *parS* sites close to *oriC* (*parS*1-10 at 3.16 MB) (Fig. 1A), whereas the imperfect *parS* sequences identified with blast clearly failed to recruit ParB. Additional smaller peaks were identified at highly transcribed DNA regions, in particular at ribosomal genes, tRNA gene clusters and at all of the rRNA operons (Fig. 1A). Magnification of the *oriC* region reveals three distinct ParB propagation zones overlapping with *parS*1-4, *parS*5-8 and *parS*9-10, respectively (Fig. 1B). Remarkably, those three regions seem to recruit decreasing amounts of ParB, from *parS1-4* (most enriched) to *parS9-10* (less enriched). Since all *parS* are identical in sequence, differences in ParB-recruitment might result from the number and distance of *parS* sequences in the context of the overall nucleoid folding patterns at the *oriC*-region.

**Figure 1.**
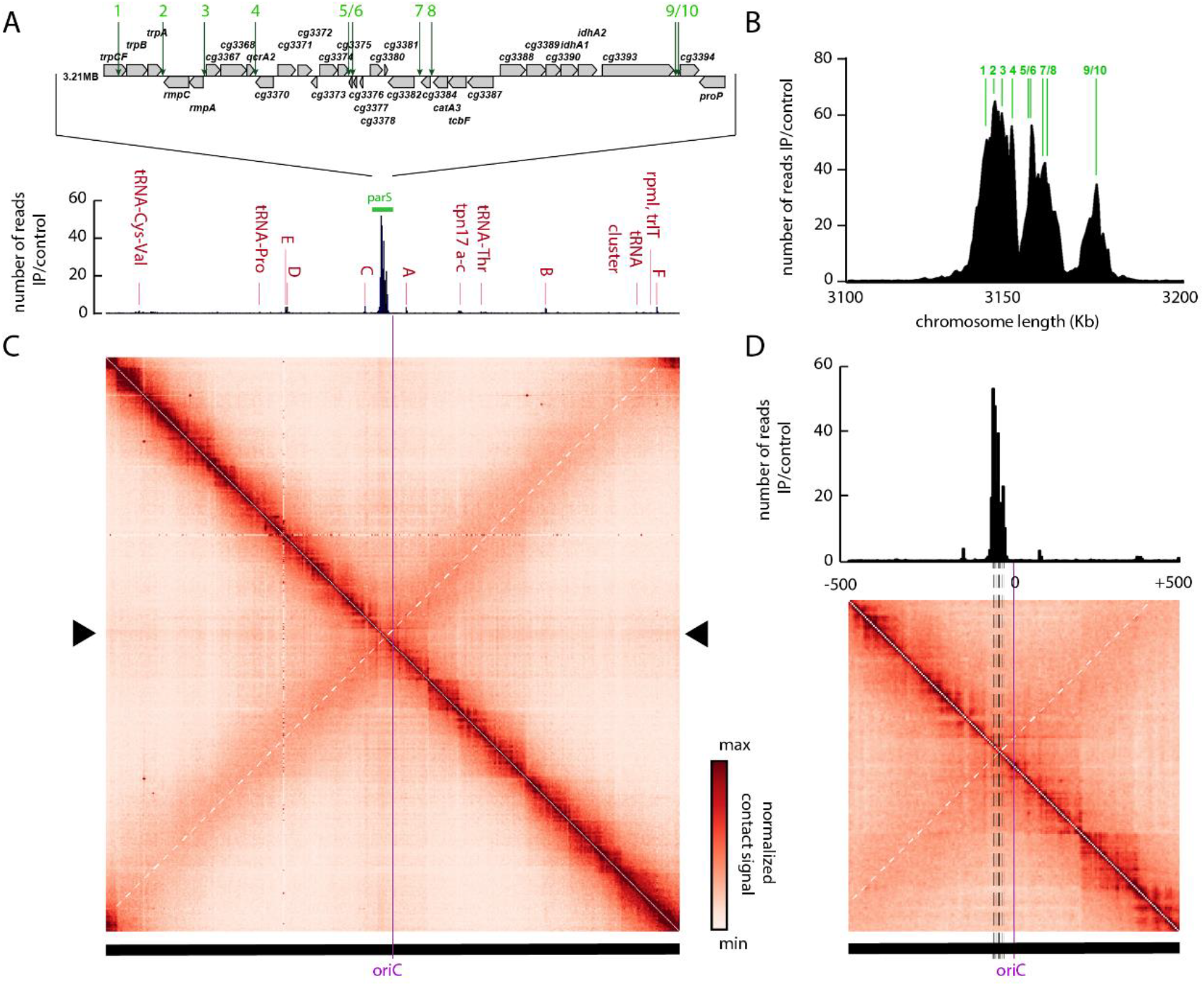
Chromosome organization hub at *oriC* domain in *C. glutamicum*. **A)** Top: Genomic region including ten *parS* sites of *C. glutamicum* with 16 bp consensus sequences. Below: ChIP-seq data on ParB-mCherry DNA binding protein confirm *parS* sites shown above. Exponentially growing *C. glutamicum parB∷parB-mCherry* cells (CBK006) were used for *in vivo* anti-mCherry ChIP-seq experiments. Shown is the ratio of ChIP signal relative to the input (fold enrichment IP/control) in 5 Kb bins in linear scale along the chromosome with an x-axis centered at *oriC*. Red labels indicate minor enrichment signals at highly transcribed regions, like rRNA operons (letters A-F). **B)** ParB-ChIP-seq enrichment encompassing 3.1-3.2 Mb genomic region; *parS* sites 1-10 are indicated (green lines). **C)** Normalized genomic contact map derived from asynchronously grown cells (fast growth µ ≥ 0.6 h^−1^, exponential phase). X-and Y-axes indicate chromosomal coordinates binned in 5 Kb; *oriC*-centered (purple bar). Color scales, indicated beside the contact map, reflect contact frequency between two genomic loci from white to red (rare to frequent contacts). White dashed line on the contact matrix indicate the mean signal of the secondary diagonal and black triangles on the side of the contact matrix indicate the “cross like” signal. **D)** Structural chromosome organization of the *oriC* region. Magnification of contacts within 500 Kb surrounding *oriC*; *oriC* is indicated as a purple line and *parS* sites are indicated by dashed lines. ParB enrichment zones at *parS* are shown above the contact map (ChIP signal relative to the input in 5 Kb bins). White dashed line on the contact matrix indicate the mean signal of the secondary diagonal.

### Higher-order organization of the *C. glutamicum* chromosome

In *B. subtilis*, SMC-mediated chromosome folding initiates at ParB-*parS* clusters surrounding the origin of replication, bridging the two replichores with each other ^10,21^. To characterize whether *C. glutamicum parS* sites play a similar role in the overall organization of the chromosome, we applied a Hi-C like approach ^10,46^ to exponentially growing wild type cells (Material and Methods). The genome-wide contact map, displaying the average contact frequencies between all 5 Kb segments of wild type chromosomes (Fig. 1C) displayed the following 3D features. First, a strong and broad diagonal reflecting frequent local contacts between adjacent loci and observed in all Hi-C experiments. Second, chromatin interaction domains (CIDs), i.e. regions making increased contact frequencies within themselves and previously described in *C. crescentus* and other species ^10,21,35,37,47^, (Fig. 1C, S1) (11 domains detected at a 200 Kb resolution). Third, a secondary diagonal perpendicular to the main one and extending from the 35 Kb *parS* cluster (Fig. 1D – white dashed line) from the *ori* region to the terminus. This structure shows that the two replichores are bridged over their entire length, similarly to *B. subtilis* and, to some extent, *C. crescentus* ^10,21,35^. Interestingly, this secondary diagonal also displays discrete long-range contact enrichments (Fig 1C), which may reflect bridging of the two chromosomal arms at specific locations. Finally, the contact map also displays a faint, cross-shaped signal corresponding to contacts between the *ori* region and the rest of the chromosome (Fig. 1C – dark triangle on the sides of the contact matrix), a feature never described before. These contacts might represent a replication signal reflecting the translocation of the ParB-*oriC* complex along the nucleoid during segregation when *oriCs* reposition at midcell. This signal is also maximal at the *parS* cluster and not at *oriC* locus. An observation that reinforce the fact that the *parS* cluster is at the tip of *Corynebacterium* chromosome fold and is one of the main actor of chromosome segregation.

### A single *parS* site is sufficient to maintain a wild type ParB binding region and chromosome architecture

Since all *parS* sites are in close proximity on the *C. glutamicum* chromosome, we tested the importance of ParB-*parS* complex titration for the overall chromosome organization. Cells with chromosomes carrying a single *parS* site grow and divide like wild type cells (Fig. 2A, S2). However, the removal of all 10 *parS* sites resulted in a cell length phenotype (Fig. S2) and 29% DNA-free mini-cells hinting to a nucleoid segregation defect similar to the Δ*parB* phenotype (Fig. 2A, Tab. S1). We further analyzed ParB localization in mutant strains carrying either one or none *parS* sites. Firstly, cellular localization of fluorescent ParB-eYFP foci is similar to wild type, positioning at cell poles and migrating to the newly formed septa (Fig. 2B, S3) ^40^. Interestingly, the combination of a single *parS* site with ParB-eYFP resulted in 7% anucleoid mini-cells (Fig. S2, Tab. S1), reflecting functional constraints of the ParB-eYFP fusion in presence of only one *parS* site. Therefore, the high number of chromosomal *parS* sites likely evolved to improve the robustness of the segregation machinery. ParB ChIP-qPCR signals of locus *parS1* were similar in both wild type and mutant strains (Fig. 2C). ParB spreading around the single *parS* site was characterized through ChIP-seq analysis (Fig. 2D, S4), where ParB binding was maximum within 2 Kb windows on both sides of *parS*, while extending up to 16 Kb on either side. If each *parS* site promotes ParB deposition in comparable ranges to *parS*1, abrupt enrichment drops downstream of *parS4* and *parS8* would have been absent in wild type, suggesting that DNA properties accountable for the regulation of ParB deposition are independent of *parS* distributions along the nucleation zone. We next investigated the role of *parS* sites and ParB in the overall chromosome folding by performing Hi-C in mutants (Fig. 2E, 2F). The absence of ParB or all *parS* sites led to the disappearance of the secondary diagonal. In addition, the cross-shaped pattern resulting from contacts between the ori and the whole chromosome disappears in those mutants, also illustrated by the ratio between wild type and mutant contact maps (Fig. 2F). This result shows that *parS* sites and ParB are two major structural components of chromosome organization and act to recruit downstream factors that fold the chromosome emanating from the *parS* cluster, and bridge the two chromosomal arms together down to the replication terminus region. The contact map of the strain deleted for *parS2-10* but carrying *parS1* maintains a secondary diagonal, showing that a single *parS* site is sufficient to ensure the loading of ParB and the overall folding of the chromosome (Fig. 2E, 2F). However, some differences appeared between wild type contact matrix and the one resulting from the strain harboring only one *parS* site. In this mutant, the large domain surrounding *oriC* appears more defined than in wild type suggesting that one *parS* site is not sufficient to fully restore *Corynebacterium* chromosome folding possibly due to a slower replication and consequently a less dynamic folding (Fig. 2E, 2F). The single *parS* site was repositioned at different genomic regions. Cells harboring an ectopic *parS* site at 9.5°, 90°, 180° or 270° positions were viable (Fig. S2, S3A, B). Unlike in cells harboring *parS*1 at its original position, ParB-*parS* complexes distribute randomly along the longitudinal cell axis in all of these mutants (Fig. S3B), leading to around 25% anucleate cells (Tab. S1). Therefore, none of these *parS*-shifts restores controlled nucleoid segregation. The number of ParB foci nevertheless correlates well with cell length (Fig. S3C), excluding replication initiation deficiencies. ParB-binding to a *parS* sequence positioned at the 90° chromosomal position (locus *cg0904*, strain CBK037) was identified in a 9 Kb range on either side of *parS* (Fig. S3D, S4), approximately half the ParB-propagation distance determined for cells harboring one *parS* at its native locus. We also analyzed mutant CBK037 (*parS* at 90° chromosomal position) using Hi-C (Fig. S5). The resulting contact map presents a “bow shape” motif at the position of the aberrant *parS* sequence demonstrating a recruitment of the ParB protein at this location as well as a local folding of the chromosome. However, this loading appears insufficient to fold the whole chromosome and to anchor the two chromosome arms over their entire length. DNA topology, origin replication and overall chromosomal localization might hereby determine *parS*-distant ParB-DNA interaction. Collectively, these results show a redundancy of *parS* sites, yet their function is restricted to a confined *oriC*-proximal region.

**Figure 2.**
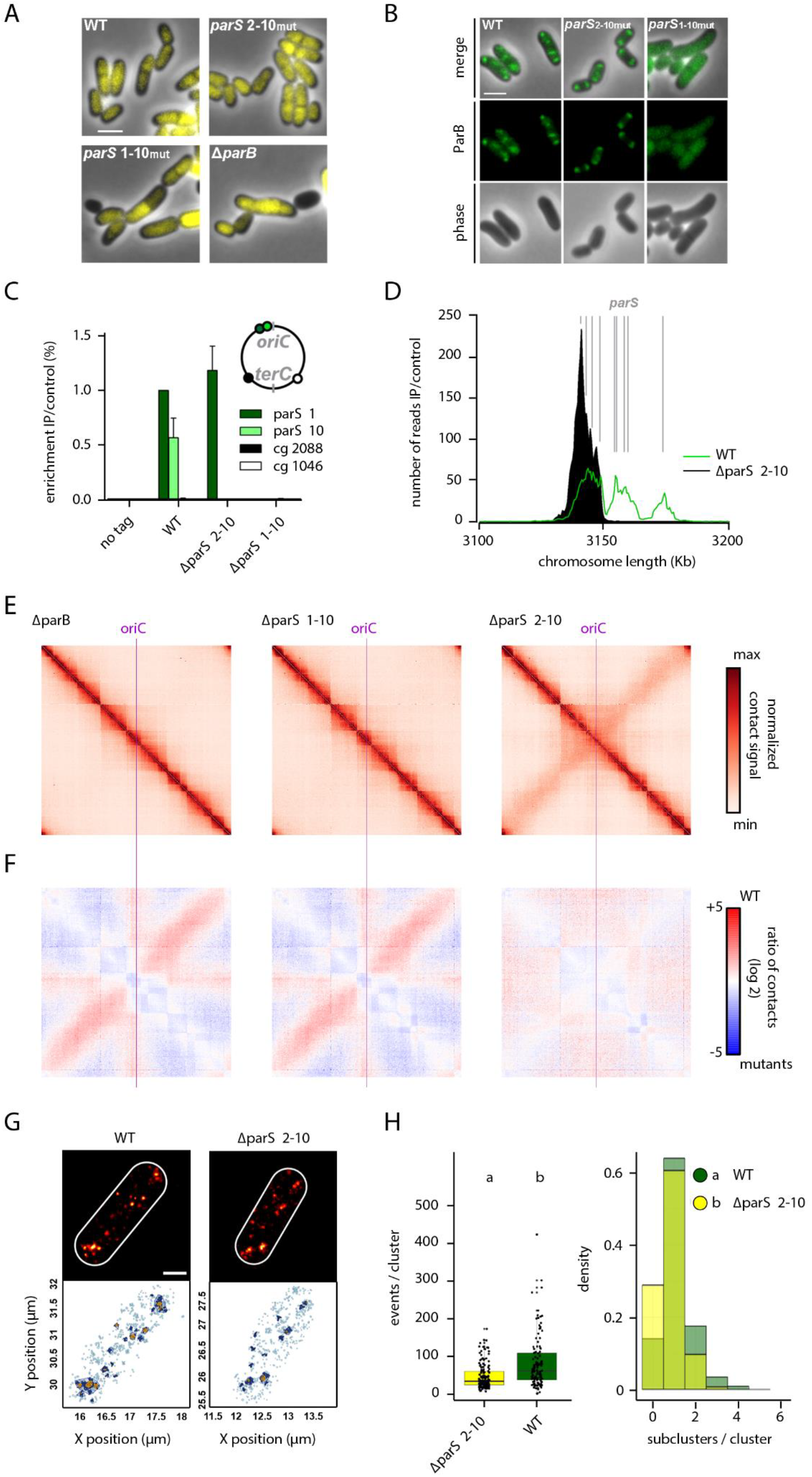
A single *parS* site mediates substantial ParB propagation and wild type-like chromosome folding. **A)** One *parS* site is necessary and sufficient for wild type-like morphology and nucleoid segregation. Shown are phase contrast images of exponentially grown cells harboring either all (WT), one (*parS*_2-10mut_, CBK023) or none (*parS*_1-10mut_, CBK024) *parS* site(s) or lacking *parB* (Δ*parB*, CDC003). DNA is stained with Hoechst (yellow). Scale bar, 2 µm. **B)** One *parS* sequence is sufficient to form wild type-like ParB clusters *in vivo*. Microscopy analysis of *parB∷parB-mCherry* (shown in green) in wild type (CBK006), *parS*_2-10mut_ (CBK027) and *parS*_1-10mut_ backgrounds (CBK028). Absence of *parS* leads to diffuse cellular ParB localizations. Scale bar, 2 µm. **C)** ChIP-qPCR for strains described before, normalized to wild type *parS*1 signal; standard deviations derive from biological triplicates. **D)** ChIP-seq of *C. glutamicum parB∷parB-mCherry parS*_2-10mut_ (black) at a 3.1-3.2 Mb chromosomal range. Wild type-like propagation (green) of ParB protein around *parS1-4*; 0.5 Kb bin size. Location of *parS* sites present in wild type or mutant sequences are indicated (gray lines). **E)** Normalized contact maps of Δ*parB*, *parS*_1-10mut_ and *parS*_2-10mut_ mutants centered at *oriC* (CDC003, CBK024, CBK023). Color codes as in Fig. 1 were applied. **F)** Differential maps correspond to the log2 of the ratio (wild type norm/ mutant norm); color scales indicate contact enrichment in mutant (blue) or wild type (red) (white indicates no differences between the two conditions). **G)** Single-molecule localization microscopy of representative wild type and *parS*_2-10mut_ cells (CBK029, CBK031). Top: Gaussian rendering of ParB-PAmCherry signals (0.71 PSF, 1 px = 10 nm), below: color-coded representation of ParB-PAmCherry events within corresponding cells ^48^; all events (light blue), macroclusters (dark blue) and subclusters (yellow) are indicated. Scale bar, 0.5 µm. For detailed parameters see Material and Methods and Fig. S6. **H)** Comparison of ParB-PAmCherry cluster properties in strains described above. Top: Events per macrocluster, medians are indicated as solid lines and whiskers mark 1.5 IQR (Inter Quartile Ranges). Note that only the two biggest clusters per cell were taken into account for analyses (clusters_wild type_: n = 130, clusters_*parS*2-10_: n = 143). Below: Subcluster numbers per macrocluster shown as overlay bar chart for both strains. Both subcluster numbers per macrocluster (Kruskal-Wallis Rank Sum Test: chi-squared = 12.284, df = 1, p<0.05) and macroclusters size (Kruskal-Wallis Rank Sum Test: chi-squared = 27.582, df = 1, p<0.05) differ significantly between both strains.

### ParB subclusters identified by PALM reflect protein enrichment at *parS* sites

To directly characterize *oriC* domain compaction via ParB, we applied photoactivated localization microscopy (PALM) to visualize individual ParB-PAmCherry molecules with nanometer resolution. PALM revealed distinct ParB-dense regions at cell poles and quarter positions regions, similar to foci observed via diffraction limited epifluorescence microscopy (Fig. 2G). These ParB-enriched regions (macro-clusters) display heterogeneous densities, with a variable number of higher density zones within sub-clusters. Macro- and sub-clusters have been identified via the Optics algorithm^48,49^ (see Material and Methods) and analyzed in strains harboring a single, two or all the *parS* sites (Fig. 2G, S6). We define a macrocluster as 32 events being localized within a maximum distance of 50 nm for macroclusters and 35 nm for subclusters. Note that high chromosome numbers promote inter-molecular *oriC*-colocalization in fast-growing cells. For more accurate cluster estimations, PALM analysis was performed using slow-growing cells resulting in significantly fewer ParB macro-clusters per cell (Fig. S6B) ^40^. Since segregation of *oriC* complexes might alter their DNA compaction, we focused on the two largest macroclusters per cell, stably tethered at cell poles. The amount of ParB contained within each macro-cluster in wild type is significantly higher than in cells containing a single *parS* site (Fig. 2H), in agreement with the ParB deposition observed via Chip-seq. A parallel between PALM and Chip-seq can also be drawn with respect to the number of sub-clusters per macro-cluster, with a higher number of sub-clusters in the wild type compared to the single *parS* site (Fig. 2H). These differences were not observed when comparing cells harboring all, or two *parS* sites (*parS*1,10) (Fig. S3, S4, S6). These observations could explain the differences observed between contact matrices of wild type and ∆*parS* 2-10 strains and the higher structuring of the *oriC* domain when only one *parS* site is present. We therefore conclude that the architecture of *C. glutamicum* partition complex is dependent on *parS* and ParB-*parS* nucleoprotein-complexes are visible as individual subclusters.

### *C. glutamicum* harbors two paralogs of condensin complexes

In bacteria, the condensin paralog complexes SMC/ScpAB and, in *E. coli* and other enterobacteria, MukBEF, are key players of chromosome folding ^10,21,35,37^. MksBEF (for MukBEF-like SMC) is another condensin occasionally found in bacteria genomes ^39^, whose role(s) remain(s) obscure. A sequence homology search of the *C. glutamicum* genome pointed at the presence of both SMC/ScpAB and MksBEF. The SMC/kleisin is encoded by genes *cg2265* (*smc*), *cg1611* (*scpA*) and *cg1614* (*scpB*) (Fig. 3A), while the Mks complex is encoded on a widely conserved operon^39^ and comprises genes *cg3103*-*cg3106* (*mksGBEF*) (Fig. 3A), including MksG which has being suggested to act in complex with MksBEF ^39^.

**Figure 3.**
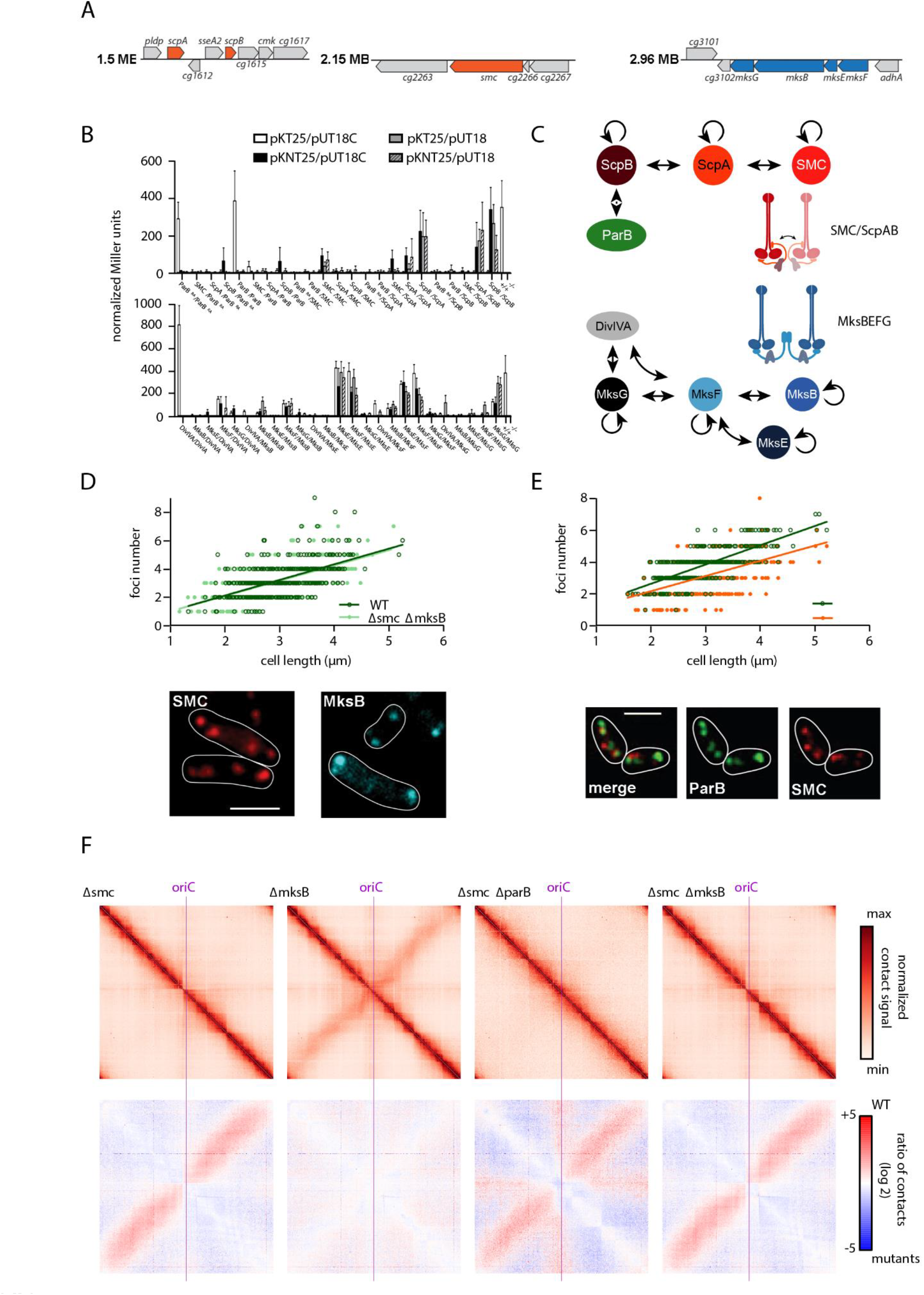
Functional characterization of two SMC-like complexes in *C. glutamicum*. **A)** Sections of the *C. glutamicum* genome map indicating localizations of condensin subunit genes. **B)** Confirmation of protein-protein interactions via bacterial two-hybrid screen. Interactions were quantified by β-galactosidase assays in all combinations of hybrid proteins: C/C-(18C/T25), N/C-(18/T25), C/N-(18C/NT25), and N/N-(18/NT25) terminal fusions of adenylate cyclase fragments, ParB^RA^: ParB mutant R175A. **C)** Illustration of SMC/ScpAB and MksBEFG subunit interactions based on bacterial two-hybrid data, cartoons indicate condensin complex formations. **D)** Top: Dependence of ParB foci numbers on cell length in *C. glutamicum* wild type (WT) and Δ*smc* Δ*mksB* (ΔΔ, CBK011) cells grown in BHI (n>350). Linear regression lines are shown r(WT)=0.57, r(ΔΔ)=0.62; slopes and intercepts are equal (ANCOVA, F(1, 770)=0.059, p >.05; ANCOVA, F(1, 771)=0.60, p <.05). Below: Cellular localization of condensin subunits in *C. glutamicum smc∷smc-mCherry* and *mksB∷mksB-mCherry* cells (CBK012, CBK015). Microscopy images exemplify cellular mCherry fluorescence of SMC (left) and MksB (right); white lines indicate cell outlines. Scale bar, 2 µm. **E)** Top: SMC and ParB foci numbers positively correlate with cell length in double labeled strain *smc∷smc-mCherry parB∷parB-mNeonGreen* (CBK013), r(ParB)=0.74, r(SMC)=0.53; (n>350). Below: Subcellular localization of ParB and SMC is exemplified in representative cells shown in overlays between mNeonGreen and mCherry fluorescence and in separate channels. Scale bar, 2 µm. **F)** Normalized contact maps of Δ*smc*, Δ*mksB*, Δ*parB*/Δ*smc* and Δ*smc*/Δ*mksB* mutants (CDC026, CBK001, CBK002, CBK004), displayed as in Fig. 1. Corresponding differential maps indicating the log of the ratio (wild type norm/ mutant norm) are presented as in Fig. 2.

To characterize condensin complex formation *in vivo*, mass spectrometry of pulldown experiments using SMC and MksB as baits of whole cell lysates were performed. Stability of SMC and MksB fluorescent fusions were confirmed by western blotting (Fig. S7). Kleisin subunit ScpA and ScpB co-precipitated significantly with SMC compared to the negative control, while subunits MksF and MksE, but not MksG, were substantially enriched in the MksB pulldown experiments (Fig. S7). ParB, which mediates SMC-loading onto DNA in *B. subtilis* and *S. pneumonia* ^20,33,34^, was not immuno-precipitated with SMC in any of the experiments. Bacterial two-hybrid analyses confirmed mass spectrometry results, pointing at the formation of SMC/ScpAB and MksBEF complex (Fig. 3B, C). No significant interactions between SMC/ScpAB and ParB were detected, and we observed ScpA-ScpA self-interaction signals well above background. Moreover, MksG connects to the MksBEF complex via interaction with MksF, while MksF and MksG subunits further interact with the *C. glutamicum* polar scaffold protein DivIVA.

### SMC-mediated cohesion of chromosomal arms

We aimed to characterize *C. glutamicum* condensin SMC/ScpAB. Mutation of the SMC/ScpAB complex causes a conditionally lethal phenotype due to chromosome mis-segregation in *B. subtilis* ^25^. In contrast, a *smc* deletion in *C. glutamicum* did not result in growth defects, DNA-segregation defects or aberrant cell length distributions and morphologies compared to the wild type in minimal or complex media (Fig. S8, Tab. S1). Nonetheless, the combination of genetic backgrounds *parB∷parB-eYFP* and Δ*smc* yield a minor fraction of anucleate cells (4-5 %) (Tab. S1), indicating that SMC and ParB function in the same pathway and have a synthetic phenotype. Hence, a functional interaction of SMC and ParB proteins regulating chromosome organization is likely. In order to further determine cellular localization of SMC/ScpAB complexes, a strain harboring a fluorescently tagged version of core subunit SMC was imaged, revealing the formation of SMC clusters along the entire longitudinal axis of the cell (Fig. 3D). Clusters of SMC and ParB investigated in a strain carrying both labelled complexes (*parB∷parB-mNeonGreen smc∷smc-mCherry*) are often proximal but do not necessarily co-localize, while the foci numbers correlate with cell length (Fig. 3E). Up to eight SMC-mCherry foci were counted per cell. On average, cells contained fewer SMC-foci than ParB nucleoprotein complexes (Fig. S7). To further characterize the role of SMC, we generated Hi-C contact maps of the mutant (Fig. 3F). Deletion of *smc* abolishes the secondary diagonal in the maps (Fig. 3F). The combination of *smc* and *parB* mutations mimics a *parB* phenotype, again resulting in the loss of contacts between chromosomal arms and further in the loss of the segregation signal described before (Fig. 3F). Therefore, an interplay of SMC/ScpAB with ParB is responsible for replichore cohesion in *C. glutamicum*, similar to *B. subtilis* and *C. crescentus* each harboring only one condensin complex ^10,21,22,35^. Thus, ParB acts epistatic to SMC.

### ParB-dependent SMC-recruitment to chromosomal loading sites

Since cellular SMC-mCherry signal hinted to distinct agglomeration clusters along the *C. glutamicum* chromosome, we investigated its putative binding sites via ChIP-seq. A small enrichment in SMC deposition was detected at and around the *parS*1-10 cluster (Fig. 4A) that disappears upon *parB* or *parS* deletion (Fig. 4A, S4, S9). In addition, comparably minor enrichment signals are present throughout the chromosome, which partially coincide with genomic loci of high transcriptional activity. Distinct SMC-mCherry foci are less frequent in the absence of ParB or *parS*. (Fig. S9). These findings suggest that ParB promote condensin loading onto DNA at *oriC*-proximal *parS* sites. In addition, ChIP-seq revealed that SMC concentrates at a 13 Kb region upstream *parS*1 (Fig. 4A). SMC enrichment in this region was lost following a partial deletion of this locus and its -reinsertion at another genomic position or following its substitution by a random DNA sequence (Fig. S9). Therefore, the accumulation of SMC at the 13 Kb region in the vicinity of *parS* sites points at roadblocks that trap SMC, rather than specific SMC-binding. This hypothesis is further supported by the study of the contact map of wild type cells (Fig. 1C, 1D, S1, S10). Indeed, the SMC enrichment region is clearly delimited by a strong border on its left (Fig. S1 – Directional Index at 100 Kb resolution and Fig. S10 – red dashed line). In the absence of ParB or SMC (Fig. S10), the strong border observed in HiC maps is shifted towards *parS* sites. Therefore, this border originates from a combination of multiple processes.

**Figure 4.**
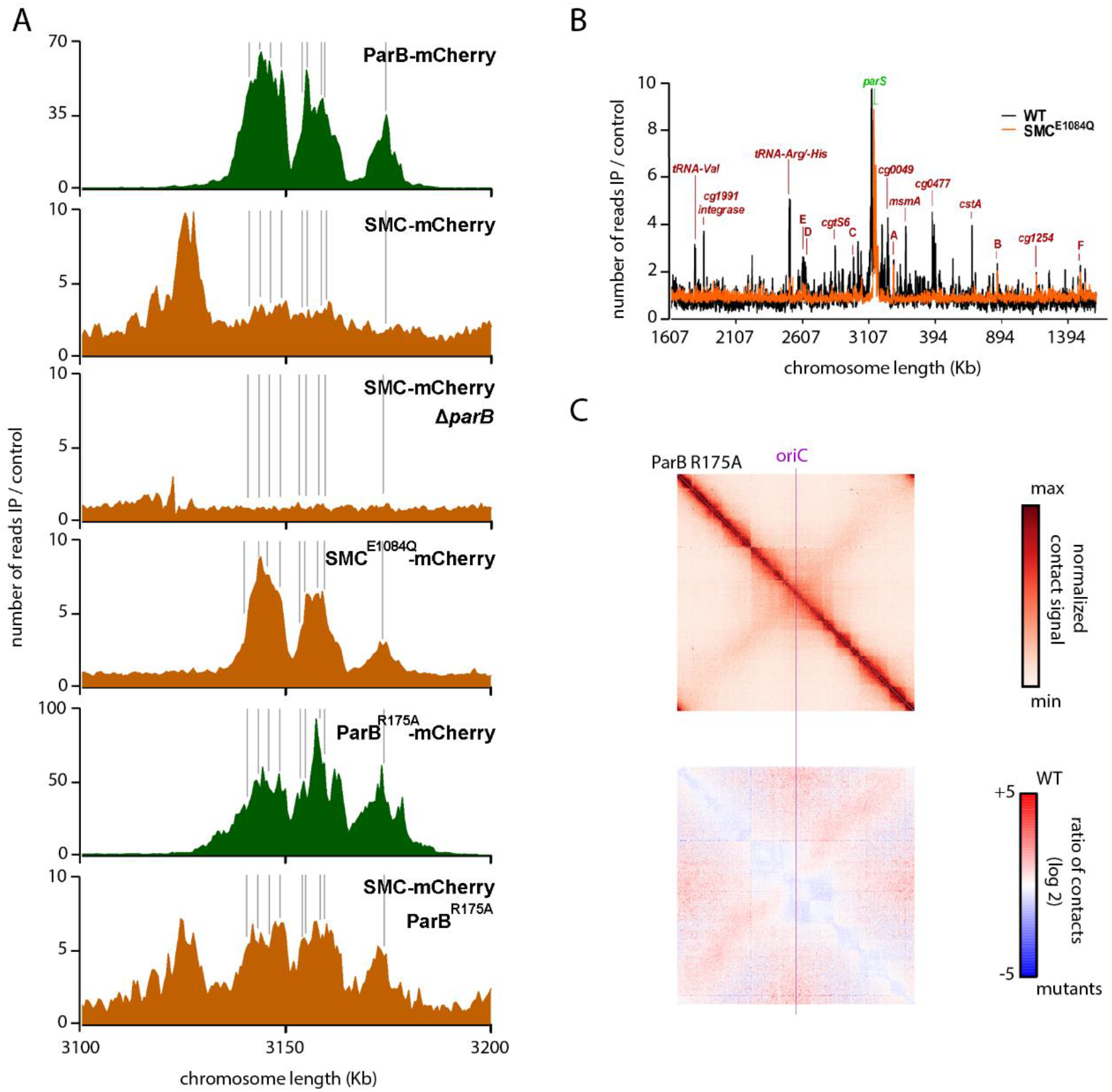
Chromosomal SMC-loading is mediated by ParB at *parS* sites. **A)** SMC-enrichment at *parS* sites (gray) is ParB-dependent. ChIP-seq of ParB-mCherry (green, CBK006, CBK047) and SMC-mCherry (orange, CBK012, CBK014, CBK051, CBK049) in strain backgrounds as indicated. Depicted are chromosomal ranges of 3.1-3.2 Mb, bin size 0.5 Kb. **B)** Whole-genome ChIP-seq data of strains harboring SMC-mCherry wild type (gray, CBK012) or E1084Q mutant (orange, CBK051). SMC enrichment at *parS* sites and at other loci (red letters), in particular tRNA gene clusters and at rRNA genes (A-F) is illustrated in 0.5 Kb bins in linear scale along the chromosome with an x-axis centered at *oriC*. **C)** Normalized contact map of mutant strains *parB∷parB*^R175A^ (CBK047) and the corresponding differential map indicating the log of the ratio (wild type norm/ mutant norm) as in Fig. 2.

SMC is also recruited to *parS* inserted in ectopic positions, e.g. the 90° *parS*-insertion (Fig S4). Indeed, in the absence of SMC (Fig. S5), the bow shape motif is no longer present at the ectopic *parS* site demonstrating that chromosomal arm cohesion is SMC-dependent and that artificial loading of SMC at non-native positions is not sufficient to fold the entire chromosome. We further assayed chromosomal SMC-loading sites by making use of a well-characterized SMC ATP-hydrolysis mutant E1084Q ^32,50–52^. SMC^E1084Q^ mutant strongly accumulates at *parS* sites in *C. glutamicum*, mimicking a ParB-enrichment pattern (Fig. 4A). Decreased ChIP-enrichment signals throughout the rest of the chromosome hint to an impaired SMC-migration along DNA (Fig. 4B). Conclusively, we confirm specific SMC-loading by ParB to an *oriC*-proximal region on the *C. glutamicum* chromosome.

Interestingly, ChIP-analysis of a *C. glutamicum* ParB^R175A^ mutation, which leads to a loss of dimer-dimer interactions in the corresponding *B. subtilis* ParB^R79A^ mutation ^8^, results in increased SMC-binding at ParB^R175A^ propagation zones (Fig. 4A). Changes in *in vitro* dsDNA-binding affinities compared to wild type ParB could not be verified (Fig. S3), neither enhanced binding affinity for SMC/ScpAB by bacterial two-hybrid analyses (Fig. 3B). The mutation results in large fractions of DNA-free cells and growth rates and ParB^R175A^ cluster formation are particularly affected in cells harboring a single *parS* site (Fig. S2, Fig. S3). ChIP-data indicate broadened and less distinct enrichment signals compared to wild type ParB in presence of all or one *parS* sites (Fig. 4A, S3, S4). Therefore, ParB^R175A^ is still capable of building up weak nucleoprotein complexes around *parS* sites. Hi-C data of the corresponding mutant show the same tendency with a conservation of the overall chromosome architecture with the presence of a secondary diagonal and the conservation of the origin domain folding (Fig. 4C, S10). However, the signal ranging from the secondary diagonal is weak compared to the wild type one as shown by the ratio matrix (Fig. 4C). Consequently, SMC-translocation along DNA appears only partially impaired in this mutant (Fig. 4A, S4). The ParB^R175A^ mutation either locks the translocation ability of SMC/ScpAB by a direct interaction or alterations of ParB^R175A^ nucleoprotein complex properties lead to SMC-trapping along DNA-loops at *parS*. Altogether, these analyses confirm that the *C. glutamicum* SMC/ScpAB complex is a *Bacillus*-like condensin that loads and redistributes to distant chromosomal regions via an explicit ParB-crosstalk at *parS*.

### MksB impacts on plasmid maintenance in *C. glutamicum*

To test whether both *C. glutamicum* condensins SMC and MksB are redundant in function, we generated mutants lacking the condensin core subunit Δ*mksB* or both Δ*smc* Δ*mksB*. Similar to Δ*smc*, no growth and morphology phenotypes could be detected for both mutants (Fig. S8, Tab. S1). A triple mutation Δ*parB* Δ*smc* Δ*mksB* did not attenuate the Δ*parB* phenotype, excluding redundancy of condensin functions in chromosome segregation (Fig. S8). Further, *oriC*-ParB foci numbers (Fig. 3D) as well as their spatiotemporal localization (Fig. S8, time-lapse microscopy not shown) remain largely unaffected upon deletion of *smc* and *mksB*. MksB fluorescence was mainly detected at the cell poles (Fig. 3D), further supporting an interaction with the polar protein DivIVA. Moreover, we applied Hi-C to characterize the role of MksB in genome folding in the different mutants (Fig. 3F). In contrast to *smc*, deletion of *mksB* had no effect on chromosome organization, as shown by the nearly white ratio map between the wild type and the mutant (Fig. 3F). Moreover, ∆*smc* and ∆*smc*∆*mksB* contact maps were nearly identical (Fig. 3F), showing that MksB and SMC are most likely not involved in the same process(es). ChIP-seq of MksB failed to detect specific loading sites along the *C. glutamicum* chromosome (Fig. S11), supporting the hypothesis that MksB, unlike other bacterial condensins studied so far, plays no direct or indirect role in *C. glutamicum* chromosomes organization. Therefore, we analyzed its impact on the maintenance of extrachromosomal DNA. The MksBEFG complex appears involved in plasmid maintenance, as shown by the qPCR copy number analysis of two low-copy number (pBHK18 and pWK0) and two high-copy number (pJC1 and pEK0) plasmids. In Δ*mksB* mutants both low-copy number plasmids were enriched up to ten-fold compared to wild type, when grown in the absence of selection marker (Fig. S11). On the contrary, the amount of high copy number vectors per cell was hardly affected. We confirmed these findings by plasmid extractions from *C. glutamicum* cells lacking MksB that yielded exceptionally large quantities of pBHK18 and pWK0, turning them into high copy number plasmids under these conditions (Fig. S11). By contrast, amounts of pJC1 and pEK0 did not differ notably compared to control strains. These analyses show a MksB-dependent decrease in plasmid level, specifically of low copy number plasmids.

Together, our data show that the two condensins in *C. glutamicum* evolved very different functions: whereas SMC/ScpAB act with ParB to promote replichore pairing and origin domain organization, MksBEFG does not organize chromosome architecture and seems involved in plasmid maintenance through a mechanism that remains to be characterized.

## Discussion

Condensins are widely conserved enzyme machineries, which have been implicated in chromosome organization of pro- and eukaryotes ^53^. For long it was considered that bacterial genomes encode one condensin complex that would either be of the Smc/ScpAB type as found in *B. subtilis* and *C. crescentus* or the MukBEF complex encoded in *E. coli* and related proteobacteria ^23^. However, recent reports suggested the existence of two or even multiple condensin systems in a single species ^39^. A prominent example is *P. aeruginosa*, where the MksBEF complex was suggested to act in chromosome organization due to a synthetic DNA segregation phenotype in combination with SMC/ScpAB ^39^. Yet, the underlying mechanisms and the precise function of these two condesnin systems remained largely untested. We report here that the Gram positive actinobacterium *C. glutamicum* also contains SMC/ScpAB and the Muk-like MksBEFG complexes. We set out to address the individual functions of the two condensin systems. Surprisingly our data provide clear evidence that the class of MksBEFG proteins do not work as chromosomal interactors, thus the function of bacterial condensins in promoting DNA segregation to daughter cells is not generally conserved. A recent bioinformatics study predicted a role for MksBEFG complexes (termed Wadjet system) in plasmid-related defense, where heterologous complex expression conveyed protection against the uptake of a high copy number plasmid ^54^. However, function of MksBEFG in its native host had not been addressed before. We could show that the Mks system is indeed involved in the control of plasmid copy numbers and that there is no involvement of this system in chromosome organization. This is a fascinating similarity to specific eukaryotic condensins such as Rad50, being the closest eukaryotic relative to MukB/MksB ^23,39^. It was recently shown that Rad50-CARD9 complexes sense foreign cytoplasmic DNA in mammalian cells. This includes antiviral effects of Rad50 that are important for innate immune responses. Thus, it takes no wonder that several DNA-viruses have evolved strategies to inhibit Rad50 signaling ^55^. Also the more distantly related eukaryotic SMC5/6 complex had been shown to act in a defense mechanism against circular hepatitis B virus DNA, resembling the specific effect of prokaryotic MksBEFG on plasmids ^56^. Together, our data lend support to the notion that condensins function in innate immunity is an ancient mechanism and condensins complexes that diverged early in evolution have specialized functions beyond chromosome organization. However notably, we provide evidence that the MksBEFG complex is the only known condensin amongst pro- and eukaryotes that exclusively targets non-chromosomal DNA. For MksBEF systems it has been proposed that a fourth subunit, MksG is important for function in plasmid maintenance ^54^. We could verify that MksG is part of the MksBEF complex of *C. glutamicum*. The direct interaction of a Mks complex with a polar scaffold protein like the *C. glutamicum* DivIVA has not been described before, but may be very well in accord of its function in plasmid control. Polar plasmid localization has been described in other systems before ^57^. A challenging question for the future will be to determine how the MksBEFG system can identify foreign DNA and how it is loaded onto the plasmids. In vitro analysis of the related MukB has revealed that the protein preferentially binds to single stranded DNA, although also autonomous loading on double stranded DNA was observed ^58^. Thus, one might speculate that MksB is loaded onto plasmid DNA once replication is initiated.

We further describe here that the second condensin SMC/ScpAB is indeed the major player of replichore cohesion and chromosome organization in *C. glutamicum*. Like in *B. subtilis*, SMC is loaded onto the chromosome by a ParB/*parS* loading complex before spreading to the entire chromosome. The mild DNA-partitioning defects of a *smc* deletion in combination with a ParB-eYFP modification (Tab. S1) strongly suggest a supportive role of SMC/ScpAB in the process of nucleoid separation, yet the *smc* phenotype appears to be entirely compensated by ParB. Therefore, our data demonstrate that the conserved role for SMC in chromosome organization ^10,20–22,34,35^ is also maintained in presence of a second condensin complex. Moreover, bacterial two-hybrid analyses of SMC/ScpAB subunits evidence a self-interaction of *C. glutamicum* kleisin ScpA (Fig. 3), that has not been described in other organisms before. Based on this result, we speculate that *C. glutamicum* SMC/ScpAB might form dimers via kleisin subunits similar to *E. coli* MukBEF complex ^59,60^. These data point to a handcuffing model, where two SMC/ScpAB complexes are physically coupled together and translocate in pairs along the chromosome, similar as suggested for *B. subtilis* ^31^. We further describe a new phenotype for a ParB^R175A^ point mutation in *C. glutamicum*, that blocks SMC-release from its loading site. Building on this, we observe a weak interaction signal of ParB^R175A^ with ScpB in bacterial two-hybrid analyses. Alternatively, SMC/ScpAB remains indirectly entrapped in higher-order ParB^R175A^-nucleocomplexes, which possess altered DNA-folding properties. In either case, this mutant underlines the crosstalk between SMC/ScpAB and ParB nucleoprotein complexes in bacterial nucleoid organization.

Analysis of ParB complexes using 2D PALM reveals ParB-dense regions within clusters that correlate to the number of ParB enrichment zones along adjacent *parS* sites. In line with a current study on ParB cluster-assembly in *V. cholerae* ^61^, we suggest that these subclusters derive from independent nucleation and caging events, which merge into one ParB-macrocomplex per *oriC* in *C. glutamicum*. Presence of a single *parS* site leads to formation of almost globular ParB densities. Using Hi-C approaches, we further show that *parS* sites as well as ParB are major players of chromosome folding in *C. glutamicum* as previously shown in other models ^10,21,22,35^. *C. glutamicum* chromosome adopts a global folding with a strong cohesion between the two chromosomal arms as expected from a bacterium harboring a longitudinal chromosomal organization similar to *B. subtilis* and to a lesser extent *C. crescentus* (Fig. 1). Our analysis also suggests the existence of a chromosomal domain at *parS* sites in *C. glutamicum* as previously observed in *B. subtilis*, but with important differences: *parS* sites in *C. glutamicum* are only found on one side of the *oriC* locus and appeared to be at the edge of the nucleoid structure as observed in *C. crescentus*. A hairpin structure as it was observed in *B. subtilis* is absent in *C. glutamicum*^10^. Contact maps of a strain with an ectopic *parS* site feature a bow-shaped structure reflecting an asymmetry in arm interaction, which has been shown before in *B. subtilis* and *C. crescentus* ^21,22^. Zipping of the chromosome is not complete and the ectopic *parS* site does not reorient the entire chromosome. Therefore, additional factors are involved in chromosome localization that supplement polar ParB-*parS* binding to DivIVA.

Importantly, we describe ParB/*parS*-dependent DNA segregation signals showing that the chromosomal *ori*-*ter* configuration allows for ParAB-driven *oriC*-segregation trailing along the nucleoid. Based on our data we propose the following model shown in Figure 5: Organisms with polarly localized *oriC*s and a longitudinal chromosome organization rely on ParAB for *oriC* segregation, since they can use the DNA-scaffold as a track. By contrast, species with a central replication factory cannot efficiently use ParAB. *B. subtilis* is an exception, since here a longitudinal chromosome orientation is present during sporulation and hence, *parAB* (*spo0J*/*soj*) phenotypes are only obvious during spore formation. Consequently, SMC/ScpAB-mediated replichore cohesion is generally dispensable for *oriC* segregation in bacteria with a strict longitudinal chromosome arrangement that allows for efficient ParABS-driven chromosome partitioning.

**Figure 5.**
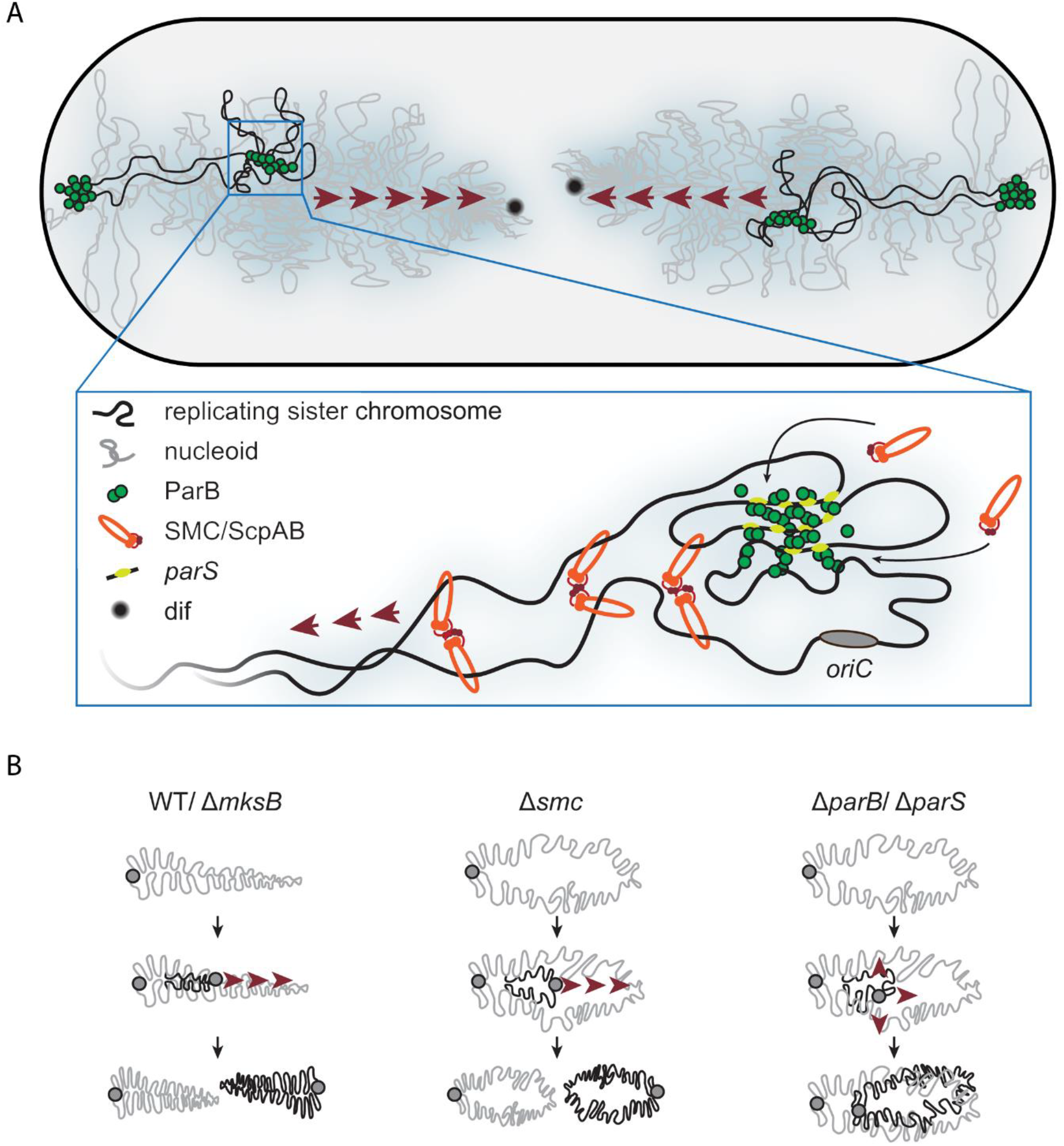
Model of ParB- and SMC-mediated chromosome organization in *C. glutamicum*. A) Top: Illustration of chromosome segregation in a diploid *C. glutamicum* cell. Newly replicated ParB-*oriC*s complexes translocate from cell poles towards *terC*s at septal positions via a ParABS system, where ParB nucleocomplexes use ParA-bound sister chromosomal loci as transient tethers for translocation across the nucleoid. Below: Condensins SMC/ScpAB are loaded ParB-dependently at *parS* sites and relocate to distant chromosomal regions (red arrows) causing inter-linkage of chromosomal arms; here illustration is based on a two-translocator model. **B)** Cartoon showing mutant phenotypes of overall chromosome folding and segregation compared to wild type (WT). Replication *oriC* (grey mark), chromosome (grey line) and replicating sister DNA (black line) are indicated; red arrows indicate direction of DNA segregation.

## Acknowledgements and Funding

This work was further funded by grants to M.B. from the Deutsche Forschungsgemeinschaft (BR2815/6-1 & 2815/6-2) and the Ministry of Science and Education (BMBF: 031A302 e:Bio-Modul II: 0.6 plus).This research was supported by funding to R.K. from the European Research Council under the Horizon 2020 Program (ERC grant agreement 260822).

We thank our colleagues from R.K. and M.B. labs for discussions, feedback and comments on the manuscripts and the interpretation of the data. We are thankful to Prof. Dr. Stephan Gruber (Department of Fundamental Microbiology, University of Lausanne) for inspiring discussions and for his help with ChIP-seq. The authors are grateful to Dr. Andreas Brachmann (LMU) for his help with the genome sequencing.

## Authors contribution

Conceptualization: Kati Böhm, Martial Marbouty, Romain Koszul, Marc Bramkamp Methodology:

- Strain constructions and analyses, epifluorescence microscopy, ChIP, bacterial two-hybrid screening, *in vitro* protein assays: Kati Böhm
- Chromosome Conformation Capture: Martial Marbouty
- PALM: Giacomo Giacomelli
- Mass spectrometry: Andreas Schmidt, Axel Imhof

Writing manuscript: Kati Böhm, Martial Marbouty, Andreas Schmidt (Material and Methods), Giacomo Giacomelli (Material and Methods), Romain Koszul, Marc Bramkamp

## Materials and Methods

### Bacterial strains, plasmids and oligonucleotides

Primers, plasmids and strains used in this study are listed in Tables S2 and S3. Detailed information on strain construction and growth conditions are provided in the **Supplementary Information**.

### Plasmid extraction from *C. glutamicum* cells

*C. glutamicum* cells were grown in 10 ml BHI medium to exponential growth phases in presence of selection antibiotic, following incubation with 20 mg/ml lysozyme in P1 buffer (NucleoSpin^®^ Plasmid Kit, Macherey-Nagel) overnight at 30 °C. Subsequently, plasmids were extracted by using the plasmid kit according to manufacturer’s instruction.

### Protein identification via immunoprecipitation and mass spectrometry

Immunoprecipitation of SMC and MksB interaction partners was performed with strains CBK012 and CBK015, further including strain CBK052 as negative control. Lysate of exponentially grown cells was used for immunoprecipitation via magnetic RFP-Trap® agarose beads. For proteomic analysis samples were further processed and analyzed by liquid chromatography tandem mass spectrometry (LC-MS/MS) to identify and quantify proteins in all samples. A detailed description of immunoprecipitation and proteomic analysis is provided in the **Supplementary Information**.

### Bacterial two-hybrid screening

Protein interactions obtained by mass spectrometry were confirmed via bacterial two-hybrid assays ^62^, using compatible vectors expressing adenylate cyclase subunits T25 and T18 (pKT25/ pKNT25 and pUT18/ pUT18C). *E. coli* BTH101 co-transformed with respective vectors were plated on indicator medium LB/ X-Gal (5-bromo-4-chloro-3-indolyl-β-D-galactopyranoside, 40 μg/ml) supplemented with IPTG (0.5 mM) and antibiotics kanamycin (50 μg/ml), carbenicillin (100 μg/ml) and streptomycin (100 μg/ml) and incubated at 30°C for 24 h. Interacting hybrid proteins were identified by blue-white screening (not shown) and β-galactosidase assays in a 96 well plate format as previously described ^63^. Cotransformants harboring empty plasmids or pUT18C-zip/ pKT25-zip plasmids served as positive and negative controls. Miller units of negative controls served as reference and were set to zero; Miller units of any other sample were normalized accordingly. All C- and N-terminal combinations of hybrid proteins were assayed and positive signals were confirmed through at least three replicates.

### Fluorescence microscopy

Microscopy images were acquired on an Axio-Imager M1 fluorescence microscope (Carl Zeiss) and a Delta Vision Elite microscope (GE Healthcare, Applied Precision). Experimental procedures and technical configurations are specified in the **Supplementary Information**. Data analysis was performed via FIJI and MicrobeJ software ^64,65^.

### Chromatin Immunoprecipitation (ChIP) combined with sequencing

A detailed description of this method is available in the **Supplementary Information**. Briefly, cells were crosslinked (1 % formaldehyde) for 30 min at room temperature and lysed. DNA was sheared by sonication, incubated with α-mCherry antibody for 2 h at 4 °C, washed subsequently and crosslinks were reverted at 65 °C o. N. DNA purification was followed by library preparation and sequencing using an Illumina MiSeq system. Reads were aligned to the *C. glutamicum* ATCC 13032 genome sequence (GeneBankID: BX927147.1). Further data analysis was performed using online tools ^66^.

### Real-time PCR

DNA amplification was performed using a 2x qPCR Mastermix (KAPA SYBR®FAST, Peqlab) according to manufacturer’s instruction, where reaction volumes of 10 µl contained 200 nM oligonucleotides and 4 μl of diluted DNA, respectively. Samples were measured in technical duplicates via an iQ5 multicolor real-time PCR detection system (Bio-Rad) and CT-values were determined via the Bio Rad-IQ™5 software version 2.1. Primer efficiencies were estimated by calibration dilution curves and slope calculation ^67^; data were analyzed via the 2^−∆CT^ method ^68^ accounting for dilution factors and sample volumes used for DNA purification. qPCR data of ChIP samples were normalized according to the ParB-mCherry2 signal obtained at locus *parS1* in the wild type background, serving as reference in each experiment.

### Electrophoretic mobility shift assay

DNA-ParB binding was assayed using purified protein (for protein purification, see **Supplementary Information**) and double-stranded DNA fragments of approximately 1100 bp length with or without two *parS* sites. Fragments were generated by PCRs of a *C. glutamicum* genomic locus surrounding *parS9* and *parS10* using primer pairs parS9mut-HindIII-up-F/ parS10mut-EcoRI-D-R. ParB concentrations of 0.05 to 25 µM were incubated with 100 ng DNA for 30 min at 30°C, following sample separation in native gels (3-12 % polyacrylamide, ServaGel™). DNA was stained using SYBR® Green I (Invitrogen).

### Photoactivated localization microscopy (PALM)

*c. glutamicum* cells were fixed with 3 % of formaldehyde prior to super resolution imaging using a Zeiss ELYRA P.1 microscope with laser lines HR diode 50 mW 405 nm and HR DPSS 200 mW 561 nm and an Andor EM-CCD iXon DU 897 camera. Cellular ParB-PAmCherry signals were further analyzed using Fiji software^64^ and identification of distinct protein clusters was carried out by applying the OPTICS algorithm in R ^48,49^. A detailed description of sample preparation, parameters for detection of ParB-PAmCherry signals, PALM image calculation and analysis is available in the **Supplementary Information**.

### Chromosome Conformation Capture Libraries

Chromosome Conformation Capture (3C/Hi-C) libraries were generated as previously described by Val *et al*. ^47^ with minor changes, as specified in the **Supplementary Information**.

### Contact map generation

Contact maps were generated as previously described ^37^. Reads were aligned independently (forward and reverse) using Bowtie 2 in local and very sensitive mode and were assigned to a restriction fragment. Non-informative events (self-circularized DNA fragments, or uncut fragments) were discarded by taking into account the pair-reads relative directions and the distribution of the different configurations as described in Cournac *et al* ^69^. We then bin the genomes into regular units of 5 Kb to generate contact maps and normalized them using the sequential component normalization procedure (SCN) ^69^.

### Contact map comparison

Ratio between contact maps was computed for each point of the map by dividing the amount of normalized contacts in one condition by the amount of normalized contacts in the other condition and by plotting the Log2 of the ratio. The color code reflects a decrease or increase of contacts in one condition compared to the other (blue or red signal, respectively). No change is represented by a white signal. Ratio maps of replicates are illustrated in Figure S12.

### Identification of domains frontiers using directional index

To quantify the degree of directional preference, we applied on correlation matrices the same procedure as in Marbouty, et al. ^10^. For each 5 Kb bin, we extracted the vector of interactions from the correlation matrix between the studied bin and bins at regular 5 Kb intervals, up to 250 Kb in left and right directions. The two vectors were then compared with a paired t•test to assess their statistical significant difference (p=0.05). The directional preferences for the bin along the chromosome are represented as a bar plot with positive and negative t•values shown as red and green bars.

### Statistical analysis

Correlation coefficients, linear regressions and analyses of nearest neighbor distance distributions were calculated using Excel, Graph Pad Prism (GraphPad Software) and R^49^.

### Data availability

Proteomic data are available via ProteomeXchange with the project identifier PXD008916 ^70^. Reviewer account details: username, password: zyJpq85V

Genome-wide sequencing reads of ChIP-seq and chromosome conformation capture assays generated in this study are available in the Sequence Read Archive (SRA) under accession numbers PRJNA529385 and PRJNA525583.

