## Supplemental Material for "Chromosome organization by a conserved condensin-ParB system in the actinobacterium *Corynebacterium glutamicum*"

### Supplementary Information

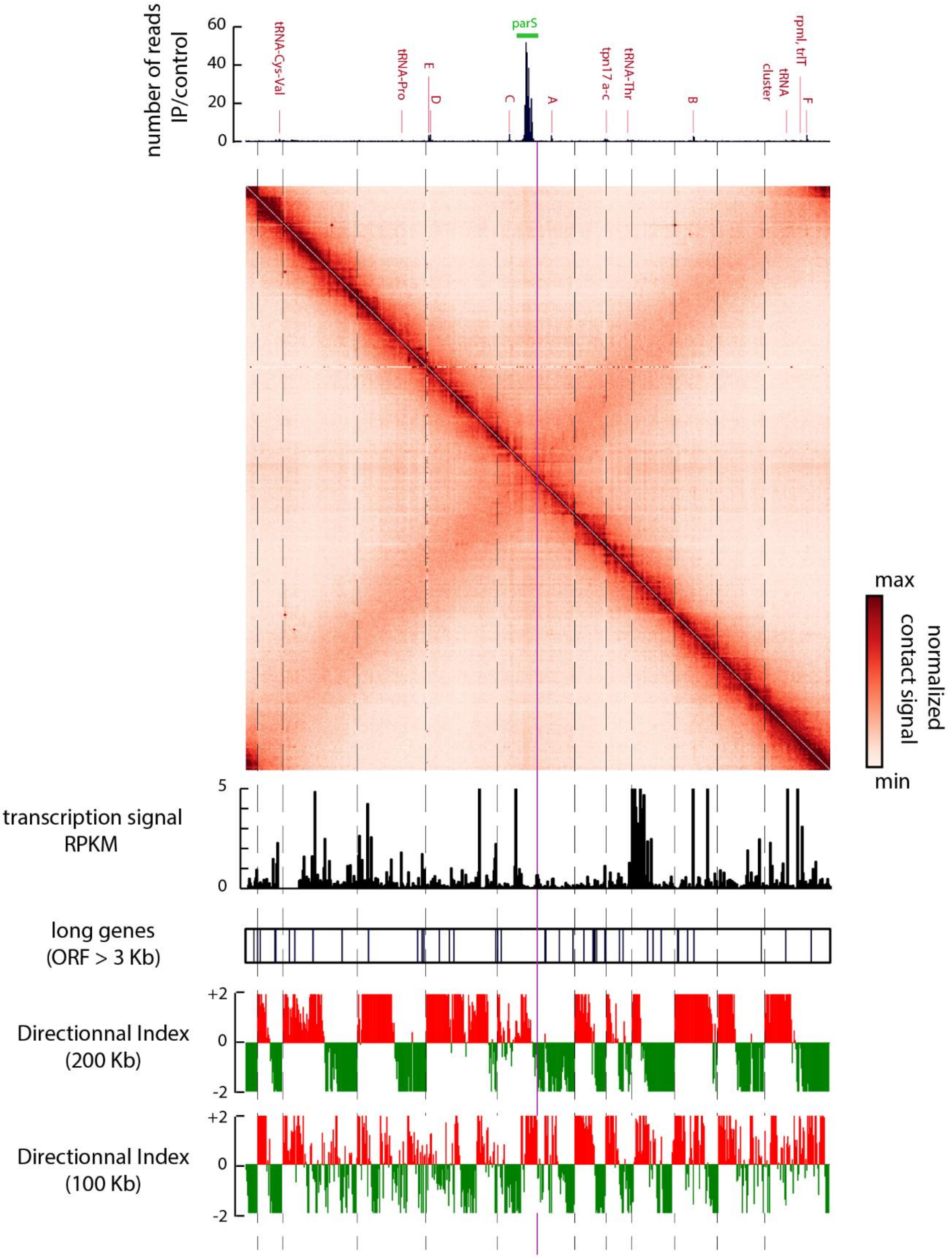


**Figure S1: Chromosome organization and domains analysis in *Corynebacterium glutamicum*.**

From top to bottom: i) Whole genome anti-mCherry ChIP-seq data of strain CKB006 harboring ParB-mCherry. ii) Normalized genomic contact map derived from asynchronously grown cells (fast growth $\mu \geq 0.6\ h^{-1}$, exponential phase). X- and Y-axes indicate chromosomal coordinates binned in 5 Kb; *oriC*-centered as in Fig. 1. Color scales, indicated above the contact map, reflect contact frequency between two genomic loci from white to red (rare to frequent contacts). iii) Transcription signal using 5Kb bins [1]. iv) long genes (black bars) upper than 3 Kb. v) domain signals at 200 Kb resolution (DI analysis) with up- and downstream regions marked in green and red are displayed. vi) domain signals at 100 Kb resolution.

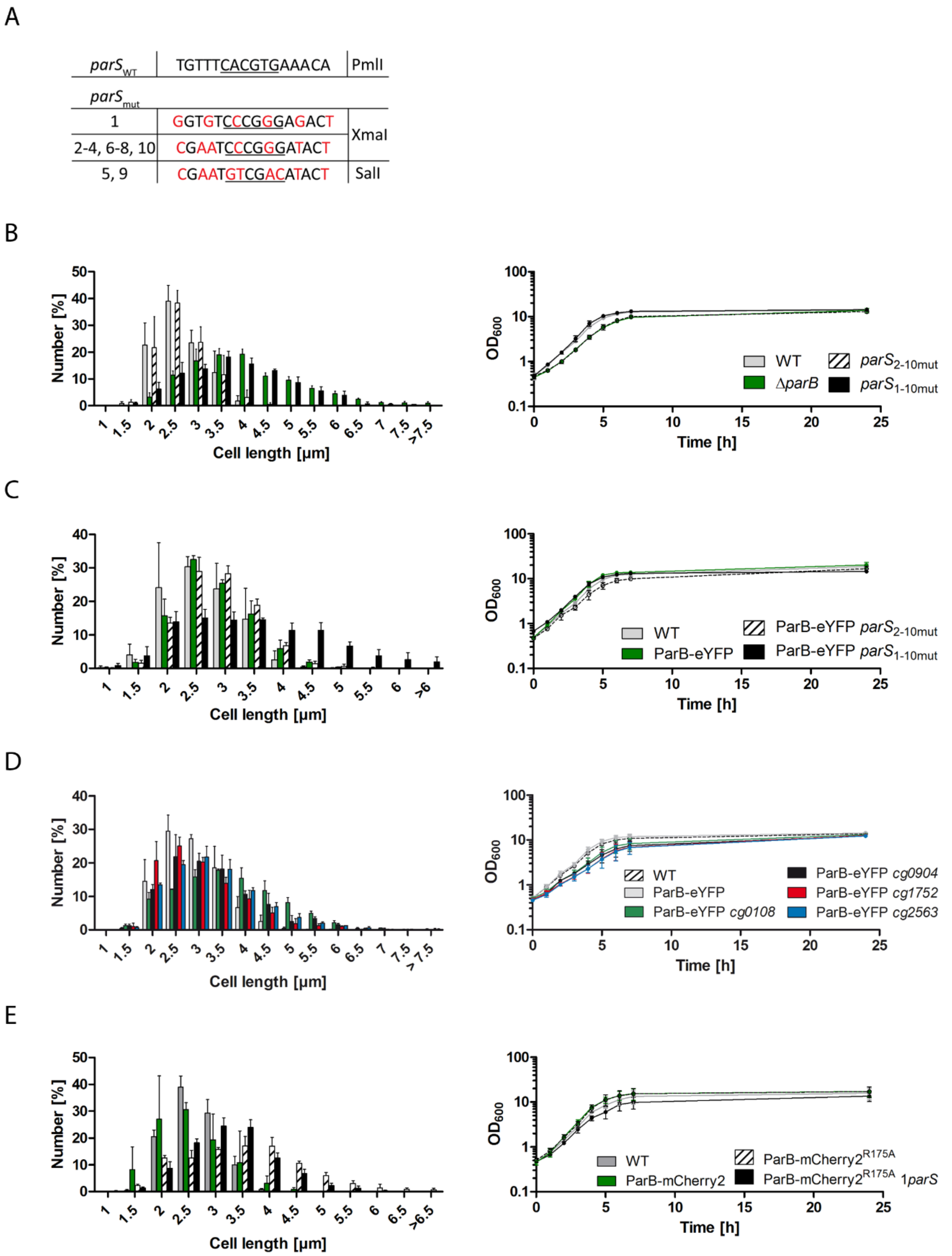


**Figure S2. Mutation and shifting of *parS* sites affects growth and morphology.** **A)** Point mutations render native *parS* sequences nonfunctional. Base pair exchanges (red letters) generate new restriction sites (underscored) for mutation screening as indicated. Notably, point mutations of the intragenic *parS* site are silent. **B), C)** Mutation of *parS* sites mimics growth and morphology of *ΔparB* cells. Comparison of cell length distributions and growth curves of *parS* mutants and *parB* deletion strain grown in BHI. Growth rates derive from triplicates: $\mu_{WT}$=0.65 $h^{-1}$, $\mu_{\Delta parB}$ =0.57 $h^{-1}$ (CDC003), $\mu_{parS2\text{-}10mut}$=0.69 $h^{-1}$ (CBK023), $\mu_{parS1\text{-}10mut}$=0.57 $h^{-1}$ (CBK024), $\mu_{ParB\text{-}eYFP}$=0.65 $h^{-1}$ (CBK007), $\mu_{ParB\text{-}eYFP\ parS2\text{-}10mut}$=0.59 $h^{-1}$ (CBK025), $\mu_{ParB\text{-}eYFP\ parS1\text{-}10mut}$=0.55 $h^{-1}$ (CBK026); error bars indicate standard deviations. **D)** Cell lengths and growth curves determined for mutant cells ParB-eYFP in combination with a single misplaced *parS* site, growth rates: $\mu_{WT}$=0.57 $h^{-1}$, $\mu_{ParB\text{-}eYFP}$=0.61 $h^{-1}$ (CBK007), $\mu_{ParB\text{-}eYFP\ parS\ 3'\ cg0108}$=0.46 $h^{-1}$ (CBK040), $\mu_{ParB\text{-}eYFP\ parS\ 3'\ cg904}$=0.46 $h^{-1}$ (CBK041), $\mu_{ParB\text{-}eYFP\ cg1752}$ =0.40 $h^{-1}$ (CBK042), $\mu_{ParB\text{-}eYFP\ parS\ 3'\ cg2563}$=0.41 $h^{-1}$ (CBK043). **E)** Growth curves and cell length distributions of strain *parB::parB-mCherry* derivatives harboring $ParB^{R175A}$ mutant protein grown in BHI medium are shown; growth rates: $\mu_{WT}$=0.63 $h^{-1}$, $\mu_{ParB\text{-}mCherry}$=0.69 $h^{-1}$ (CBK006), $\mu_{ParBR175A\text{-}mCherry}$=0.68 $h^{-1}$ (CBK047), $\mu_{ParBR175A\text{-}mCherry\ parS2\text{-}10mut}$=0.57 $h^{-1}$ (CBK048).

Figure S3

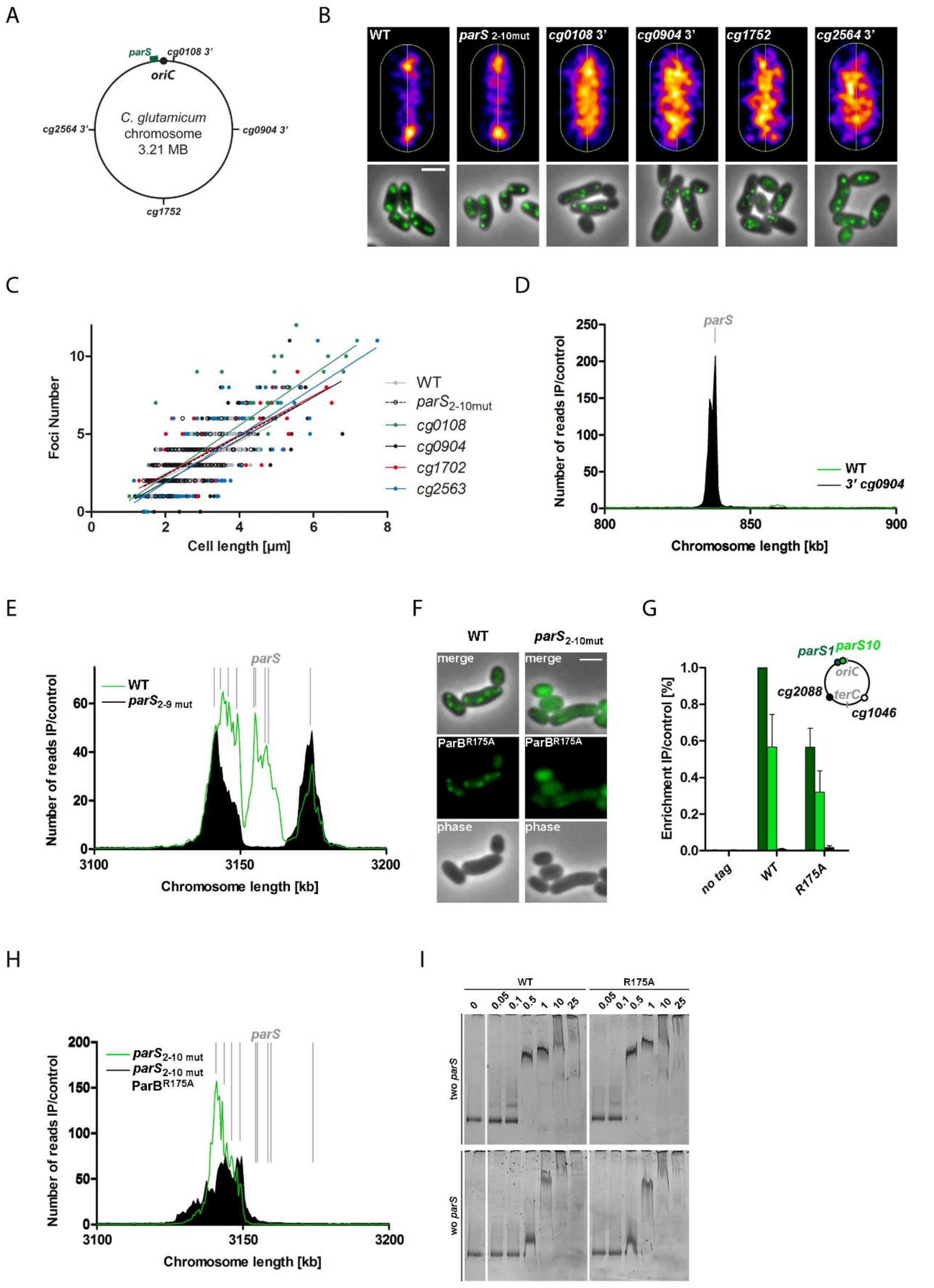


**Figure S3. Confined chromosomal positioning of *parS* impacts on its function.**

**A)** Scheme of chromosomal *parS* insertions, native *parS* cluster shown in green, *parS* shifted to intergenic regions 3' of *cg0108*, 3' of *cg0904*, 3' of *cg2563* or *cg1702::parS* (CBK040, CBK041, CBK043, CBK044). **B)** ParB recruitment to *parS* sites at any chromosomal position remains intact. Top: average ParB-*parS* cluster localizations of *parB::parB-eYFP* cells with native *parS* cluster (WT, CBK007) or one single *parS* site (*parS* $_{2\text{-}10mut}$, CBK025) and *parS* sequences located at respective chromosomal loci illustrated via MicrobeJ (4). Below: ParB-eYFP clusters (green) of representative mutant cells harboring *parS* sites at above-mentioned chromosomal positions. Scale bar, 2 µm. **C)** ParB foci number in relation to cell length in *parS*-shifted mutant strains (n>150). Linear regression lines are shown, r(WT)=0.71, r(*parS* $_{2\text{-}10mut}$)=0.74, r(*cg0108*)=0.86, r(*cg0904*)=0.68, r(*cg1752*)=0.75, r(*cg2563*)=0.79. **D)** α-mCherry-ChIP-seq of *C. glutamicum parB::parB-mCherry parS* 3' of *cg0904* (black, CBK042) compared to wild type enrichment signal (green, CBK006) within genomic region 0.8 - 0.9 Mb including mutant *parS* location (gray), bin size 0.5 Kb. **E)** ChIP-seq of cells harboring *parS*1 and 10 (*parS* $_{2\text{-}9mut}$, CBK30), performed and compared to the wild type-signal as in D). **F)** ParB$^{R175A}$-mCherry localization in wild type and *parS*$_{2\text{-}10mut}$ mutant cells (CBK47, CBK048). Shown are phase contrast, mCherry fluorescence and an overlay of both channels. Scale bar, 2 µm. **G)** ParB-mCherry ChIP-qPCR at specified chromosomal markers performed with cells harboring ParB wild type or R175A mutant. Standard deviations derive from biological triplicates; data normalization to wild type *parS*1 signals. **H)** ChIP-seq enrichment as described in D) at region of native *parS* cluster (gray); ParB$^{WT}$- (green, CBK027) or ParB$^{R175A}$-mCherry protein (black, CBK047) in strain background *parS*$_{2\text{-}10mut}$. **I)** Recombinant ParB$^{WT}$ and ParB$^{R175A}$ proteins bind *parS* sites and nonspecific DNA. Electrophoretic mobility shift assay of ParB proteins pre-incubated with 100 ng DNA sized 1084 bp with or without two *parS* sites.

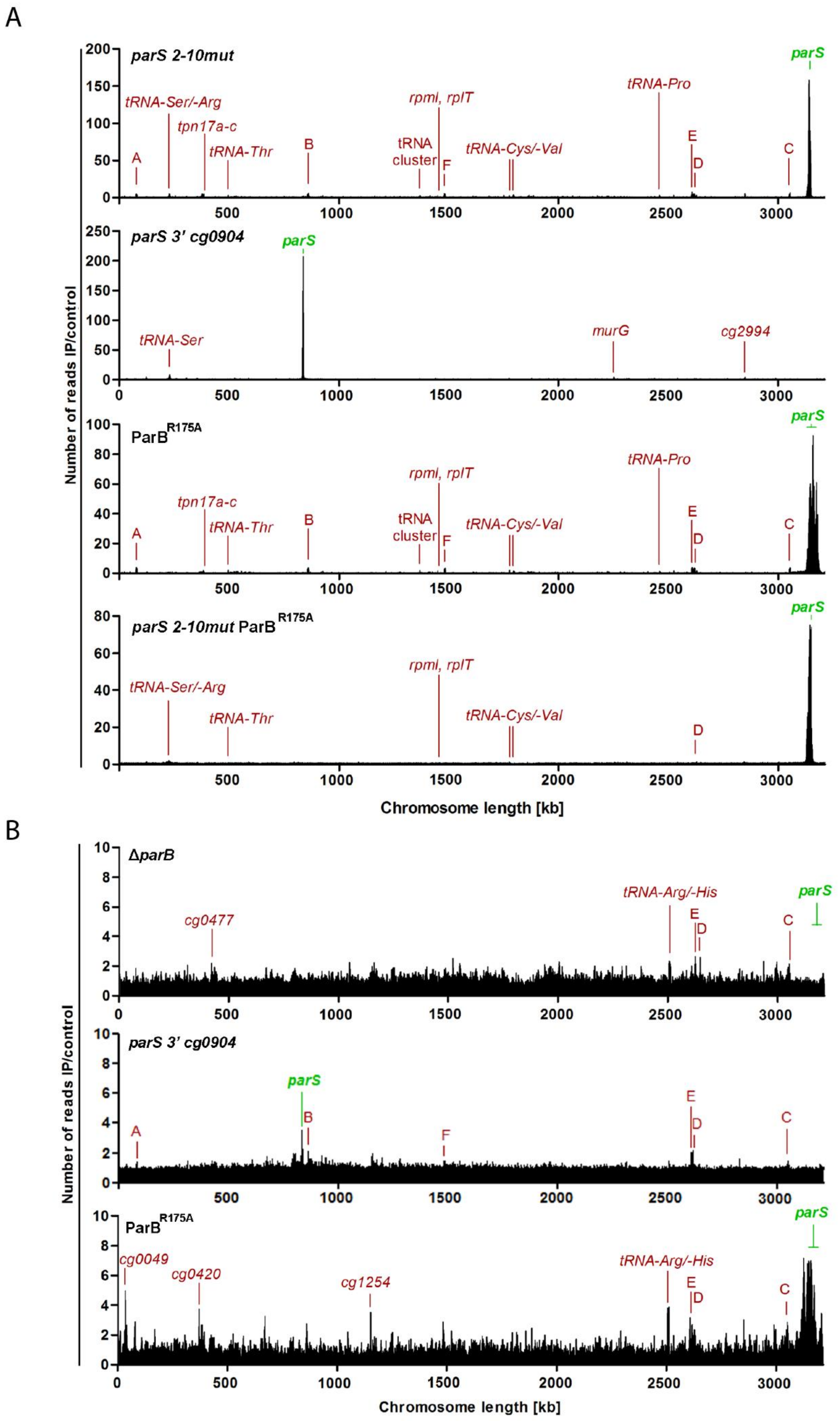


**Figure S4. Whole genome ParB- and SMC-ChIP-seq data.**

**A)** ParB -ChIP-seq analyses in different mutant backgrounds; enrichments at *parS* sites (green) and highly transcribed genes (red) are depicted with a bin size of 0.5 Kb. Fold enrichments of immunoprecipitation relative to extract samples (IP/control) are shown for the entire genome lengths, respectively; x-axes are centered at *terC*. Strains analyzed from top down are: *parB::parB-mCherry* $parS_{2\text{-}10mut}$ (CBK029), *parB::parB-mCherry parS 3' of cg0904* (CBK042), *parB::parB-mCherry* $parS_{2\text{-}9mut}$ (CBK030), $parB{::}parB^{R175A}$*-mCherry* (CBK47) and $parB{::}parB^{R175A}$*-mCherry* $parS_{2\text{-}10mut}$ (CBK048). **B)** DNA-enrichment patters were determined for SMC via ChIP-seq as described above. Genotypes analyzed are *smc::smc-mCherry* in combination with either Δ*parB* (CBK014), *parS* 3' of *cg0904* (CBK045) or $parB{::}parB^{R175A}$ (CBK049).

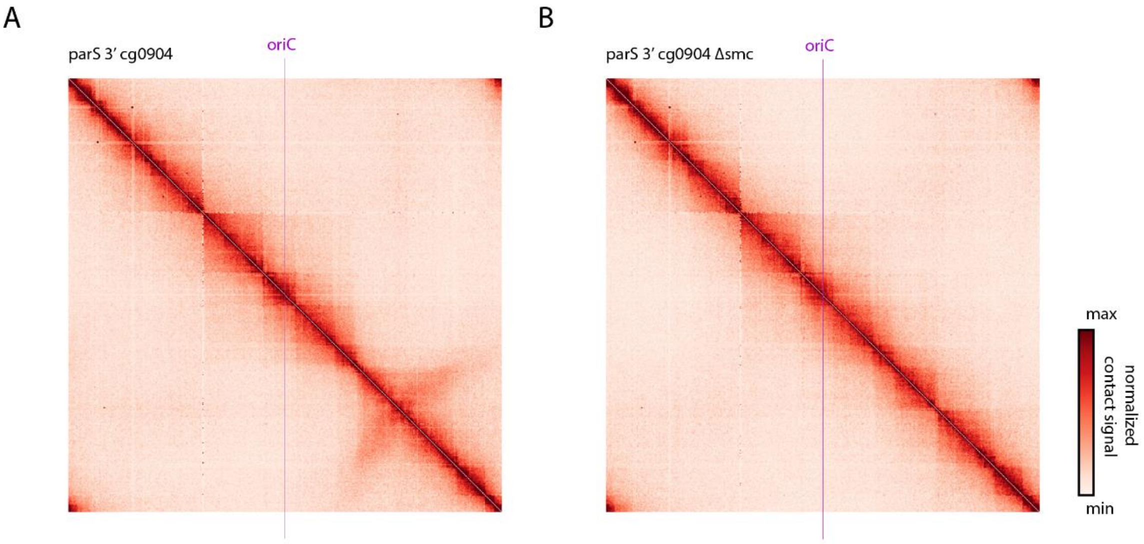


**Figure S5. Whole chromosome organization of *C. glutamicum* mutants.**

Normalized contact maps of mutant strains. **A)** Chromosomal contacts of *parS*$_{1\text{-}10mut}$ *parS 3' cg0904* in a wild type (CBK037). **B)** Chromosomal contacts of *parS*$_{1\text{-}10mut}$ *parS 3' cg0904* in a Δ*smc* background (CBK046). Contact maps are *ori* centered (purple lines).

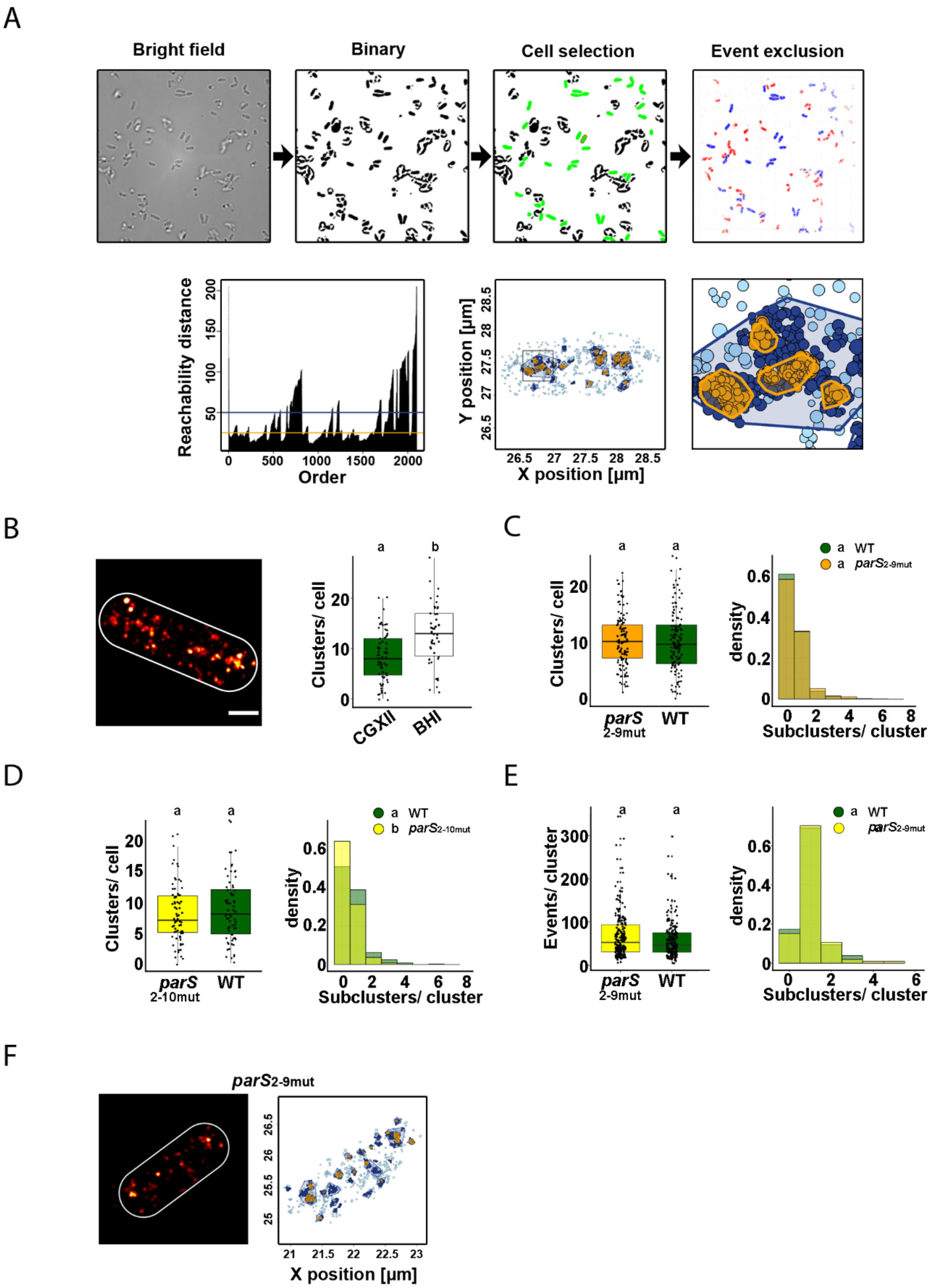


**Figure S6. PALM analyses on cellular ParB localization.**

**A)** Workflow for protein cluster analysis. Top: Bright field image areas were masked out using a binary mask following cell selection (for parameters see Material and Methods) and, accordingly, event exclusion. Below: Illustration of the cluster-ordering in reachability plot (left) showing cluster-order of events within one *C. glutamicum* cell and their reachability distances [nm]. Threshold lines for macro- (dark blue, – 50 nm) and subclusters (yellow, – 35 nm) are plotted with parameters as indicated. Events within the cell are assigned to clusters accordingly; a detail magnification is attached (right). **B)** Left: Gaussian rendered PALM microscopy image (0.71 PSF, 1 px = 10 nM) exemplifying ParB-PAmCherry localization in fast-grown *C glutamicum* cell, strain CBK009. Scale bar, 0.5 µm. Right: Significantly more ParB-PAmCherry macroclusters are present in fast-growing (BHI medium) compared to slow-growing cells (CGXII medium) as indicated by letters above data sets, (Kruskal-Wallis Rank Sum Test: chi-squared = 5.6107, df = 1, $p<0.05$, $cells_{BHI}$: n = 47, $cells_{CGXII}$: n = 68). **C)** Properties of all ParB-PAmCherry clusters per cell were compared between wild type and *parS*$_{2\text{-}9mut}$ mutant (CBK009, CBK031). Differences of events per macroclusters (left, Kruskal-Wallis Rank Sum Test: chi-squared = 1.9737, df = 1, p-value = 0.16) or subcluster numbers per macrocluster (right, Kruskal-Wallis Rank Sum Test: chi-squared = 1.5435, df = 1, p-value = 0.2141) amongst both strain backgrounds are not significant, as indicated by letters above data sets. **D)** ParB-PAmCherry cluster analysis of wild type and *parS*$_{2\text{-}10mut}$ cells (CBK029) as in C). Mean subcluster numbers per macrocluster (Kruskal-Wallis Rank Sum Test: chi-squared = 6.8861, df = 1, $p < 0.05$) and macrocluster size (Kruskal-Wallis Rank Sum Test: chi-squared = 18.923, df = 1, p-value < 0.05) are significantly higher in presence of all *parS* sites. **E)** Comparison of ParB-PAmCherry cluster properties in wild type and *parS*$_{2\text{-}9mut}$ as in Fig. 2H. Kruskal-Wallis Rank Sum test yielded no significant differences of macrocluster sizes (chi-squared = 1.7848, df = 1, p = 0.1816) or subcluster numbers (chi-squared = 0.38145, df = 1, p=0.5368) amongst both strain backgrounds. **F)** Representative *parS*$_{2\text{-}9mut}$ mutant cell analyzed via single molecule localization microscopy. Left: Gaussian rendering of ParB-PAmCherry signals, right: color-coded representation of ParB-PAmCherry events within corresponding cells as described in Fig. 2E).

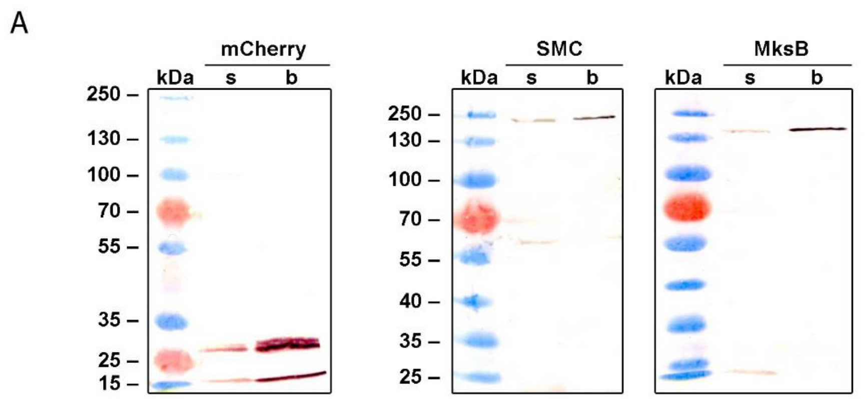


B

| Foci number | |
|---|---|
| ParB | SMC Ø |
| 1 | 1 |
| 2 | 1.9 |
| 3 | 2.7 |
| 4 | 3.2 |
| 5 | 3.9 |
| 6 | 4.6 |
| 7 | 4.5 |

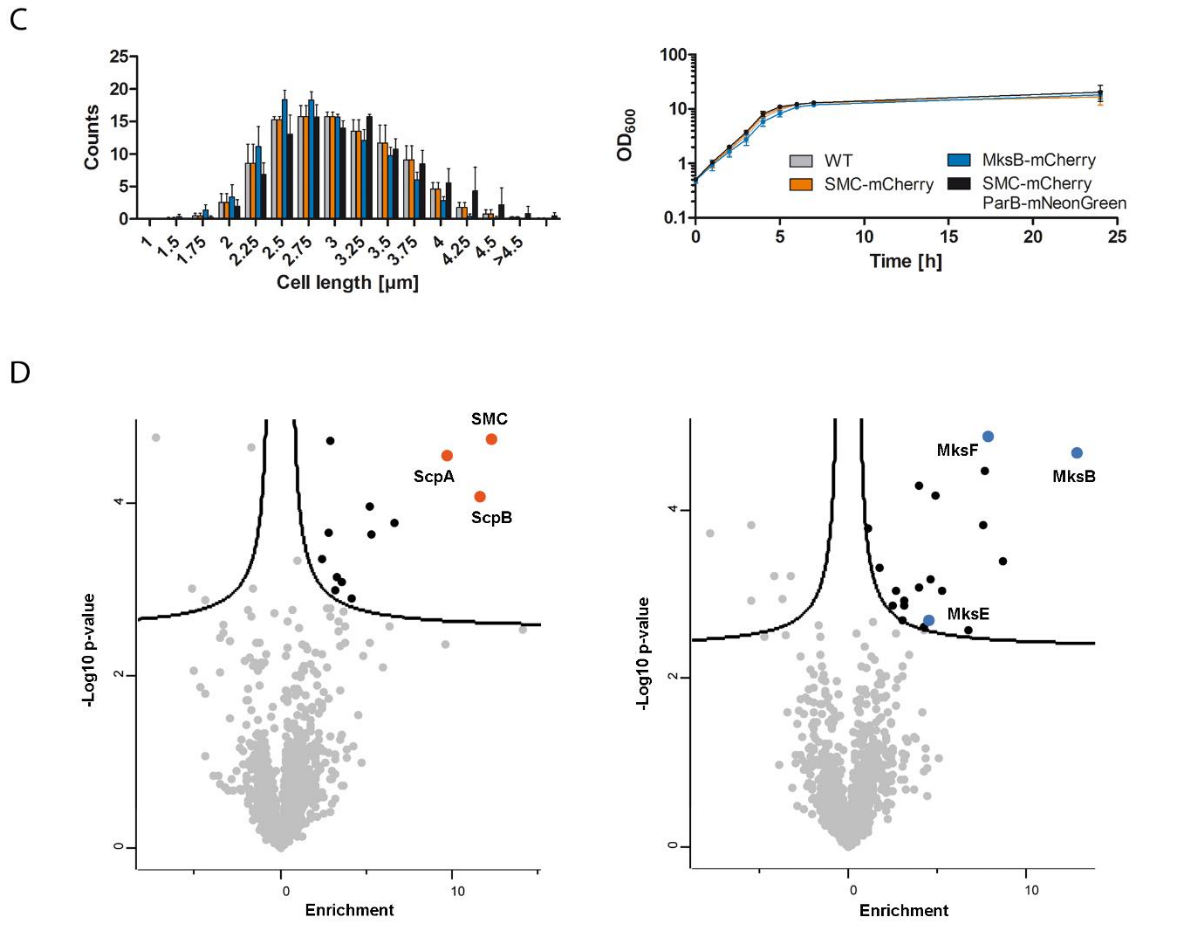


**Figure S7. Validation of full-length fusion proteins SMC-/ MksB-mCherry and co-immunoprecipitation of subunits.**

**A)** Full length fluorescent fusion proteins and their enrichment during immunoprecipitation were validated via western blotting. Whole cell lysates of *C. glutamicum* strains CBK052 (pEKEx2-mCherry), CBK012 (*smc::smc-mCherry*) and CBK017 (*mksB::mksB-mCherry*) strains were used for pulldown experiments. Western blots show proteins SMC-mCherry (155 kDa), MksB-mCherry (151 kDa) and mCherry

(26.7 kDa) trapped on 10 µl magnetic RFP-Trap® beads (b) and in the 10 µl supernatant (s) detected via polyclonal α-mCherry antibody. **B)** Average SMC foci numbers increase in dependence of ParB- clusters per cell in strain *smc::smc-mCherry parB::parB-mNeonGreen* (CBK013), n>200. **C)** Growth curves and cell length distributions (n>1000) of *C. glutamicum* strains grown in BHI harboring allelic replacements of condensin subunits or ParB by fluorescent versions as indicated. Values derive from triplicates, error bars display standard deviations; $\mu_{WT}$=0.64 $h^{-1}$, $\mu_{smc\text{-}mCherry}$=0.67 $h^{-1}$ (CBK012), $\mu_{mksB\text{-}mCherry}$=0.60 $h^{-1}$ (CBK015) $\mu_{smc\text{-}mCherry\ parB\text{-}mNeonGreen}$=0.68 $h^{-1}$ (CBK014). **D)** Condensin subunit interactions identified by pull-down experiments and mass spectrometry. Cell lysates of *C. glutamicum smc::smc-*mCherry, *mksB::mksB-mCherry* and a negative control strain (CBK052) were used for co-immunoprecipitations via anti-mCherry agarose beads. Volcano plots show the difference in means plotted against the -log10 adjusted p value for each protein identified by mass spectrometry; condensin subunits are highlighted respectively. The cutoff curves indicate significant hits (two-tailed t-test, p<0.05, FC >0.1).

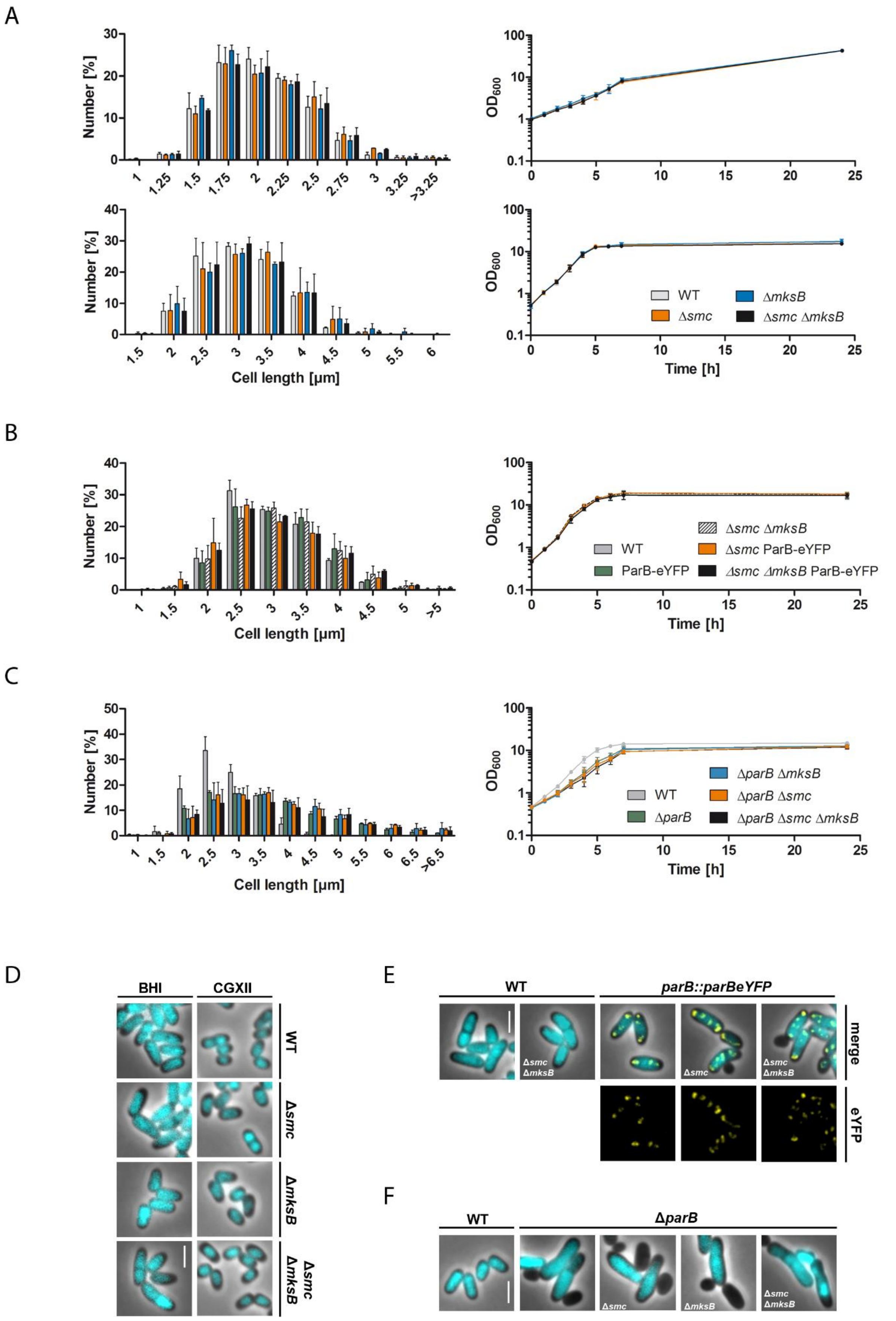


**Figure S8. Characterization of growth and cell length phenotypes of condensin deletion strains.**

**A)-C)** Growth curves and cell length distributions (n>1000) of *C. glutamicum* mutant strains. Data points derive from biological triplicates where standard deviations are indicated by error bars; growth rates are listed. **A)** Growth experiments performed in CGXII medium (top): $\mu_{WT}$=0.26 $h^{-1}$, $\mu_{\Delta smc}$=0.22 $h^{-1}$ (CDC026), $\mu_{\Delta mksB}$=0.27 $h^{-1}$ (CBK001), $\mu_{\Delta smc\ \Delta mksB}$=0.25 $h^{-1}$ (CBK004); in BHI medium (below): $\mu_{WT}$=0.71 $h^{-1}$, $\mu_{\Delta smc}$=0.70 $h^{-1}$, $\mu_{\Delta mksB}$=0.72 $h^{-1}$, μΔ*smc*Δ*mksB*=0.70 $h^{-1}$. **B)** Analysis in BHI medium: $\mu_{WT}$=0.70 $h^{-1}$, $\mu_{ParB\text{-}eYFP}$=0.69 $h^{-1}$ (CBK007), $\mu_{\Delta smc\Delta mksB}$=0.72 $h^{-1}$ (CBK004), $\mu_{\Delta smc\ ParBeYFP}$=0.70 $h^{-1}$ (CBK010), $\mu_{\Delta smc\ \Delta mksB\ ParBeYFP}$=0.71 $h^{-1}$ (CBK011). **C)** Analysis in BHI medium: $\mu_{WT}$=0.67 $h^{-1}$, $\mu_{\Delta parB}$=0.51 $h^{-1}$ (CDC003), $\mu_{\Delta parB\ \Delta smc}$=0.46 $h^{-1}$ (CBK002), $\mu_{\Delta parB\ \Delta mksB}$=0.49 $h^{-1}$ (CBK003), $\mu_{\Delta parB\ \Delta smc\ \Delta mksB}$ =0.43 $h^{-1}$ (CBK005). **D)-F)** Phenotypes of exponentially grown *C. glutamicum* mutant cells exemplified in overlays of ParB-eYFP fluorescence (yellow) with Hoechst-stained DNA (cyan) and the phase contrast image, respectively. The eYFP fluorescence channel is additionally depicted in separate images. Scale bar, 2 µm.

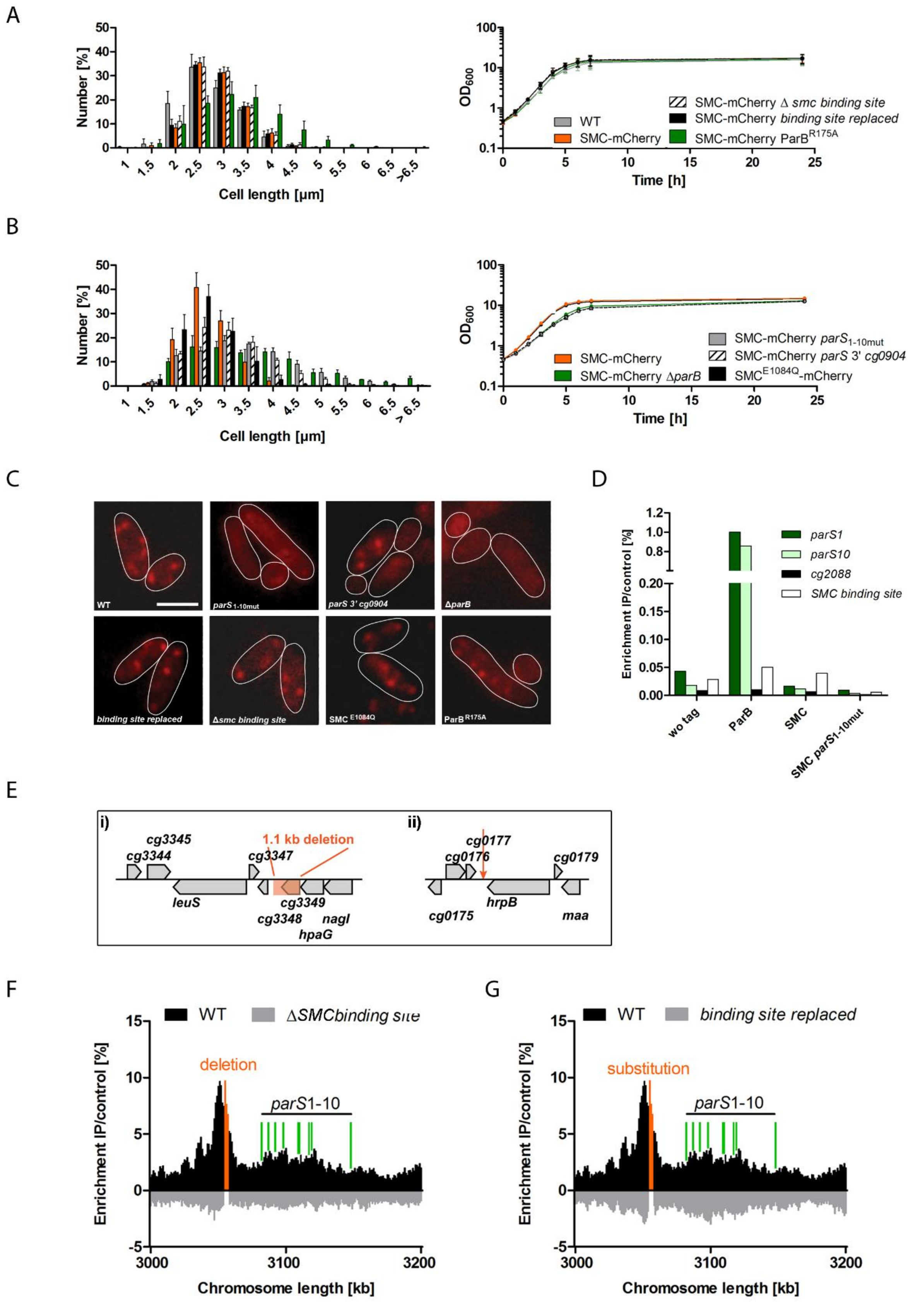


**Figure S9. Characterization of *C. glutamicum smc::smc-mCherry* mutant strains.**

**A)** Analysis of cell lengths distributions, n>1000 (left) and growth in BHI medium (right) of *smc::smc-mCherry* mutant strain derivatives, growth rates: $\mu_{WT}$=0.63 $h^{-1}$, $\mu_{SMC\text{-}mCherry}$=0.69 $h^{-1}$ (CBK012), $_{SMC\text{-}mCherry\ \Delta SMC\ binding\ site}$ =0.68 $h^{-1}$ (CBK033), $\mu_{SMC\text{-}mCherry\ SMC\ binding\ site\ replaced}$ =0.68 $h^{-1}$ (CBK034), $\mu_{SMC\text{-}mCherry\ ParBR175A}$ =0.69 $h^{-1}$ (CBK049). Error bars indicate standard deviations derived from biological triplicates. **B)** Growth analyses using BHI medium as shown in A): $\mu_{SMC\text{-}mCherry}$=0.71 $h^{-1}$ (CBK012), $_{SMC\text{-}mCherry\ \Delta parB}$ =0.60 $h^{-1}$ (CBK014), $\mu_{SMC\text{-}mCherry\ parS1\text{-}10mut}$=0.56 $h^{-1}$ (CBK032), $\mu_{SMC\text{-}mCherry\ parS\ 3'\ cg904}$=0.55 $h^{-1}$ (CBK045), $\mu_{SMC\text{-}mCherryE1084Q}$=0.72 $h^{-1}$ (CBK050). **C)** Localization of SMC-mCherry foci in strain backgrounds described in A) and B). Images show mCherry fluorescence in representative cells; cell outlines are indicated (white lines). Scale bar, 2 µm. **D)** ChIP-qPCR of the wild type and cells harboring ParB- or SMC-mCherry tagged proteins in a wild type (SMC) or a *parS*-mutation ($parS_{1\text{-}10mut}$) background. Genomic loci including *parS1*, *parS10*, *cg2088* and the region of additional SMC binding upstream of *parS* sequences were amplified; values derive from biological duplicates normalized to ParB signal at *parS*1. **E)** Scheme showing mutation sites within the *C. glutamicum* CBK034 genome with a partially deleted SMC-enrichment region upstream of *parS* reinserted into intergenic region 3' of *cg0177*. i) ChIP-seq enrichment site of SMC-mCherry protein covers genes *cg3345* to *hpaG*. Deletion including the non-essential gene *cg3349* is indicated in orange (31.3 Mb). ii) Arrow points to intergenic insertion site of the 1.1 Kb fragment (0.15 Mb). **F)** Partial deletion of SMC-enrichment site collapses entire SMC-binding nearby. *In vivo* ChIP-seq was performed using exponentially grown *C. glutamicum* SMC-mCherry expressing wild type (upper y-axis, black) or CBK034 cells (lower y-axis, gray). The fold enrichment of the IP versus a control is displayed in a bin size of 0.5 Kb. The deletion site and *parS* sequences are indicated in orange and by green lines, respectively. **G)** SMC-ChIP-seq analysis in CBK035 cells with a partial substitution of the SMC-binding site by a noncoding *B. subtilis* genome sequence of identical size which does not lead to SMC enrichment.

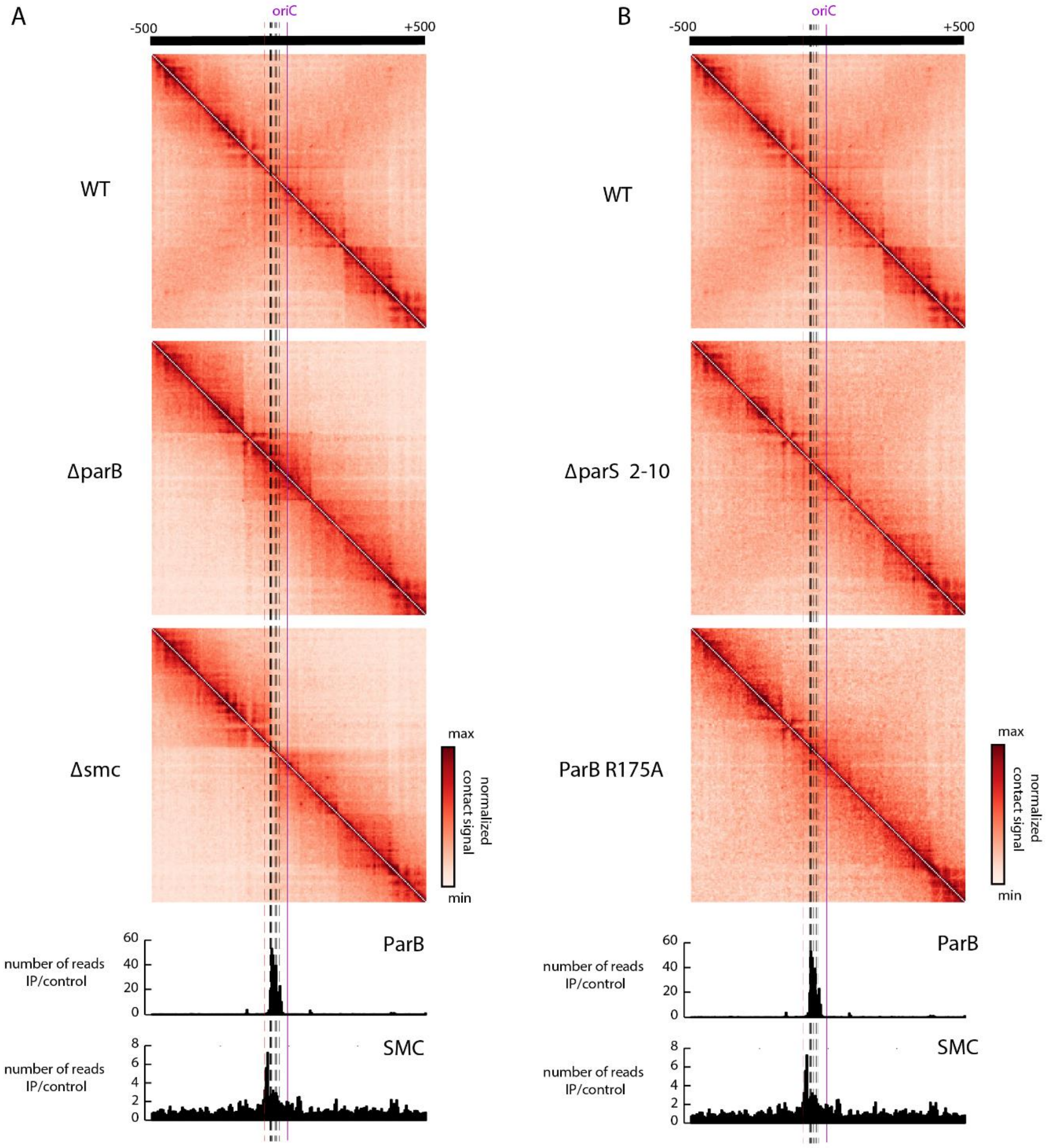


**Figure S10. Disorganization of *oriC*-regions in Par and SMC mutants.**

**A)** Magnification of normalized genomic contact matrices of wild type, Δ*parB* and Δ*smc*, (CDC003, CDC026) encompassing 500 Kb regions surrounding *oriC* (purple line). *parS* sites are indicated as dashed lines. Color codes as in Fig. 1 were applied. ParB and SMC enrichment zones are shown below the contact maps (ChIP signal relative to the input in 5 Kb bins). **B)** Magnification of normalized genomic contact matrices of wild type, *parS*$_{2\text{-}10mut}$ and *parB::parB*$^{R175A}$ (CBK023, CBK047) as in A.

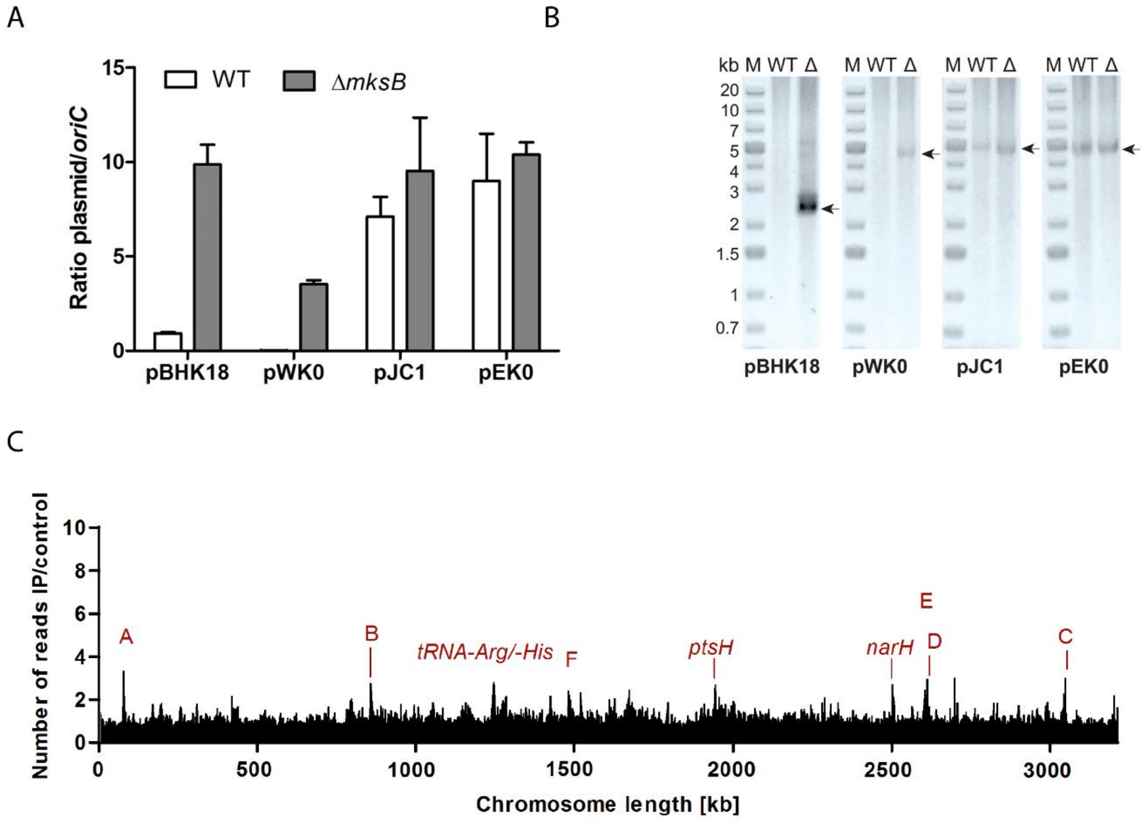


**Figure S11. MksB impacts on plasmid copy numbers.**

**A)** Plasmid copy numbers of low copy (pBHK18 and pWK0) and high copy number vectors (pJC1 and pEC0) relative to *oriC* numbers per cell, assayed by qPCR. Ratios were compared between *C. glutamicum* wild type and Δ*mksB* mutant cells grown in BHI medium without addition of plasmid selection antibiotic after overnight pre-incubation without antibiotic, error bars display standard deviations (n=3). **B)** Plasmids named in A) were extracted from *C. glutamicum* wild type and *mksB* deletion strains grown in BHI medium including selection antibiotic, visualization of extracted DNA on 1% agarose gels (corresponds to yield from approx. $1*10^9$ cells each). Arrows indicate size of plasmid DNA. **C)** Anti-mCherry-ChIP-seq analysis of *mksB*::*mksB-mCherry* strain CBK015 as described in Fig. S4.

Figure S12

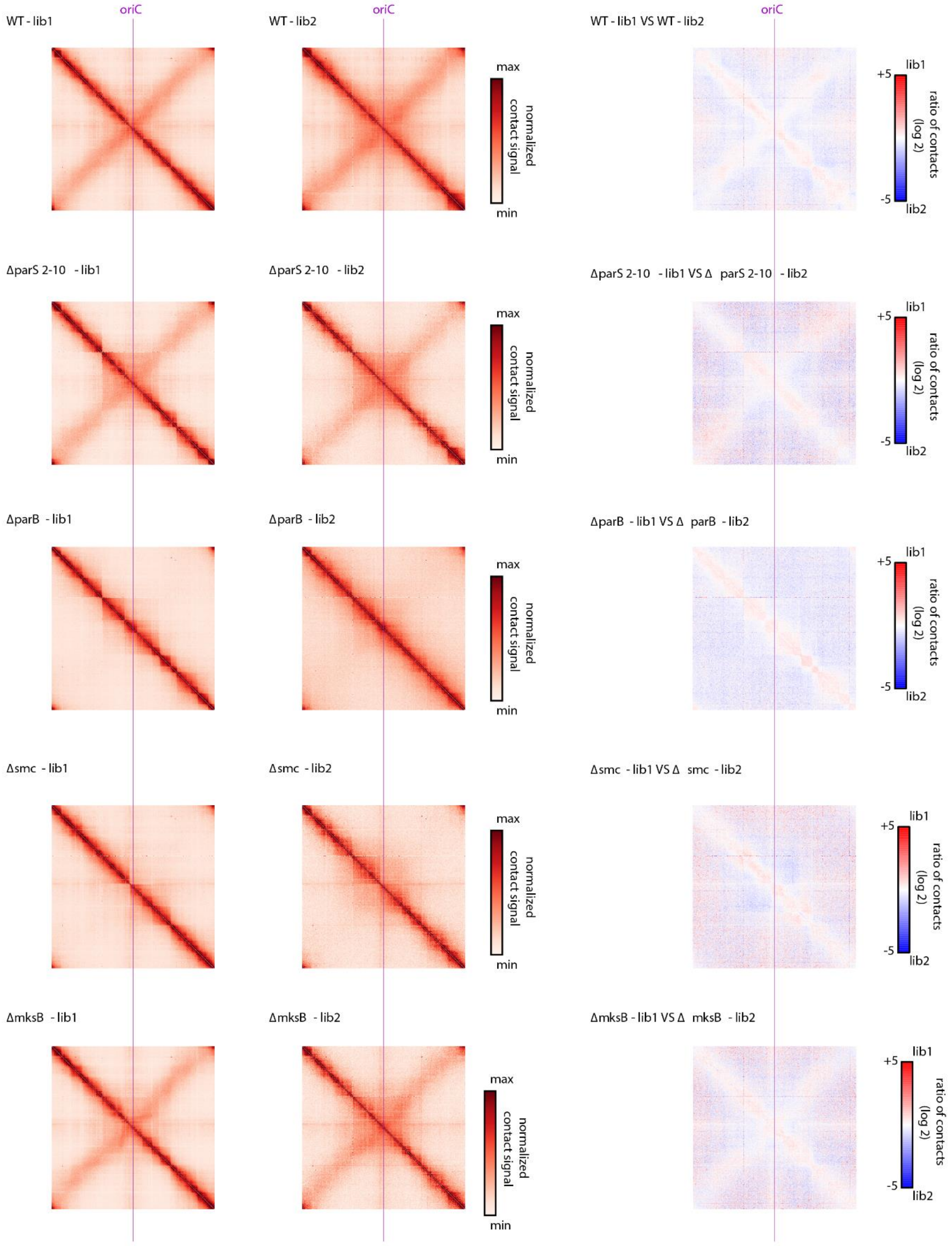


**Figure S12 Replicates of chromosomal contact maps in *C. glutamicum* mutants.**

Normalized contact maps of replicates of wild type, *parS*$_{2\text{-}10mut}$ (CBK023), Δ*parB* (CDC003), Δ*smc* (CDC026) and Δ*mksB* (CBK001) mutants centered at *oriC*. Color codes as in Fig. 1 were applied. Differential maps of each strain correspond to the log of the ratio (replicate 1 norm/ replicate 2 norm); color scales indicate contact enrichment in replicate 2 (blue) or replicate 1 (red).

**Table S1. Percentage of DNA-free cells of relevant *C. glutamicum* RES167 strains used in this study.**

Color code visualizes severity of nucleoid missegregation: virtually none (blue), up to 10 % (green) and more than 20 % (red) DNA-free cells, (n>1000).

| Strain | Genotype | Anucleate cells [%] |
|---|---|---|
| **RES 167** | wild type | 0 |
| **CDC002** | *ΔparB* | 26.14 |
| **CDC026** | *Δsmc* | 0.1 |
| **CBK001** | *ΔmksB* | 0 |
| **CBK002** | *ΔparB Δsmc* | 27.08 |
| **CBK003** | *ΔparB ΔmksB* | 24.95 |
| **CBK004** | *Δsmc ΔmksB* | 0 |
| **CBK005** | *ΔparB Δsmc ΔmksB* | 25.62 |
| **CBK006** | *parB::parB-mCherry2* | 0 |
| **CBK007** | *parB::parB-eYFP* | 0.1 |
| **CBK010** | *Δsmc parB::parB-eYFP* | 4.25 |
| **CBK011** | *Δsmc ΔmksB parB::parB-eYFP* | 3.98 |
| **CBK012** | *smc::smc-mCherry* | 0 |
| **CBK016** | *smc::smc-mCherry ΔparB* | 24.18 |
| **CBK017** | *mksB::mksB-mCherry* | 0 |
| **CBK025** | *parS*$_{2\text{-}10mut}$ | 0.19 |
| **CBK026** | *parS*$_{1\text{-}10mut}$ | 29.27 |
| **CBK027** | *parS*$_{2\text{-}10mut}$ *parB::parB-eYFP* | 6.71 |
| **CBK028** | *parS*$_{1\text{-}10mut}$ *parB::parB-eYFP* | 28.5 |
| **CBK034** | *parS*$_{1\text{-}10mut}$ *smc::smc-mCherry* | 30.38 |
| **CBK036** | *smc::smc-mCherry Δ SMC loading site* | 0 |
| **CBK037** | *smc::smc-mCherry SMC loading site replaced* | 0 |
| **CBK042** | *parS*$_{1\text{-}10mut}$ *parS* 3‘ *cg0108 parB::parB-eYFP* | 25.33 |
| **CBK043** | *parS*$_{1\text{-}10mut}$ *parS* 3‘ *cg0904 parB::parB-eYFP* | 23.48 |
| **CBK045** | *parS*$_{1\text{-}10mut}$ *parS* 3‘ *cg2563 parB::parB-eYFP* | 24.28 |
| **CBK046** | *parS*$_{1\text{-}10mut}$ *int::parS parB::parB-eYFP* | 24.98 |
| **CBK047** | *parS*$_{1\text{-}10mut}$ *parS* 3‘ *cg0904 smc::smc-mCherry* | 25.07 |
| **CBK051** | *parB::parB-*$^{R175A}$*mCherry2* | 17.83 |
| **CBK052** | *parB::parB-*$^{R175A}$*mCherry2 parS*$_{2\text{-}10mut}$ | 31.74 |
| **CBK053** | *smc::smc-mCherry parB::parB-*$^{R175A}$ | 19.09 |
| **CBK054** | *smc::smc*$^{E1084Q}$*-mCherry* | 0 |

**Table S2. Oligonucleotides used for strain construction and qPCR.**

Restriction sites are indicated by an underline; primer modifications resulting in point mutations are highlighted in green and bold letters indicate *parS* sites.

| **Oligonucleotides**<br>**Strain construction** | **Sequence 5'–3'** | **Restric-tion sites** |
|---|---|---|
| **DivIVA-XbaI-F** | CATTCTAGAGCCGTTGACTCCAGCTG | XbaI |
| **DivIVA-BamHI-R** | CATGGATCCCTCACCAGATGGCTTGTTGT | BamHI |
| **ParB-XbaI-F** | CATTCTAGAGGCTCAGAACAAGGGTT | XbaI |
| **ParB-BamHI-R** | CATGGATCCTTGGCCCTGGATCAAGGACA | BamHI |
| **SMC-XbaI-F** | CGCTCTAGAGATGTATTTGAAATCGTTGACGCTCAAGGGG | XbaI |
| **SMC-KpnI-R** | ATAGGTACCCGCCCCGCCACAGTTTCCA | KpnI |
| **ScpA-XbaI-F** | CGCTCTAGAGGTGCAGCTCGATAATTTT | XbaI |
| **ScpA-XmaI-R** | TATCCCGGGGCTCCCAGTCAC | XmaI |
| **ScpB-XbaI-F** | CGCTCTAGAGATGGAATCAATCTTGT | XbaI |
| **ScpB-XmaI-R** | ATACCCGGGGGAAGTCTTCAT | BamHI |
| **MksB-XmaI-F** | TAACCCGGGTAGTGACCAGCGAACAAGCTTTA | XmaI |
| **MksB-KpnI-R** | CGCGGTACCCATTTCTCGATCCTAGAGAAACTGG | KpnI |
| **MksE-XbaI-F** | CTATCTAGAGATGAATGATCAGCTGTGG | XbaI |
| **MksE-XmaI-R** | ATACCCGGGGCTTCTGTTCC | XmaI |
| **MksF-XbaI-F** | ATATCTAGAGATGACCGTTGTATCGCA | XbaI |
| **MksF-XmaI-R** | ATACCCGGGGTTTATCCATCTC | XmaI |
| **MksG-XbaI-F** | ATATCTAGAGATGCCATTGTTTATCGACGAC | XbaI |
| **MksG-XmaI-R** | ATACCCGGGGCCCACGAATTACTTT | XmaI |
| **SMC-seq-689bp-F** | CTGGCTTTGAGATCGTGAAG | |
| **SMC-seq-1589bp-F** | AGGCGCTGGCTGGCGAGG | |
| **MksB-seq-875bp-F** | CACTGAAGAAGGCGCTGCCG | |
| **MksB-seq-1595bp-F** | GCGTGGTGACAACGGGGGAG | |
| **pKNT25/pUT18-seq-F** | GAGTTAGCTCACTCATTAGGCACC | |
| **pKNT25-seq-R** | GGCGTTTGCGTAACCAGCCTG | |
| **pUT18-seq-R** | TGATCACGCCGATATTCATGTC | |
| **pUT18C-seq-F** | ATGTACTGGAAACGGTGCCG | |
| **pUT18C-seq-R** | TAACTATGCGGCATCAGAGCAG | |
| **pKT25-seq-F** | ACATGTTCGCCATTATGCCGCATC | |
| **pKT25-seq-R** | CGAAAGGGGGATGTGCTGCAA | |
| **pUT18C-mcs-HindIII-F** | TATAAGCTTAGCCGCCAGCGAGG | HindIII |
| **pKNT25-mcs-NheI-F** | ATAGCTAGCGCCCAATACGCAAACC | NheI |
| **pUT18-mcs-PvuII-F** | ATACAGCTGGCACGACAGGTTTCC | PvuII |
| **pKT25-mcs-HindIII-F** | CGGAAGCTTTAATGCGGTAGTTTAT | HindIII |

| | | |
|---|---|---|
| **pUT18(C)/pK(N)T25-mcs-KpnI-R** | TAAGGTACCATTACCCGGGGATCCTCTAGA | KpnI |
| **Δsmc-BamHI-up-F** | CAGGGATCCAGC ACA CGC GTG TGA AAA | BamHI |
| **Δsmc-up-R** | **GGAATGAGTATGGAAGTTGG**AACGGGGTTAAAGTCTAG | |
| **Δsmc-D-F** | **CCAACTTCCATACTCCATCC**AACCTGCGCTGAGCACGG | |
| **Δsmc-EcoRI-D-R** | CAGGAATTCTGC GAA GAG CTT TTC GGT | EcoRI |
| **Δsmc-seq-700up-F** | GGTCACTGCAGGAACACT | |
| **Δsmc-seq-700D-R** | AGTTCTTGAACTCCGCCG | |
| **ΔmksB-HindIII-up-F** | CAGAAGCTTCAAGATGCGCTCAATGCT | HindIII |
| **ΔmksB-PstI-up-R** | ATACTGCAGTCACTTCTGTTCCTCTT | PstI |
| **ΔmksB-PstI-D-F** | ATACTGCAGATTCAGGGCAATT | PstI |
| **ΔmksB-XbaI-D-R** | CAGTCTAGAATTTGGTATGCACGCCTT | XbaI |
| **ΔmksB-seq-700up-F** | ATTACAGGATGGCAGTTCATCAG | |
| **ΔmksB-seq-700D-R** | CAGAATTACTTGCGGTTCTTGTAATTC | |
| **PAmCherry-SalI-F** | CATGTCGACATGGTGAGCAAGGGCGAG | SalI |
| **PAmCherry-XbaI-R** | CATTCTAGACTTGTACAGCTCGTCCAT | XbaI |
| **ParB-N-ter-SalI-F** | CAGGTCGACATGGCTCAGAACAAGGGTTCC | SalI |
| **ParB-seq-800D-R** | CAGCTGCAGCCAACCCCGATGACCTGG | PstI |
| **SMC-HindIII-up-F** | CATAAGCTTAAGAAGTCGATGCG | HindIII |
| **SMC-SphI-up-R** | CATGCATGCCCCCGCCACAGTTT | SphI |
| **mCherry-XbaI-F** | CATTCTAGAATGGTGAGCAAGGGCGAG | XbaI |
| **mCherry-BamHI-R** | CATGGATCCTTACTTGTACAGCT | BamHI |
| **SMC-BamHI-D-F** | CATGGATCCTAAAACCTGCGCTG | BamHI |
| **SMC-EcoRI-D-R** | CATGAATTCTGCGAAGAGCTTTT | EcoRI |
| **MksB-HindIII-up-F** | CATAAGCTTAAGCCTTCGCCCGTTATG | HindIII |
| **MksB-Sph-up-R** | CATGCATGCTTTCTCGATCCTAGAGAA | SphI |
| **mCherry-XbaI-R** | GCGTCTAGATTACTTGTACAGCTCGTC | XbaI |
| **MksB-BamHI-D-F** | CATGGATCCTAACATGCCATTGTTTAT | BamHI |
| **MksB-EcoRI-D-R** | CATGAATTCATTTGGTATGCACGCCTT | EcoRI |
| **ΔSMCload-HindIII-up-F** | ATAAAGCTTGGGCTAACAAGGCTCTCG | HindIII |
| **ΔSMCload-up-R** | GGAGCGGAGTATGGAAGTTGGTTATTCAAACACCGTCTTGTTTTCA | |
| **ΔSMCload-D-F** | CCAACTTCCATACTCCGCTCCTAGGTAAGGAAGATGATTTTGGGG | |
| **ΔSMCload-SalI-D-R** | ATAGTCGACCTGCGAGCTCTCCTGA | SalI |
| **cg0177-HindIII-up-F** | ATAAAGCTTAATATCCACAACAGTCACAGTC | HindIII |
| **cg0177-SalI-up-R** | ATAGTCGACGAACGAGTATTTGTATTTCAAT | SalI |
| **SMCload-SalI-F** | ATAGTCGACATGACAACTTTCCACGAT | SalI |
| **SMCload-XmaI-R** | ATACCCGGGTGACGCAATACGTTATG | XmaI |
| **cg0177-XmaI-D-F** | ATACCCGGGGCTTTTAAGTTTTCTCG | XmaI |
| **cg0177-EcoRI-D-R** | ATAGAATTCGTACCCAGCGGGAATCA | EcoRI |
| **cg0177-seq-700up-F** | TGGTTGTCGTCATGAGCGTC | |
| **cg0177-seq-700D-R** | TGAAATTCATCCACGAACAC | |
| **SMCload-SphI-up-R** | TATGCATGCTTATTCAAACACCGTCTTGTTT | SphI |
| **SMCloadr-SphI-F** | CGTGCATGCGGAGCGGGGAAGATTGCA | SphI |
| **SMCloadr-PstI-R** | CGTCTGCAGAATCCATGCTGTAGATCGGG | PstI |

| | | |
|---|---|---|
| **SMCload-PstI-D-F** | ATACTGCAGTAGGTAAGGAAGATGATTTTGG | PstI |
| **parS1mut-HindIII-up-F** | ATAAAGCTTGCGTTACTTTGGTGCTGATGC | HindIII |
| **parS1mut-XmaI-up-R** | ATACCCGGGACACCTTGCGTGGCGATG | XmaI |
| **parS1mut-XmaI-D-F** | TATCCCGGGAGACTTTGCAAAAAATCATCGCC | XmaI |
| **parS1mut-EcoRI-D-R** | TATGAATTCGAAAATTTTCAGCAGCGCGCCGG | EcoRI |
| **parS2mut-HindIII-up-F** | ATAAAGCTTAGTTCGCTGAAGCTGGCG | HindIII |
| **parS2mut-XmaI-up-R** | TTTCCCGGGATTCGTTGCCACATTTAAAG | XmaI |
| **parS2mut-XmaI-D-F** | TTTCCCGGGATACTTTGCCCTAAAGTACA | XmaI |
| **parS2mut-EcoRI-D-R** | ATAGAATTCCATCGTGGCTGAACCC | EcoRI |
| **parS3mut-HindIII-up-F** | ATAAAGCTTGCTTTAGCCAATGCCG | HindIII |
| **parS3mut-XmaI-up-R** | TTTCCCGGGATTCGTTTCGCGTGGGAAC | XmaI |
| **parS3mut-XmaI-D-F** | TTTCCCGGGATACTTCTGCTGATTTTTGTCT | XmaI |
| **parS3mut-EcoRI-D-R** | GCGGAATTCTGAGACTGACTTTCCTTCTG | EcoRI |
| **parS4mut-HindIII-up-F** | ATAAAGCTTCGATTATTGGTGTCGCCCT | HindIII |
| **parS4mut-XmaI-up-R** | ATACCCGGGATTCGTCGCCATTTTTTGT | XmaI |
| **parS4mut-XmaI-D-F** | ATACCCGGGATACTTCGCCATTTTTCTA | XmaI |
| **parS4mut-EcoRI-D-R** | ATAGAATTCTAAACGTACTGATATTTATGG | EcoRI |
| **parS5mut-HindIII-up-F** | ATAAAGCTTGGAGCTTGCGGAGCTTTT | HindIII |
| **parS5mut-SalI-up-R** | ATAGTCGACATTCGTTCTCCCTCTTTTCG | SalI |
| **parS5mut-SalI-D-F** | CGCGTCGACATACTTTCTCCCTCTTTAG | SalI |
| **parS6mut-XmaI-up-R** | ATACCCGGGATTCGTTGACCGAG | XmaI |
| **parS6mut-XmaI-D-F** | ATACCCGGGATACTTTGACCTTAAAGA | XmaI |
| **parS6mut-EcoRI-D-R** | ATAGAATTCTTCAGGAGCAGTTCCAGC | EcoRI |
| **parS7mut-HindIII-up-F** | TGTAAGCTTATGTGGCCCAGCATGAC | HindIII |
| **parS7mut-XmaI-up-R** | ATACCCGGGATTCGTTTCCCTCATTTGC | XmaI |
| **parS7mut-XmaI-D-F** | GTACCCGGGATACTTTGAAAGAAGTCAGA | XmaI |
| **parS7mut-EcoRI-D-R** | ATAGAATTCAAAGTGTTCGGGGCGAT | EcoRI |
| **parS8mut-HindIII-up-F** | ATAAAGCTTCTCTGCATCATTGAAAGCCAC | HindIII |
| **parS8mut-XmaI-up-R** | ATACCCGGGATTCGTTGGCATTCTTTGGAT | XmaI |
| **parS8mut-XmaI-D-F** | ATACCCGGGATACTTTGGCATTCTGGAGGGTTG | XmaI |
| **parS8mut-EcoRI-D-R** | GTCGAATTCCTGAAGGCAATCCCGTTCCGAATG | EcoRI |
| **parS9mut-HindIII-up-F** | ATAAAGCTTGAAGTGTCCTACGAGCAGCTCG | HindIII |
| **parS9mut-SalI-up-R** | ATAGTCGACATTCGTTGAAAAACGCTCCAC | SalI |
| **parS9mut-SalI-D-F** | ATAGTCGACATACTTTGCAACAAAAATGGCG | SalI |
| **parS10mut-XmaI-up-R** | ATACCCGGGATTCGTTGAGGGTTCATC | XmaI |

| | | |
|---|---|---|
| **parS10mut-XmaI-D-F** | ATACCCGGGATACTTTGAGGGTTTTACACCC | XmaI |
| **parS10mut-EcoRI-D-R** | ATAGAATTCGTGACTATCGGCGCGGAGTA | EcoRI |
| **parS-cg0108-SalI-up-F** | ATAGTCGACCGCCGCTTCACCATGA | SalI |
| **parS-cg0108-up-R** | **TGTTTCACGTGAAACA**GAATTCCGTGTAGCCGTGTCTGGG | |
| **parS-cg0108-D-F** | CGGAATTC**TGTTTCACGTGAAACA**AGTGCTACAGTTCGACTTCACG | |
| **parS-cg0108-XmaI-D-R** | ATACCCGGGCGTAACAATCGTGGCGAAAG | XmaI |
| **cg0108-seq-400up-F** | GCGTCGTAAAGCAATTAAAGGC | |
| **cg0108-seq-200D-R** | CACAAATTGGCGCCTATATAGAT | |
| **parS-cg0904-HindIII-up-F** | ATAAAGCTTCGTATTCGTACTGCCGAGT | HindIII |
| **parS-cg0904-up-R** | **TGTTTCACGTGAAACA**CGGACACGTCCATCAGCG | |
| **parS-cg0904-D-F** | TGTCCG**TGTTTCACGTGAAACA**TTACTTTGGCTTTTCGCAGAAG | |
| **parS-cg0904-NheI-D-R** | ATAGCTAGCTCACATAACCCTTTCGTTAC | NheI |
| **cg0904-seq-100up-F** | TCCGGGTACCACTGTGG | |
| **cg0904-seq-100D-R** | TAACCACCTGAAGCGCTT | |
| **parS-cg2563-HindIII-up-F** | ATTAAGCTTCCGCGCTGACTGGTCTGCA | HindIII |
| **parS-cg2563-up-R** | **TGTTTCACGTGAAACA**CGAAGACTCCCCGAAACTCAC | |
| **parS-cg2563-D-F** | GAGTCTTCG**TGTTTCACGTGAAACA**GCTGCCTAGTTTGGTGTCCAAG | |
| **parS-cg2563-NheI-D-R** | ATAGCTAGCAAGTATTAACTCCCTCGGAAA | NheI |
| **cg2563-seq-200up-F** | CCTTCCGCTGTACTCGATCA | |
| **cg2563-seq-300D-R** | AGCATAGGCATAAGCGCAGT | |
| **parS-Δint-HindIII-up-F** | ATAAAGCTTATTACCAGGAGCGCC | HindIII |
| **parS-Δint-up-R** | **TGTTTCACGTGAAACA**CCGTTTGTTATGTGGACCCTAC | |
| **parS-Δint-D-F** | ACGG**TGTTTCACGTGAAACA**AACGAAACAGTCTTGACCAGCATA | |
| **parS-Δint-NheI-D-R** | ATAGCTAGCGGCGGCATCGTCAC | NheI |
| **Δint-seq-700up-F** | AGCAGATAAAGTTCCAATTGAATGG | |
| **Δint-seq-700D-R** | TTTTCCCAGAACCAGCACC | |
| **ParB-N-ter-HindIII-F** | GCCAAGCTTATGGCTCAGAACAAGGGTTCC | HindIII |
| **ParB-R175A-R** | ACGAGCCTCACCCATGATCAGCT | |
| **ParB-R175A-F** | CATGGGTGAGGCTCGTTGGC | |
| **ParB-C-ter-SalI-R** | ATAGTCGACTTGGCCCTGGATCAAGGA | SalI |
| **ParB-seq-800up-F** | CAGGGTACCATTCATGGGCTTAAAGTTCTC | KpnI |
| **E1084Q-HindIII-up-F** | ATAAAGCTTTCGCAGAATTGCTGCG | HindIII |

| | | |
|---|---|---|
| **E1084Q-up-R** | TCTAGAGCTGCTTCCACCTGATCC | |
| **E1084Q-D-F** | CAGGTGGAAGCAGCTCTAGATGATG | |
| **E1084Q-BamHI-D-R** | ATAGGATCCATGAATGCGCTCGAGC | BamHI |
| **ParB-NdeI-F** | CAGCATATGGCTCAGAACAAGGGTTCC | NdeI |
| **ParB-XhoI-R** | CAGCTCGAGTTATTGGCCCTGGATCAA | XhoI |
| **mCherry-SacI-F** | CAGGAGCTCATGGTGAGCAAGGGCGAG | SacI |
| **mCherry-EcoRI-R** | CATGAATTCTTACTTGTACAGCTCGTC | EcoRI |

| **Oligonucleotides qPCR** | **Sequence 5'–3'** | **Genomic binding region** |
|---|---|---|
| **parS1-F** | CAAGCTCATTCCAGCAGATG | cg3362 |
| **parS1-R** | AACGAGGAATGCATTGGAGT | cg3362 |
| **parS10-F** | CCGTTGAAGAACCAATGAGC | cg3394 |
| **parS10-R** | CGAGAACCAAGGAAAGGCTA | cg3394 |
| **control1-F** | TTGTTCGAGAGCTTGGGTTC | cg2088 5‘ |
| **control1-R** | TCGGGTCACCTGGACTTAAC | cg2088 5‘ |
| **control2-F** | TGATTCTGGAAGGGCTCCAT | cg1046 3‘ |
| **control2-R** | AGCGATTCGTGACGAGAAGT | cg1046 3‘ |
| **SMCload-F** | CGGTTCCGATGGAGTCACTT | cg3349 |
| **SMCload-R** | ATTCCCGCAAAGGAACATGG | cg3349 |
| **pBHK18-F** | CAGTGGGCTTACATGGCGATA | intergenic |
| **pBHK18-R** | AGGGCTTCCCAACCTTACCA | intergenic |
| **pWK0-F** | AATACGCAAACCGCCTCTCC | intergenic |
| **pWK0-R** | TAATTGCGTTGCGCTCACTG | intergenic |
| **pJC1-F** | CTTAACCGGCGCATGACTTC | intergenic |
| **pJC1-R** | TCTTGAGTCCAACCCGGAAA | intergenic |
| **pEC0-F** | TAATTGCGTTGCGCTCACTG | intergenic |
| **pEC0-R** | AATACGCAAACCGCCTCTCC | intergenic |

**Table S3. Bacterial strains and plasmids used in this study.**

| Strain | Characteristics | Reference |
|---|---|---|
| ***E. coli*** **DH5α** | F⁻ ϕ80*lacZ*ΔM15 (*lacZYA-argF*)*U169 recA1 endA1 hsdR17*($r_K^-$ $m_K^+$) *supE44 phoA thi-1 gyrA96 relA1* λ⁻ | Invitrogen |
| ***E. coli*** **BTH101** | F-, *cya*-99, *ara*D139, *gal*E15, *gal*K16, *rps*L1 ($Str^r$), *hsd*R2, *mcr*A1, *mcr*B1 strain | [2] |
| ***E. coli*** **BL21 pLysS** | F–, *omp*T, *hsd*$S_B$ ($r_B$–, $m_B$–), *dcm*, *gal*, λ(DE3), pLysS, $Cm^r$ | Thermo Fisher Scientific |
| ***B. subtilis 168*** | *trpC2* | Laboratory collection |
| ***C. glutamicum*** **RES 167** | Restriction-deficient mutant, otherwise considered wild type | [3] |
| **CDC003** | RES167 derivative, Δ*parB* | [4] |
| **CDC026** | RES167 derivative, Δ*smc* | This study |
| **CBK001** | RES167 derivative, Δ*mksB* | This study |
| **CBK002** | RES167 derivative, Δ*parB*, Δ*smc* | This study |
| **CBK003** | RES167 derivative, Δ*parB*, Δ*mksB* | This study |
| **CBK004** | RES167 derivative, Δ*smc*, Δ*mksB* | This study |
| **CBK005** | RES167 derivative, Δ*parB*, Δ*smc*, Δ*mksB* | This study |
| **CBK006** | RES167 derivative, *parB::parB-mCherry2* | [5] |
| **CBK007** | RES167 derivative, *parB::parB-eYFP* | [5] |
| **CBK008** | RES167 derivative, *parB::parB-mNeonGreen* | This study |
| **CBK009** | RES167 derivative, *parB::parB-PAmCherry* | This study |
| **CBK010** | RES167 derivative, Δ*smc*, *parB::parB-eYFP* | This study |
| **CBK011** | RES167 derivative, Δ*smc*, Δ*mksB parB::parB-eYFP* | This study |
| **CBK012** | RES167 derivative, *smc::smc-mCherry* | This study |
| **CBK013** | RES167 derivative, *smc::smc-mCherry*, *parB::parB-mNeonGreen* | This study |
| **CBK014** | RES167 derivative, *smc::smc-mCherry* Δ*parB* | This study |
| **CBK015** | RES167 derivative, *mksB:mksB-mCherry* | This study |
| **CBK016** | RES167 derivative, *parS 3* mutated | This study |
| **CBK017** | RES167 derivative, *parS 2-3* mutated | This study |
| **CBK018** | RES167 derivative, *parS 2-4* mutated | This study |
| **CBK019** | RES167 derivative, *parS 2-6* mutated | This study |
| **CBK020** | RES167 derivative, *parS 2-7* mutated | This study |
| **CBK021** | RES167 derivative, *parS 2-8* mutated | This study |
| **CBK022** | RES167 derivative, *parS 2-9* mutated | This study |
| **CBK023** | RES167 derivative, *parS 2-10* mutated | This study |
| **CBK024** | RES167 derivative, *parS 1-10* mutated | This study |
| **CBK025** | RES167 derivative, *parB::parB-eYFP*, *parS 2-10* mutated | This study |
| **CBK026** | RES167 derivative, *parB::parB-eYFP*, *parS 1-10* mutated | This study |
| **CBK027** | RES167 derivative, *parB::parB-mCherry2*, *parS 2-10* mutated | This study |

| | | |
|---|---|---|
| **CBK028** | RES167 derivative, *parB*∷*parB-mCherry2*, *parS 1-10* mutated | This study |
| **CBK029** | RES167 derivative, *parB*∷*parB-PAmCherry*, *parS 2-10* mutated | This study |
| **CBK030** | RES167 derivative, *parB*∷*parB-mCherry2*, *parS 2-9* mutated | This study |
| **CBK031** | RES167 derivative, *parB*∷*parB-PAmCherry*, *parS 2-9* mutated | This study |
| **CBK032** | RES167 derivative, *smc*∷*smc-mCherry*, *parS 1-10* mutated | This study |
| **CBK033** | RES167 derivative, *smc*∷*smc-mCherry*, partial *smc loading site* (1.1 Kb) deleted | This study |
| **CBK034** | RES167 derivative, *smc*∷*smc-mCherry*, partial *smc loading site* deleted and reinserted into intergenic region 3’ of *cg0177* | This study |
| **CBK035** | RES167 derivative, *smc*∷*smc-mCherry*, partial *smc loading site* (1.1 Kb) substituted by *B. subtilis* genomic locus of identical size | This study |
| **CBK036** | RES167 derivative, *parS 1-10* mutated, *parS* in intergenic region 3’ of *cg0108* | This study |
| **CBK037** | RES167 derivative, *parS 1-10* mutated, *parS* in intergenic region 3’ of *cg0904* | This study |
| **CBK038** | RES167 derivative, *parS 1-10* mutated, *parS* in intergenic region 3’ of *cg2563* | This study |
| **CBK039** | RES167 derivative, *parS 1-10* mutated, *cg1752*∷*parS* | This study |
| **CBK040** | RES167 derivative, *parB*∷*parB-eYFP*, *parS 1-10* mutated, *parS* in intergenic region 3’ of *cg0108* | This study |
| **CBK041** | RES167 derivative, *parB*∷*parB-eYFP*, *parS 1-10* mutated, *parS* in intergenic region 3’ of *cg0904* | This study |
| **CBK042** | RES167 derivative, *parB*∷*parB-mCherry2*, *parS 1-10* mutated, *parS* in intergenic region 3’ of *cg0904* | This study |
| **CBK043** | RES167 derivative, *parB*∷*parB-eYFP*, *parS 1-10* mutated, *parS* in intergenic region 3’ of *cg2563* | This study |
| **CBK044** | RES167 derivative, *parB*∷*parB-eYFP*, *parS 1-10* mutated, *cg1752*∷*parS* | This study |
| **CBK045** | RES167 derivative, *smc*∷*smc-mCherry*, *parS 1-10* mutated, *parS* in intergenic region 3’ of *cg0904* | This study |
| **CBK046** | RES167 derivative, *parS 1-10* mutated, *parS* in intergenic region 3’ of *cg0904*, Δ*smc* | This study |
| **CBK047** | RES167 derivative, *parB*∷*parB*$^{R175A}$*-mCherry2* | This study |
| **CBK048** | RES167 derivative, *parB*∷*parB*$^{R175A}$*-mCherry2*, *parS 2-10* mutated | This study |
| **CBK049** | RES167 derivative, *parB*∷*parB*$^{R175A}$, *smc*∷*smc-mCherry* | This study |
| **CBK050** | RES167 derivative, *smc*∷*smc*$^{E1084Q}$ | This study |
| **CBK051** | RES167 derivative, *smc*∷*smc*$^{E1084Q}$*-mCherry* | This study |
| **CBK052** | RES167 derivative, pEKEx2-mCherry | This study |
| **CBK053** | RES167 derivative, pBHK18 | This study |
| **CBK054** | RES167 derivative, pWK0 | This study |

| | | |
|---|---|---|
| **CBK055** | RES167 derivative, pJC1 | This study |
| **CBK056** | RES167 derivative, pEK0 | This study |
| **CBK057** | RES167 derivative, Δ*mksB*, pBHK18 | This study |
| **CBK058** | RES167 derivative, Δ*mksB*, pWK0 | This study |
| **CBK059** | RES167 derivative, Δ*mksB*, pJC1 | This study |
| **CBK060** | RES167 derivative, Δ*mksB*, pEK0 | This study |
| **Plasmid** | **Characteristics** | **Reference** |
| **pUT18** | Cloning/expression vector, pUC19 derivative, T18 domain of CyaA, MCS 5' of T18, Amp$^r$ | [6] |
| **pUT18C** | Cloning/expression vector, pUC19 derivative, T18 domain of CyaA, MCS 3' of T18, Amp$^r$ | [6] |
| **pKNT25** | Cloning/expression vector, pSU40 derivative, T25 domain of CyaA, MCS 5' of T25, Kan$^r$ | [7] |
| **pKT25** | Cloning/expression vector, pSU40 derivative, T25 domain of CyaA, MCS 3' of T25, Kan$^r$ | [6] |
| **pUT18C-zip** | Control plasmid, T18 domain of CyaA fused to leucine zipper of GCN4, Amp$^r$ | [6] |
| **pKT25-zip** | Control plasmid, T25 domain of CyaA fused to leucine zipper of GCN4, Kan$^r$ | [6] |
| **pUT18_mcs** | pUT18 plasmid, insertion (TAATGG) between XmaI and KpnI restriction sites | This study |
| **pUT18C_mcs** | pUT18C plasmid, insertion (TAATGG) between XmaI and KpnI restriction sites | This study |
| **pKNT25_mcs** | pKNT25 plasmid, insertion (TAATGG) between XmaI and KpnI restriction sites | This study |
| **pKT25_mcs** | pKT25 plasmid, insertion (TAATGG) between XmaI and KpnI restriction sites | This study |
| **pUT18-divIVA** | pUT18 plasmid, *divIVA-cyaAT18* fusion | This study |
| **pUT18C-divIVA** | pUT18C plasmid, *cyaAT18-divIVA* fusion | This study |
| **pKNT25-divIVA** | pKNT25 plasmid, *divIVA-cyaAT25* fusion | This study |
| **pKT25-divIVA** | pKT25 plasmid, *cyaAT25-divIVA* fusion | This study |
| **pUT18-parB** | pUT18 plasmid, *parB-cyaAT18* fusion | This study |
| **pUT18C-parB** | pUT18C plasmid, *cyaAT18-parB* fusion | This study |
| **pKNT25-parB** | pKNT25 plasmid, *parB-cyaAT25* fusion | This study |
| **pKT25-parB** | pKT25 plasmid, *cyaAT25-parB* fusion | This study |
| **pUT18-parBR175A** | pUT18 plasmid, *parB*$^{R175A}$-*cyaAT18* fusion | This study |
| **pUT18C-parBR175A** | pUT18C plasmid, *cyaAT18-parB*$^{R175A}$ fusion | This study |
| **pKNT25-parBR175A** | pKNT25 plasmid, *parB*$^{R175A}$-*cyaAT25* fusion | This study |
| **pKT25-parBR175A** | pKT25 plasmid, *cyaAT25-parB*$^{R175A}$ fusion | This study |
| **pUT18-smc** | pUT18 plasmid, *smc-cyaAT18* fusion | This study |
| **pUT18C-smc** | pUT18C plasmid, *cyaAT18-smc* fusion | This study |
| **pKNT25-smc** | pKNT25 plasmid, *smc-cyaAT25* fusion | This study |
| **pKT25-smc** | pKT25 plasmid, *cyaAT25-smc* fusion | This study |
| **pUT18-scpA** | pUT18 plasmid, *scpA-cyaAT18* fusion | This study |
| **pUT18C-scpA** | pUT18C plasmid, *cyaAT18-scpA* fusion | This study |
| **pKNT25-scpA** | pKNT25 plasmid, *scpA-cyaAT25* fusion | This study |

| | | |
|---|---|---|
| **pKT25-scpA** | pKT25 plasmid, *cyaAT25-scpA* fusion | This study |
| **pUT18-scpB** | pUT18 plasmid, *scpB-cyaAT18* fusion | This study |
| **pUT18C-scpB** | pUT18C plasmid, *cyaAT18-scpB* fusion | This study |
| **pKNT25-scpB** | pKNT25 plasmid, *scpB-cyaAT25* fusion | This study |
| **pKT25-scpB** | pKT25 plasmid, *cyaAT25-scpB* fusion | This study |
| **pUT18-mksB** | pUT18_mcs plasmid, *mksB-cyaAT18* fusion | This study |
| **pUT18C-mksB** | pUT18C_mcs plasmid, *cyaAT18-mksB* fusion | This study |
| **pKNT25-mksB** | pKNT25_mcs plasmid, *mksB-cyaAT25* fusion | This study |
| **pKT25-mksB** | pKT25_mcs plasmid, *cyaAT25-mksB* fusion | This study |
| **pUT18-mksE** | pUT18 plasmid, *mksE-cyaAT18* fusion | This study |
| **pUT18C-mksE** | pUT18C plasmid, *cyaAT18-mksE* fusion | This study |
| **pKNT25-mksE** | pKNT25 plasmid, *mksE-cyaAT25* fusion | This study |
| **pKT25-mksE** | pKT25 plasmid, *cyaAT25-mksE* fusion | This study |
| **pUT18-mksF** | pUT18 plasmid, *mksF-cyaAT18* fusion | This study |
| **pUT18C-mksF** | pUT18C plasmid, *cyaAT18-mksF* fusion | This study |
| **pKNT25-mksF** | pKNT25 plasmid, *mksF-cyaAT25* fusion | This study |
| **pKT25-mksF** | pKT25 plasmid, *cyaAT25-mksF* fusion | This study |
| **pUT18-mksG** | pUT18 plasmid, *mksG-cyaAT18* fusion | This study |
| **pUT18C-mksG** | pUT18C plasmid, *cyaAT18-mksG* fusion | This study |
| **pKNT25-mksG** | pKNT25 plasmid, *mksG-cyaAT25* fusion | This study |
| **pKT25-mksG** | pKT25 plasmid, *cyaAT25-mksG* fusion | This study |
| **pK19mobsacB** | Integration vector, *ori* pUC, $Km^r$, *mob sac* | [8] |
| **pK19mobsacB-Δsmc** | Integration vector, *ori* pUC, $Km^r$, *mob sac,* deletion of *smc* | This study |
| **pK19mobsacB-ΔmksB** | Integration vector, *ori* pUC, $Km^r$, *mob sac,* deletion of *mksB* | This study |
| **pK19mobsacB-ΔparB** | Integration vector, *ori* pUC, $Km^r$, *mob sac,* deletion of *parB* | [4] |
| **pK19mobsacB-parB-eYFP** | Integration vector, *ori* pUC, $Km^r$, *mob sac, parB-eYFP* | [5] |
| **pK19mobsacB-parB-mCherry2** | Integration vector, *ori* pUC, $Km^r$, *mob sac, parB-mCherry2* | [5] |
| **pK19mobsacB-parB-mNeonGreen** | Integration vector, *ori* pUC, $Km^r$, *mob sac, parB-mCherry2* | This study |
| **pK19mobsacB-parB-PAmCherry** | Integration vector, *ori* pUC, $Km^r$, *mob sac, parB-PAmCherry2* | This study |
| **pK19mobsacB-smc-mCherry** | Integration vector, *ori* pUC, $Km^r$, *mob sac, smc-mCherry* | This study |
| **pK19mobsacB-smc-PAmCherry** | Integration vector, *ori* pUC, $Km^r$, *mob sac, smc-PAmCherry* | This study |
| **pK19mobsacB-mksB-mCherry** | Integration vector, *ori* pUC, $Km^r$, *mob sac, mksB-mCherry* | This study |
| **pK19mobsacB-ΔSMCload** | Integration vector, *ori* pUC, $Km^r$, *mob sac,* deletion of SMC binding site (1.1 Kb *hpaG* 3’) | This study |

| | | |
|---|---|---|
| **pK19mobsacB-SMCload-cg0177** | Integration vector, *ori* pUC, $Km^r$, *mob sac,* partial SMC binding site (1.1 Kb *hpaG* 3') *cg0177* 3' | This study |
| **pK19mobsacB-SMCload-r** | Integration vector, *ori* pUC, $Km^r$, *mob sac,* partial replacement of SMC binding site (1.1 Kb *hpaG* 3') by *B. subtilis* genomic region | This study |
| **pK19mobsacB-parS1mut** | Integration vector, *ori* pUC, $Km^r$, *mob sac,* point mutations in *parS1* | This study |
| **pK19mobsacB-parS2mut** | Integration vector, *ori* pUC, $Km^r$, *mob sac,* point mutations in *parS2* | This study |
| **pK19mobsacB-parS3mut** | Integration vector, *ori* pUC, $Km^r$, *mob sac,* point mutations in *parS3* | This study |
| **pK19mobsacB-parS4mut** | Integration vector, *ori* pUC, $Km^r$, *mob sac,* point mutations in *parS4* | This study |
| **pK19mobsacB-parS5_6mut** | Integration vector, *ori* pUC, $Km^r$, *mob sac,* point mutations in *parS5* and *parS6* | This study |
| **pK19mobsacB-parS7mut** | Integration vector, *ori* pUC, $Km^r$, *mob sac,* point mutations in *parS7* | This study |
| **pK19mobsacB-parS8mut** | Integration vector, *ori* pUC, $Km^r$, *mob sac,* point mutations in *parS8* | This study |
| **pK19mobsacB-parS9_10mut** | Integration vector, *ori* pUC, $Km^r$, *mob sac,* point mutations in *parS9* and *parS10* | This study |
| **pK19mobsacB-parS-cg0108** | Integration vector, *ori* pUC, $Km^r$, *mob sac,* *parS cg0108* 3' | This study |
| **pK19mobsacB-parS-cg0904** | Integration vector, *ori* pUC, $Km^r$, *mob sac,* *parS cg0904* 3' | This study |
| **pK19mobsacB-parS-cg02563** | Integration vector, *ori* pUC, $Km^r$, *mob sac,* *parS cg2564* 3' | This study |
| **pK19mobsacB-parS-Δint** | Integration vector, *ori* pUC, $Km^r$, *mob sac,* replacement of *cg1752* by *parS* | This study |
| **pK19mobsacB-parBR175A** | Integration vector, *ori* pUC, $Km^r$, *mob sac,* *parB*∷*parB*$^{R175A}$ | This study |
| **pK19mobsacB-smcE1084Q** | Integration vector, *ori* pUC, $Km^r$, *mob sac,* *smc*∷*smc*$^{E1084Q}$ | This study |
| **pET-16b** | *E. coli* protein expression vector, $p_{T7\text{-}lac}$, $Amp^R$, N-10xHis tag, pBR322 | Novagen |
| **pET-16b-ParB** | pET-16b, *parB* | This study |
| **pET-16b-ParBR175A** | pET-16b, *parB*$^{R175A}$ | This study |
| **pEKEx2** | *E. coli-C. glutamicum* shuttle expression vector, $P_{tac}$, $lacI_q$, $Km^R$, pBL1 *oriV*$_{C.g.}$, pUC18 *oriV*$_{E.c.}$ | [9] |
| **pEKEx2-mCherry** | pEKEx2, *mCherry* | This study |
| **pBHK18** | *E. coli-C. glutamicum* shuttle vector, $Km^R$, pNG2 *oriV*$_{C.g.}$, low copy number | [10] |
| **pWK0** | *E. coli-C. glutamicum* shuttle vector, $Km^R$, pNG2 *oriV*$_{C.g.}$, low copy number | [11] |
| **pJC1** | *E. coli-C. glutamicum* shuttle vector, $Km^R$, pCG1 *oriV*$_{C.g}$. | [12] |
| **pEC0** | *E. coli-C. glutamicum* shuttle vector, $Km^R$, pBL1 *oriV*$_{C.g}$. | [9] |

#### Supplementary Methods

##### Strain construction

The following section provides detailed information on bacterial strain construction and plasmids used for this purpose.

For protein-protein interaction screens genes of interest were amplified via PCR, digested with respective enzymes and ligated into bacterial two-hybrid vectors [2]. *E. coli* DH5α were utilized for plasmid cloning. Genes *divIVA* and *parB*/*parB*$^{R175A}$ were amplified using primer pairs DivIVA-XbaI-F/ DivIVA-BamHI-R and ParB-XbaI-F/ ParB-BamHI-R from genomic DNA or pK19mobsacB-ParBR175A and resulting fragments were digested with XbaI/ BamHI. For amplification of *scpA*, *scpB*, *mksE*, *mksF* and *mksG* primer pairs ScpA-XbaI-F/ ScpA-XmaI-R, ScpB-XbaI-F/ ScpB-XmaI-R, MksE-XbaI-F/ MksE-XmaI-R, MksF-XbaI-F/ MksF-XmaI-R and MksG-XbaI-F/ MksG-XmaI-R were utilized, followed by restriction digests with XmaI/ XbaI. Primer pairs SMC-XbaI-F/ SMC-KpnI-R and MksB-XmaI-F/ MksB-KpnI-R were used for PCR amplification of genes *smc* and *mksB*, which were subsequently digested with XbaI/ KpnI or XmaI/ KpnI. In order to increase the distance of XmaI and KpnI restriction sites a short sequence was inserted in between these sites by overhang PCRs using pUT18C-mcs-HindIII-F, pUT18-mcs-PvuII-F, pKNT25-mcs-NheI-F or pKT25-mcs-HindIII-F in combination with pUT18(C)/pK(N)T25-mcs-KpnI-R for plasmids pUT18C, pUT18, pKT25 and pKNT25, respectively. Resulting fragments and corresponding vectors were digested with HindIII/ KpnI, PvuII/ KpnI or NheI/ KpnI and subsequently ligated, resulting in plasmids pUT18_mcs, pUT18C_mcs, pKNT25_mcs and pKT25_mcs. All digested gene fragments mentioned above were ligated into pUT18, pUT18C, pKNT25 and pKT25 or pUT18_mcs, pUT18C_mcs, pKNT25_mcs and pKT25_mcs, respectively.

Derivatives of the suicide integration vector pK19mobsacB were used for clean allelic replacements in *C. glutamicum*, containing the modified genomic region of interest including its 500 bp up- and downstream homologous flanking sequences. Plasmid cloning was performed using *E. coli* DH5α.

In order to construct pK19mobsacB-Δsmc 500 bp upstream and downstream of *smc* were PCR-amplified using primer pairs Δsmc-BamHI-up-F/ Δsmc-up-R and Δsmc-D-F/ Δsmc-EcoRI-D-R, respectively. Both fragments served as templates in an overhang PCR, yielding a 1000 bp fragment, which was digested with BamHI and EcoRI and subsequently ligated into pK19mobsacB. pK19mobsacB-ΔSMCload was constructed accordingly, using primer pairs ΔSMCload-HindIII-up-F/ ΔSMCload-up-R and ΔSMCload-D-F/ ΔSMCload-SalI-D-R and HindIII in combination with SalI for restriction digest. For construction of pK19mobsacB-ΔmksB up-/and downstream regions of *mksB* were PCR amplified using primers ΔmksB-HindIII-up-F/ ΔmksB-PstI-up-R and ΔmksB-PstI-D-F/ ΔmksB-XbaI-D-R. Resulting 500 bp fragments were digested with HindII/ PstI and PstI/ XbaI and consecutively ligated into pK19mobsacB.

Fluorescent C-terminal fusions of ParB protein with PAmCherry or mNeongreen were obtained by utilizing plasmids pK19mobsacB-parB-mNeonGreen and pK19mobsacB-parB-PAmCherry. To this end, the *eYFP* sequence of plasmid pK19mobsacB-parB-eYFP (Böhm et al) was replaced by respective fluorophore sequences, which were amplified via PCR using PAmCherry-SalI-F/ PAmCherry-XbaI-R primers and digested with SalI and XbaI.

For fluorescent versions of SMC and MksB proteins plasmids pK19mobsacB-smc-mCherry and pK19mobsacB-mksB-mCherry were constructed. At first, 500 bp regions up- and downstream of the 3' end of *smc* or *mksB* were amplified using primer pairs SMC-HindIII-up-F/ SMC-SphI-up-R and SMC-BamHI-D-F/ SMC-EcoRI-D-R or MksB-

HindIII-up-F/ MksB-Sph-up-R and MksB-BamHI-D-F/ MksB-EcoRI-D-R. Fluorophore sequences were amplified with primers mCherry-XbaI-F/ mCherry-BamHI-R for SMC-mCherry fusion or with primers PAmCherry-SalI-F/ mCherry-XbaI-R for the MksB-mCherry fusion construct. Up- and downstream fragments were digested via HindIII/SphI and BamHI/ EcoRI, while enzymes SalI/ BamHI or SalI/ XbaI were utilized for restriction digest of fluorophore sequences fused to *smc* or *mksB*, respectively. Fragments were subsequently ligated into the pK19mobsacB plasmid, starting with the corresponding downstream region, followed by the fluorophore sequence and finally the upstream region.

In order to place part of a putative SMC binding site upstream of the *parS* cluster into an intergenic region 3' of cg0177 (Fig. S9), genomic sequences 500 bp up-and downstream of the insertion site were amplified using primer pairs cg0177-HindIII-up-F/ cg0177-SalI-up-R and cg0177-XmaI-D-F/ cg0177-EcoRI-D-R; part of the genomic SMC binding site (1.1 Kb) was amplified using primers SMCload-SalI-F and SMCload-XmaI-R. Resulting fragments were digested with HindII/ SalI, SalI/ XmaI and XmaI/ EcoRI and consecutively ligated into pK19mobsacB, obtaining the plasmid pK19mobsacB-SMCload-cg0177.

Plasmid pK19mobsacB-SMCload-r was constructed for the partial replacement of the SMC binding site (1.1 Kb) with a *B. subtilis* genomic region of identical size. For amplification of up- and downstream 500 bp regions primer pairs ΔSMCload-HindIII-up-F/ SMCload-SphI-up-R and SMCload-PstI-D-F/ ΔSMCload-SalI-D-R were utilized, while the replacement sequence was amplified from *B. subtilis* genomic DNA via SMCloadr-SphI-F/ SMCloadr-PstI-R. After digestion with enzymes HindIII/ SphI, PstI/ SalI or SphI/ PstI fragments were successively ligated into pK19mobsacB.

Further, all *parS* sites were mutated comprising new XmaI or SalI restriction sites (See Fig. S2). For mutation of *parS1* primer pairs parS1mut-HindIII-up-F/ parS1mut-XmaI-up-R and parS1mut-XmaI-D-F/ parS1mut-EcoRI-D-R were utilized to mutate *parS1* and to amplify sequences 500 bp up- and downstream of *parS1*. Restriction digest was performed with both fragments using HindIII/ XmaI or XmaI/ EcoRI, respectively. Subsequent ligation into pK19mobsacB yielded plasmid pK19mobsacB-parS1mut. In order to mutate *parS2*, *parS3*, *parS4*, *parS7* and *parS8* plasmid construction was performed in the same way using primers parS2mut-HindIII-up-F/ parS2mut-XmaI-up-R and parS2mut-XmaI-D-F/ parS2mut-EcoRI-D-R, parS3mut-HindIII-up-F/ parS3mut-XmaI-up-R and parS3mut-XmaI-D-F/ parS3mut-EcoRI-D-R, parS4mut-HindIII-up-F/ parS4mut-XmaI-up-R and parS4mut-XmaI-D-F/ parS4mut-EcoRI-D-R, parS7mut-HindIII-up-F/ parS7mut-XmaI-up-R and parS7mut-XmaI-D-F/ parS7mut-EcoRI-D-R or parS8mut-HindIII-up-F/ parS8mut-XmaI-up-R and parS8mut-XmaI-D-F/ parS8mut-EcoRI-D-R for amplification of fragments up- and downstream of the respective *parS* site. Matching fragments were each digested and ligated into pK19mobsacB, as exemplified for pK19mobsacB-parS1mut construction, resulting in plasmids pK19mobsacB-parS2mut, pK19mobsacB-parS3mut, pK19mobsacB-parS4mut, pK19mobsacB-parS7mut and pK19mobsacB-parS8mut.

Since *parS5* and *parS6* as well as *parS9* and *parS10* are localized in close proximity on the genome (<100 bp distance), their deletions were accomplished using in each case one plasmid for both *parS* sites. For construction of pK19mobsacB-parS5_6mut genomic region upstream of *parS5*, downstream of *parS6* and in between both sides were PCR-amplified using parS5mut-HindIII-up-F/ parS5mut-SalI-up-R, parS6mut-XmaI-D-F/ parS6mut-EcoRI-D-R and parS5mut-SalI-D-F/ parS6mut-XmaI-up-R and fragments were digested with HindIII/ SalI, XmaI/ EcoRI or SalI/ XmaI, respectively and ligated into pK19mobsacB. Construction of pK19mobsacB-parS9_10mut was performed accordingly, using primer pairs parS9mut-HindIII-up-F/ parS9mut-SalI-up-R, parS10mut-XmaI-D-F/ parS10mut-EcoRI-D-R and parS9mut-SalI-D-F/ parS10mut-XmaI-up-R for fragment amplification.

Insertion of *parS* 3' of *cg0108*, *cg0904* and *cg2563* were achieved via plasmids pK19mobsacB-parS-cg0108, pK19mobsacB-parS-cg0904 and pK19mobsacB-parS-cg02563. Primers containing *parS* sites were used to amplify regions 500 bp up- and downstream of the corresponding *parS* insertion site, namely parS-cg0108-SalI-up-F/ parS-cg0108-up-R and parS-cg0108-D-F/ parS-cg0108-XmaI-D-R, parS-cg0904-HindIII-up-F/ parS-cg0904-up-R and parS-cg0904-D-F/ parS-cg0904-NheI-D-R or parS-cg2563-HindIII-up-F/ parS-cg2563-up-R and parS-cg2563-D-F/ parS-cg2563-NheI-D-R, respectively. Each fragment pair served as template in an overhang PCR, yielding 1000 bp sequences with central *parS* sites. After restriction digest with SalI/ XmaI or HindIII/NheI each fragment was ligated into pK19mobsacB. Plasmid pK19mobsacB-parS-Δint for *parS* insertion at *terC* was constructed in the same way, however by replacing an entire gene (*cg1752*). Regions 500 bp N- and C-terminally of *cg1752* were amplified using parS-Δint-HindIII-up-F/ parS-Δint-up-R and parS-Δint-D-F/ parS-Δint-NheI-D-R.

For construction of pK19mobsacB-parBR175A primer pairs ParB-N-ter-HindIII-F/ ParB-R175A-R and ParB-R175A-F/ ParB-C-ter-SalI-R were used to amplify the N-and C-terminal parts of *parB* surrounding the coding region of $ParB^{R175}$. Primers introduce point mutations into this codon as well as into a neighboring SacI restriction site, resulting in fragments of 528 bp and 625 bp length. Overhang PCR yielded a full *parB* sequence that was cut with HindIII/ SalI and ligated into pK19mobsacB. pK19mobsacB-smcE1084Q was obtained in an analogous manner. Amplification of 500 bp genomic regions surrounding codon $SMC^{E1084}$ were performed using primer pairs E1084Q-HindIII-up-F/ E1084Q-up-R and E1084Q-D-F/ E1084Q-BamHI-D-R, which further yield in an E1084Q mutation and an additional XbaI restriction site 3' of the codon sequence.

His-tagged versions of ParB and ParB$^{R175A}$ were generated by applying PCR (ParB-NdeI-F/ ParB-XhoI-R) following a restriction digest (NdeI/ XhoI) of the respective DNA fragment and ligation into pET-16b expression vector yielding pET-16b-ParB and pET-16b-ParBR175A.

For construction of the *E. coli* -*C. glutamicum* shuttle expression vector pEKEx2-mCherry the mCherry sequence was amplified via PCR using mCherry-SacI-F/ mCherry-EcoRI-R, digested with corresponding restriction enzymes and ligated into the empty pEKEx2.

Vectors were transformed via electroporation into *C. glutamicum* cells [8]. Genomic integration of pK19mobsacB plasmids were selected on kanamycin, while the second crossover event was confirmed by growth on 10% sucrose. Screening of allelic replacements in *C. glutamicum* Δ*smc*, Δ*mksB* and Δ*parB* was performed by colony PCR using primer pairs Δsmc-seq-700up-F/ Δsmc-seq-700D-R, ΔmksB-seq-700up-F/ ΔmksB-seq-700D-R and ParB-seq-800up-F/ ParB-seq-800D-R. Fluorescent fusions of ParB, SMC and MksB were confirmed via primer pairs ParB-N-ter-SalI-F/ ParB-seq-800D-R, SMC-seq-1589bp-F/ Δsmc-seq-700D-R and MksB-seq-1595bp-F/ ΔmksB-seq-700D-R, respectively. Insertions of the partial *smc loading site* in an intergenic region 3' of *cg0177* were screened using primer pairs cg0177-seq-700up-F/ cg0177-seq-700D-R. In order to identify genomic *parS* mutations respective regions were amplified using upstream-forward and downstream-reverse primers as used for plasmid construction and digested with either XmaI or SalI. Sequencing of *parS* loci was performed for further verification. For verification of *parS* insertions 3' of *cg0108*, *cg0904* and *cg2563* or for replacement of *cg1752* by *parS* genomic loci were amplified with primers cg0108-seq-400up-F/ cg0108-seq-200D-R, cg0904-seq-100up-F/ cg0904-seq-100D-R, cg2563-seq-200up-F/ cg2563-seq-300D-R or Δint-seq-700up-F/ Δint-seq-700D-R, respectively, followed by a control restriction digest using PmlI. Screening for *parB*$^{R175A}$ was performed by amplification of *parB* including 800 bp up- and downstream regions via primers ParB-seq-800up-F/ ParB-seq-800D-R. A control digest was conducted with the resulting fragment using SacI. Integration of the point mutation smc$^{E1084Q}$ was verified by amplification of the respective genomic region (E1084Q-HindIII-up-F/ mCherry-EcoRI-R), followed by restriction digests using XbaI.

Assembly strategies of multiple consecutive allelic replacements are explained hereafter. *C. glutamicum* strains CBK002, CBK004 and CBK010 were obtained via transformation of pK19mobsacB-ΔparB, pK19mobsacB-ΔmksB or pK19mobsacB-parB-eYFP into strain CDC026 lacking *smc* and strain CBK003 (Δ*mksB* Δ*parB*) was constructed using the genetic background of CBK001 (Δ*mksB*). Further, CBK004 served as parent strain for construction of CBK005 and CBK011 harboring additional mutations Δ*parB* and *parB::parB-eYFP*, respectively. The dual-reporter strain CBK013, expressing ParB-mNeonGreen in combination with SMC-mCherry, was constructed via transformations of pK19mobsacB-smc-mCherry into CBK008; strain CBK014 derives from CBK012 transformed with pK19mobsacB-ΔparB. The complete loss of *parS* sites in strain CBK024 was accomplished via successive allelic replacements of *parS* by mutated sequences: the mutation of *parS2* (CBK017) followed the mutation of *parS3* (CBK016); thereupon *parS4* (CBK018) was mutated followed by *parS5* and *parS6* (CBK019). Next, *parS7* (CBK020) mutation, *parS8* mutation (CBK021), *parS9* mutation (CBK022), *parS10* mutation (CBK023) and *parS1* mutation (CBK024) were accomplished consecutively. CBK025, CBK027 and CBK029 derive from strain CBK023, which was transformed with pK19mobsacB plasmids coding for *parB-eYFP*, *parB-mCherry2*, *parB-PAmCherry*, respectively. Accordingly, stains CBK026, CBK28, and CBK032 are CBK024-derivatives harboring either endogenous *parB-eYFP*, *parB-mCherry2* or *smc-mCherry*, while strains CBK030 and CBK031 obtained from CBK022 via transformation of pK19mobsacB-parB-mCherry2 or pK19mobsacB-parB-PAmCherry. CBK033 and CBK035 were generated by transformation of strain CBK012 expressing SMC-mCherry with plasmids pK19mobsacB-ΔSMCload or pK19mobsacB-SMCload-r; a further transformation of CBK033 with pK19mobsacB-SMCload-cg0177 yielded CBK034. In order to introduce *parS* sites at different regions within the *C. glutamicum* genome CBK024 served as parental strain: *parS*-insertions at chromosomal 9.5°, 90°, 270° and 180° positions were achieved via transformation of either pK19mobsacB-parS-cg0108 (CBK036), pK19mobsacB-parS-cg0904 (CBK037), pK19mobsacB-parS-cg2563 (CBK038) or pK19mobsacB-parS-Δint (CBK039). Additional allelic replacements of *parB* or *smc* with fluorophore-coupled versions *parB-eYFP* or *parB-mCherry2* and *smc-mCherry* in above-named strains resulted in CBK040-CBK045. Secondly, *parS*-insertion in CBK037 was combined with a *smc* deletion by transformation of pK19mobsacB-Δsmc yielding CBK046. Lastly, strains CBK047-CBK051 which express mutant ParB$^{R175A}$ or SMC$^{E1084Q}$ proteins derive from CBK006, CBK027 and CBK012 transformed with plasmid pK19mobsacB-parBR175A or pK19mobsacB-smcE1084Q, respectively.

**Growth conditions and media**

*E. coli* cells were grown at 37°C in Lysogeny Broth (LB) medium supplemented with 50 µg/ml Kanamycin when appropriate. Growth experiments of *C. glutamicum* cells were performed using brain heart infusion medium (BHI, Oxoid™) or CGXII medium [13] supplemented with 4% glucose or 120 mM acetate at 30°C. Cells were always preinoculated in BHI overnight; for growth in minimal media cells were first inoculated in BHI and rediluted in the corresponding growth media overnight for pre-cultivation. Finally, cell cultures were adjusted to an $OD_{600}$ of 0.5 for BHI and to an $OD_{600}$ of 1 for growth in CGXII medium. 25 µg/ml Kanamycin was added where applicable.

**Protein identification via immunoprecipitation and mass spectrometry**

For immunoprecipitation of interacting proteins strains CBK012, CBK015 and CBK052 were cultivated in BHI medium using culture flasks pretreated with 0,5 % sodium hypochlorite. CBK052 was induced at $OD_{600}$ ~ 1 with 0,5 mM IPTG. Exponentially growing cells ($OD_{600}$=3, 10 ml) were harvested, washed once in 10 ml washing buffer (Tris-HCl pH 7,5 10 mM; NaCl 150 mM; EDTA 0,5mM) and resuspended in 1,5 ml washing buffer supplemented with 1 mM PMSF in EtOH. All following steps were performed at 4°C. After cell disruption via FastPrep®-24 (MP Biomedicals) at 10 x 6,5 m/ s 30 sec cell debris was removed by centrifugation at 18000 g. Immunoprecipitation was performed with 25 µl magnetic RFP-Trap® agarose beads (Chromotek) incubated in 1 ml Lysate for 1 h. Thereupon, beads were washed three times in washing buffer and again washed three times in 100 mM ammonium bicarbonate prior to storage at -20°C.

For proteomic analysis of interacting proteins, the magnetic beads were first washed with 50 µl of 100 mM TRIS, pH 7.6. Subsequently, 50 µl of 100 mM TRIS, pH 7.6 containing 4 M urea, 5 mM dithiothreitol for reduction of disulfide bond and 0.2 µg of LysC for predigestion of proteins were added to each sample. After incubation of 3h, 100 µl of 100 mM TRIS, pH 7.6, 10 mM iodoacetamide was added for blocking of free cysteine side chains and samples were incubated in the dark for 5 min. Samples were diluted with 100 µl TRIS, pH 7.6 to reduce the urea concentration and 1 µg of trypsin was added to each sample. The samples were incubated for 14 h to complete protein digestion and subsequently trifluoroacetic acid was added to a final concentration of

0.5 % to acidify the samples. Peptide mixture were separated from the magnetic beads before the desalting step. The beads were washed 2x with 75 µl of 0.1% formic acid (FA) and the wash solvent was combined with the peptide mixtures. For sample desalting, 3 discs were stamped from C18 discs (Empore C18, 3M) and placed into a 200 µl pipette tip. Following binding of peptides, stage tips were washed 2x with 60 µl of 0.1% FA and peptides were eluted with 40% acetonitrile containing 30% methanol and 0.1% FA. Samples were dried in a speedvac and resuspended in 10 µl of 0.1% FA. Peptide mixtures were analyzed by liquid chromatography tandem mass spectrometry (LC-MS/MS) to identify and quantify proteins in all samples. First, peptides were separated by nano-reversed phase chromatography using a linear gradient from 2 to 35% acetonitrile over 50 min in 0.1% formic acid on an in house-packed chromatography column in a nano-electrospray emitter tip. Eluting peptides were directly infused into the mass spectrometer (QExactive, Thermo-Fisher) and detected in positive ionization mode. The operating cycle was programmed to detect peptides in the range from 300 to 1600 m/z and up to 10 precursors were selected for MSMS analysis by CID fragmentation. Precursor ions required a charge state between +2 and +6 and a minimal signal intensity of 6x10e4.

Protein mapping and quantitative analysis raw LC-MS/MS data were searched against a *C. glutamicum* database retrieved from Uniprot (vs. 03/2017, 3093 protein entries) using a forward/reversed search by the Andromeda algorithm within the MaxQuant software suite. Peptides hits were searched with 17 ppm precursor mass deviation in the first search and 3 ppm for the main search. For MS/MS spectra, a mass accuracy of 25 ppm was set. As variable modifications, acetylation of the protein N-terminus, STY-phosphorylation, and methionine oxidation were selected. Carbamidomethylation of cysteine was the only fixed modification. Peptide match results were sorted by their probability score and filtered for 2% reversed peptide hits and 5% reversed protein hits.

To calculate protein enrichments and significance values, reversed protein hits and proteins with less than 3 quantitative values in any of the three sample types (control, mksB IP and smc IP) were filtered out. The iBAQ-values were log2 transformed and median normalized. In case of one missing value in the triplicate measurements the value was imputed using a closest neighbor method, for more missing data points a random value from a standard distribution downshifted by a factor of 1.8 from the sample distribution and width of 0.3 was selected. Samples were compared using a students t-test which was FDR rate controlled by sample permutation.

**Fluorescence microscopy**

Fluorescence microscopy was performed with exponentially grown cells mounted on agarose coated slides (1% agarose). Images were acquired on an Axio-Imager M1 fluorescence microscope (Carl Zeiss) with an EC Plan Neofluar 100x/ 1.3 oil Ph3 objective and a 2.5 x optovar. Fluorescence of protein fusions with eYFP and mCherry/mCherry2 or DNA stained via Hoechst 33342 (1 µg/ml, Thermo Scientific) were detected using filter sets 46 HE YFP (EX BP 500/25, BS FT 515, EM BP 535/30), 43 HE Cy 3 shift free (EX BP 550/25, BS FT 570, EM BP 605/70) and 49 DAPI shift free (EX G 365, BS FT 395, EM BP 445/50). Life cell imaging as well as detection of fluorescently labeled condensin subunits were carried out using a Delta Vision Elite microscope (GE Healthcare, Applied Precision) with a standard four color InsightSSI module, a 100×/ 1.4 oil PSF U-Plan S-Apo objective and the YFP (EX BP 513/17, EM BP 548/22) and mCherry (EX BP 575/25, EM BP 625/45) specific filter sets. In order to conduct time-lapse experiments exponentially grown cells were diluted to an $OD_{600}$ of 0.01 in BHI and loaded in a microfluidic chamber (B04A CellASIC®, Onix); the environmental chamber was heated to 30°C and 0.75 psi were applied for nutrient supply throughout the experiment. Images were taken in 5 min intervals.

**Detailed description of chromatin immunoprecipitation**

In vivo ChIP experiments with *C. glutamicum* ParB, SMC or MksB proteins were conducted using strains with allelic replacements of respective proteins with mCherry-tagged versions. Exponentially growing cells were crosslinked in 1 % formaldehyde for 30 min at room temperature; for SMC- and MksB-mCherry ChIP experiments cells were treated with Crosslink Gold (Diagenode) for 30 min at room temperature and washed twice in phosphate-buffered saline (PBS; 137 mM NaCl, 10 mM $Na_2HPO_4$, 1.8 mM $KH_2PO_4$, 2.7 mM KCl, pH 7.4.) prior to formaldehyde crosslinking. Fixed cells were subsequently washed in PBS and suspended in protoplast buffer (50 mM Tris pH 7.4, 50 mM NaCl, 10 mM EDTA, 0.5 M sucrose, EDTA-free protease inhibitor cocktail) supplemented with 20 mg/ml of lysozyme for 2 h at 37 °C. After washing in protoplast buffer pellets were resuspended in buffer L (50 mM HEPES-KOH pH 7.55, 40 mM NaCl, 1 mM EDTA, 1 % Triton X-100, 0.1 % deoxycholate, 0.1 mg/ml RNaseA; EDTA-free protease inhibitor cocktail) and DNA was sheared into fragments of around 800 bp length by sonication using an ultrasonic cell disruptor (Branson Ultrasonics Sonifier™; 20 % amplitude, pulse 0.5 sec on/off, 6 x 20 sec). following removal of cell debris (20000 g, 10 min, 4°C). Aliquots of cell extracts were stored for later use. Dynabeads™ Protein G (Thermo Fisher Scientific) were bound to an α-mCherry antibody (BioVision Inc.) in buffer L for 1.5 h at 4°C, washed in buffer L and subsequently incubated with cell extract for 2 h at 4℃. Thereafter, beads were washed in buffer L, in buffer L5 (50 mM HEPES-KOH pH 7.55, 500 mM NaCl, 1 mM EDTA, 1 % Triton X-100, 0.1 % deoxycholate), in buffer W (10 mM Tris-HCl pH 8, 250 mM LiCl, 0.5 % NP-40, 0.5 % deoxycholate, 1 mM EDTA), and TE buffer (10 mM Tris-HCl pH 8, 1 mM EDTA) consecutively and finally resuspended in TES buffer (10 mM Tris-HCl pH 8, 10 mM EDTA, 1 % SDS). Extract samples were also supplemented with TES buffer and SDS to a final concentration of 1 % SDS; cross-links were reverted at 65℃

overnight. Phenol-chloroform extraction yielded DNA pellets which were further purified using a DNA purification kit (QIAquick®, Qiagen). qPCR was applied in order to confirm protein enrichment at specific chromosomal loci. Immunoprecipitation and extract samples were diluted 1:10 and 1:100 in water, yielding concentrations of approximately 0.2-0.4 ng/ µl.

**ChIP-seq analyses**

For sequencing analyses libraries of ChIP samples were prepared followed by sequencing utilizing an Illumina MiSeq system. Reads were aligned to the *C. glutamicum* ATCC 13032 genome sequence (GeneBankID: BX927147.1), where RES167-specific genome deletions were manually cut using CLC Genomics Workbench. Data of extract and corresponding ChIP sample were each normalized based on read counts and the ratio of the number of reads per 0.5 Kb bin were determined via the Galaxy web platform [14,15].

**Protein purification**

ParB protein production was performed in *E. coli* BL21 pLysS via the pET-16b vector-based system. Cells were grown in LB at 37°C; gene expression was induced adding 1 mM IPTG following growth for 12 h at 18°C. Subsequently, cells were suspended in washing buffer (50 mM Tris-HCl, pH 7.4; 100 mM NaCl; 5 mM; $MgCl_2$; 1 mM dithiothreitol) containing EDTA-free proteinase inhibitor (cOmplete™, Sigma) and DNAseI and lysed using a high-pressure cell homogenizer. Cell debris and membranes were removed by centrifugation at 4°C, 1700 g for 20 min, and 150000 g for 45 min, respectively. Thereupon, batch purifications of His-tagged protein were performed under native conditions using Ni-NTA agarose (Protino®, Macherey-Nagel) according to manufacturer's instruction. In brief, the equilibrated gel was incubated with clarified lysate for 60 min at 4°C under gentle agitation and washed twice in washing buffer containing 80 mM imidazole. Proteins were eluted in three steps using washing buffer with an imidazole concentration of 300 mM, concentrated via Amicon filter units (Merck) and further purified by applying size exclusion chromatography using an ÄKTApurifier system with a Superdex™ 200 gel filtration column (GE Healthcare Life Sciences).

**PALM microscopy**

For sample preparation, *C. glutamicum* cells were harvested in exponential growth phases, washed twice in PBS and fixated in PBS + 3 % formaldehyde solution (36.5-38% in $H_2O$ + 10-15% methanol, Sigma Aldrich) for 30 min at 30°C. Excess formaldehyde was subsequently quenched by adding 10 mM glycine, cells were sedimented at 5000 g for 1 min, resuspended in PBS containing 10 mM glycine and incubated for 5 min at room temperature. This quenching step was repeated three times; cells were finally diluted in buffer containing 50 mM Tris pH 7.4, 50 mM NaCl, 10 mM EDTA and 0.5 M sucrose.

Super-resolution imaging was performed on a Zeiss ELYRA P.1 microscope (laser lines HR diode 50 mW 405 nm and HR DPSS 200 mW 561 nm). Cellular PAmCherry-tagged proteins were detected via an Andor EM-CCD iXon DU 897 camera as described before, using a long pass 570 nm filter (LP570) and an alpha Plan-Apochromat 100x/ 1,46 Oil DIC M27 objective for imaging. Further, 100 nm TetraSpeck microspheres and the implemented drift correction tool served for drift correction; the Z-axis was stabilized via the “definite focus” system. PALM image calculation was performed applying the 2D x/ y Gaussian fit (Zen2 software, Zeiss) using a peak mask size of 9 pixels, where one pixel corresponds to 100 nm and a peak intensity to noise ratio of 6. In order to exclude background and events resulting from the co-emission of co-localizing molecules, events were filtered for photon numbers between 70-350 and point spread function (PSF) width at 1/ e maximum (70-170 nm) were applied. As a last step, events were grouped according to the following parameters: 3 on-frames with 0 off-frames allowed and a search radius of 30 nm.

When imaging strains containing ParB-PAmCherry, four imaging series were taken for each field of view, where each subsequent serie was characterized by a specific 405 nm laser linear gradient intensity range (0.001% to 0.01%, 0.01% to 0.1%, 0.1% to 1% and 1% to 10%). Every other imaging parameter remained the same in between the time series. The frame count for each collection was 10000 frames and converted molecules were imaged using the 561 nm laser at 15 % (transfer mode) for 50 ms at a 200-fold EMCCD gain.

The workflow of protein cluster analysis is illustrated in Figure S6. The field of view in the bright field channel was correct for illumination unevenness by dividing the field of view containing the cells of interest with an empty one (Process - Calculator Plus, Fiji) and enlarged 10 times (bicubic interpolation). The resulting image was thresholded (Image – Adjust - Threshold) with default parameters and converted to a binary mask. A Fiji macro was then run on the binary mask to close the mask holes present within cells and to enlarge the cells mask themselves. Cells that were in contact with each other were separated via water shading. The perimeter coordinates corresponding to masks representing cells lying within the focus were extracted and used to exclude events originating from cells lying outside the focal plane and the background. The clustering structures of events within a cell were identified via the OPTICS algorithm in R [16,17]. Macro- and subclusters of ParB protein were identified by setting the following parameters within the OPTICS algorithm: minimum points = 32, epsilon =3000. The threshold epsilon is 50 for the macroclusters and 35 for the subclusters.

**Chromosome conformation capture libraries**

In the following, a more detailed outline is given of the preparation of *C. glutamicum* chromosome conformation capture libraries. Briefly, cells were grown in 200 ml of BHI medium at 30°C to an $OD_{600}$ of 3 and rediluted to a final concentration of ~1 x $10^7$ cells/ ml. Cells were crosslinked using fresh formaldehyde for 30 minutes at room temperature (3 % final concentration; Sigma Aldrich Formalin 37 %) followed by 30 minutes at 4°C. Formaldehyde was quenched using a final concentration of 0.25 M glycine for 20 minutes at room temperature (RT). Cells were then collected by centrifugation, frozen in dry ice and stored at -80°C until use. Frozen pellets of ~$10^9$ cells were thawed on ice and suspended in a final volume of 1.1 mL 1X TE (pH 8) and transfer in a VK01 Precellys Tube (beads beating). Fixed cells were disrupted using the following program on a precellys apparatus: 9 cycles x [20“ – 3500 rpm; 30“ – pause]. Lysate was transferred to a 1.5 ml tube, SDS 10% was added to the mix to a final concentration of 0.5% and the mix was incubated for 10 minutes at RT. 1 ml of lysate was then transferred in a 5 ml tube containing 4 ml of digestion mix (1X NEB 3 buffer, 1% Triton X-100, and 1000 U MluCI enzyme). DNA was digested for 3 hours at 37°C under shaking. Insoluble fraction was then recovered through centrifugation (16,000 x g – 20 min) and the obtained pellet was resuspended in 1 ml of water and diluted in 15 ml of ligation reaction mix (1X ligation buffer NEB without ATP, 1 mM ATP, 0.1 mg/ ml BSA, 125 Units of T4 DNA ligase 5 U/ ml). Ligation was allowed to proceed for 4 hours at 16° C, followed by incubation overnight at 65°C in presence of 250 mg/ml proteinase K, 0.5 % SDS and 5 mM EDTA. Next morning, DNA was precipitated using 1/10 th volume of 3 M Na-Acetate (pH 5.2) and one volume of iso-propanol. After one hour at -80° C, DNA was pelleted, resuspended in 900 µl 1X TE buffer and extracted with 900 µl phenol-chloroform pH 8.0. DNA was again precipitated using 1/10 th volume of 3 M Na-Acetate (pH 5.2) and 2.5 volume of cold Ethanol. Finally, DNA was resuspended in 100 µl 1X TE buffer supplemented with RNAse and incubated 30 min at 37°C. 3C libraries were then processed as described [18] and paired end sequenced on an Illumina NextSeq apparatus (2 x 35 bp).
